# Retinal development driven by TET-dependent DNA demethylation

**DOI:** 10.64898/2026.04.09.717491

**Authors:** Galina Dvoriantchikova, Michelle Fleishaker, Byron L. Lam, Dmitry Ivanov

**Affiliations:** Bascom Palmer Eye Institute, Department of Ophthalmology, University of Miami Miller School of Medicine, Miami, FL 33136, USA; Illinois Eye and Ear Infirmary, Department of Ophthalmology, University of Illinois College of Medicine, Chicago, IL, 60612, USA; Department of Microbiology and Immunology, University of Miami Miller School of Medicine, Miami, FL, 33136, USA

**Keywords:** TET dioxygenases, DNA demethylation, retinal development, RPC proliferation, photoreceptors

## Abstract

Inactivation of the TET-dependent DNA demethylation pathway in retinal progenitor cells (RPCs) disrupts retinal development and leads to blindness. In this study, we demonstrated that this is due to the global control exerted by this pathway at all stages of retinal development. TET-deficient RPCs exhibit characteristics of both early and late progenitors, a factor that most likely contributes to the disrupted cellular composition of the retina, wherein the cone population expands significantly at the expense of all other cell types. The differentiation of TET-deficient RPCs also results in the formation of populations of abnormal retinal cell types, a phenomenon particularly evident in the development and function of rod and cone photoreceptors. This global control of a single epigenetic pathway is explained by the high level of methylation in RPCs of many genes critical for retinal development. These genes must be demethylated by TET enzymes to be activated in developing retina.

## Introduction

The retina consists of several types of neurons whose function is to convert light into electrical signals (rod and cone photoreceptors) and then transmit them to the brain (bipolar, horizontal, amacrine, and ganglion cells [RGCs] are responsible for this) to create the sensation of vision^1–4^. Impaired development of these neurons can significantly affect retinal function and even lead to blindness^5^. All retinal neurons and Müller glia differentiate from retinal progenitor cells (RPCs), which are classified—based on their properties—as either early or late. Early RPCs differentiate into early-born neurons (cones, RGCs, horizontal cells, and a subpopulation of amacrine cells) and late RPCs^1–4^. Late RPCs differentiate into late-born neurons (rods, bipolar cells, and a subpopulation of amacrine cells) and Müller glia^1–4^. A multitude of signaling cascades govern the differentiation of proliferating RPCs into various types of non-proliferative retinal cells^1–4,6,7^. There is growing evidence that epigenetic mechanisms, through their ability to control the expression of many genes in these signaling cascades, play a critical role in the development of the retinal cell types^8,9^. In this study, we examine the role of DNA methylation, as an epigenetic mark, and the process of DNA demethylation in retinal development.

The development of a tissue is accompanied by changes in the patterns of methylated cytosines in the DNA of its constituent cell types^9–11^. These methylation patterns maintain optimal patterns of gene expression necessary for normal cell activity in tissue^9–11^. There are two families of enzymes that are responsible for the formation of DNA methylation patterns in cells: enzymes belonging to the DNA methyltransferase (DNMT) family catalyze the transfer of a methyl group to the cytosine nucleotide, while the Ten-Eleven Translocation (TET) family promotes DNA demethylation^9–11^. Changes in DNA methylation patterns are critical to retinal development since many genes essential for the development and function of the various retinal cell types are hypermethylated in RPCs from which these cell types originate^12–16^. Active demethylation of these genes is carried out by TET enzymes, which explains their crucial role in the development of the retina: genetic ablation of TET enzymes in RPCs impairs retinal development and leads to blindness^17–19^. However, the underlying mechanisms, controlled by TET enzymes during retinal development, the disruption of which leads to pathology, remain unknown.

To address the knowledge gaps, we monitored changes in the structure and size of the developing TET-deficient and control retinas and studied RPC proliferation in them. We linked observed changes to the level of gene expression at different stages of retinal development and how all these changes affected the cellular composition of the retina. We also studied how DNA methylation controls gene expression in the developing retina. Our results indicate that DNA methylation (by directly and indirectly inhibiting the expression of multiple critical genes) and the TET-dependent DNA demethylation pathway exert global control over retinal development, ranging from the RPC stage to the formation of functional retinal cell types.

## Results

### TET-deficient developing retinas are characterized by slow growth, a large number of non-dividing cells, and a small number of RPCs that continue to divide at the very latest stages

We have previously shown that TET-deficient mature retinas differ significantly in structure from control retinas^17^. To track the emergence and manifestation of these changes, we used developing TET-deficient experimental (Chx10TET) retinas and control (TET) retinas collected from embryonic day 16 (E16), postnatal day 0 (P0), P4, P7, and P10 Chx10TET and TET mice. We found that the size of the experimental retinas was smaller than that of the controls at all time points examined (**Fig. 1A**). We then studied RPC proliferation in developing retinas based on EdU incorporation into DNA (**Fig. 1B**). In the embryonic E16 control retinas, EdU incorporation was relatively higher than in TET-deficient retinas (**Fig. 1C, 1D**). After the mice were born, no significant difference was observed until P7. The 7th postnatal day (P7) revealed a significant effect of TET enzymes on RPC proliferation in the developing retinas: RPC proliferation can be detected in the center, middle, and in large quantities at the periphery of P7 TET-deficient (Chx10TET) retinas, while in P7 control (TET) retinas cell proliferation can be partially detected only at the periphery (**Fig. 1C, 1D, Supplementary Fig. S1**). Individual dividing RPCs could still be detected throughout the retina on the 10th postnatal day (P10); however, their number was insufficient to establish a difference between the experimental and control groups (**Fig. 1C, 1D**). Thus, at a late stage of development, many undifferentiated proliferating RPCs are still present in TET-deficient retinas. Analysis of the thickness of the retinal layers revealed that the inner neuroblastic layer (INbL), outer neuroblastic layer (ONbL), and outer nuclear layer (ONL) were relatively thinner in TET-deficient (Chx10TET) retinas (**Fig. 1C, 1E, 1F, Supplementary Fig. S2**). This difference was more than compensated for by the enormous thickness of the ganglion cell layer (GCL) in TET-deficient retinas (**Fig. 1C, 1F, Supplementary Fig. S2**). The thickness of the inner plexiform layer (IPL) and outer plexiform layer (OPL) remained lower in Chx10TET retinas at all time points examined, suggesting issues with the formation of synaptic contacts (**Fig. 1C, 1F, Supplementary Fig. S2**). The thickness of the inner nuclear layer (INL) was approximately the same in experimental and control retinas on the 10th day of postnatal development. The low thickness of the ONbL, which contains RPCs, and the enormous thickness of the GCL, which contains non-dividing retinal cells, suggest that TET-deficient RPCs divide predominantly asymmetrically (one RPC and one non-dividing retinal precursor), which is characteristic of late RPCs. The predominantly asymmetric division of RPCs may also explain the slow increase in size of TET-deficient retinas.

**Figure 1.**
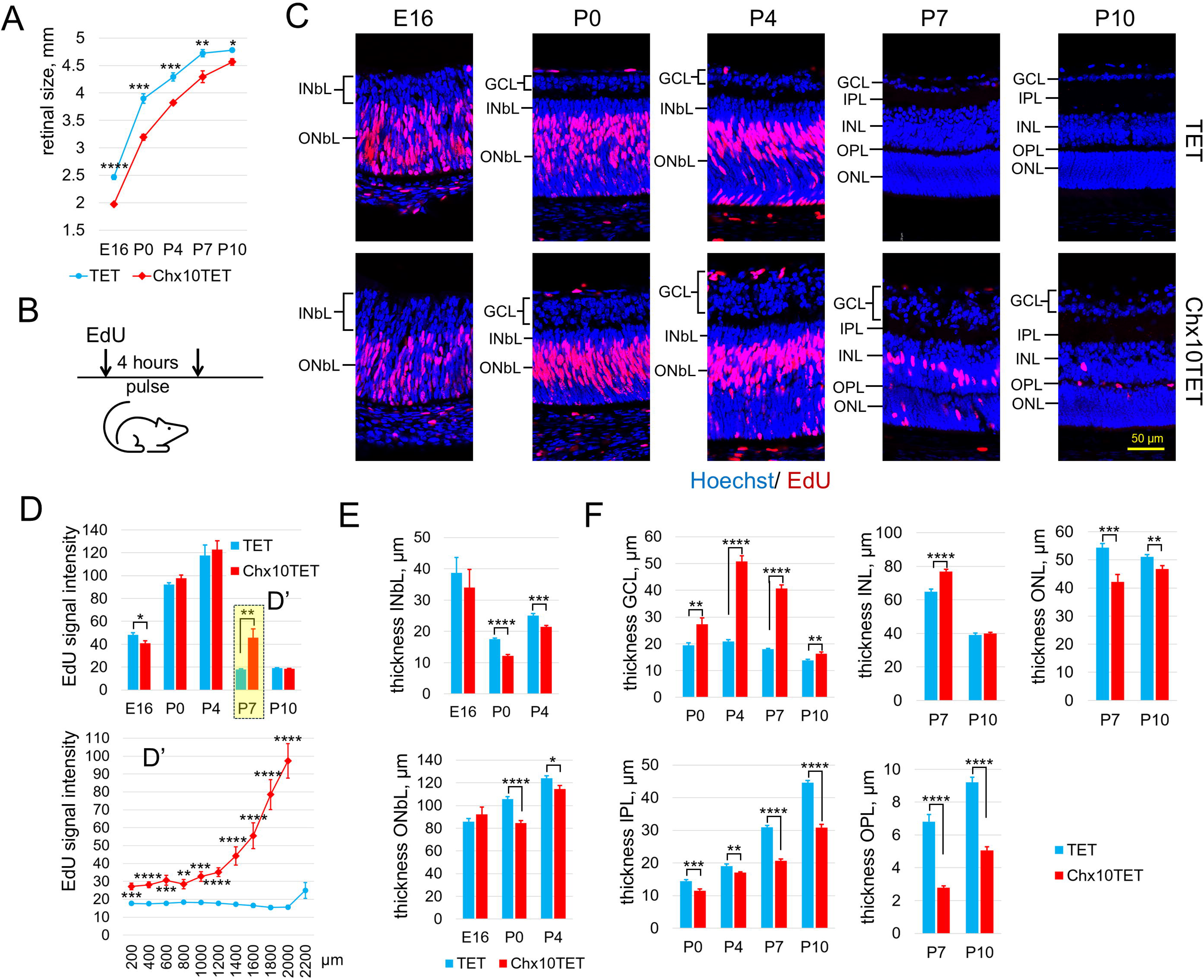
Predominantly asymmetric and sustained division of RPCs likely accounts for the phenotype of TET-deficient developing retinas. **A)** Genetic ablation of the TET family in RPCs affects the growth of the developing retina (n = 5 mice per group, per age; ****P value < 0.0001, ***P value <0.001, *P value < 0.05). **B)** The EdU solution was injected intraperitoneally into mice. After 4 hours (pulse), the retinas were collected and used for analysis. **C)** Representative confocal images of developing TET-deficient (Chx10TET) and control (TET) retinas treated with EdU. **(D-F)** The results of measurements of mean fluorescent EdU signal intensity (**D**) and layer thickness (**E, F**) indicate impaired development of the TET-deficient retina (n ≥ 5 mice per group, per age; INbL - inner neuroblastic layer, ONbL - outer neuroblastic layer, GCL – ganglion cell layer, INL – inner nuclear layer, ONL – outer nuclear layer, IPL – inner plexiform layer, OPL – outer plexiform layer).

### TET-deficient RPCs exhibit a high rate of proliferation

During retinal development, the population of rapidly dividing early RPCs is replaced by a population of slowly dividing late RPCs, most of which differentiate by the 7th postnatal day^20^. The presence of numerous dividing RPCs in TET-deficient retinas on the 7th postnatal day suggests a slower transition of RPCs from the rapid-division phase to the slow-division phase. To test this hypothesis, we used a short pulse (30 min) to incorporate EdU into the DNA of dividing RPCs (**Fig. 2A**). With this approach, only rapidly dividing RPCs would be capable of incorporating EdU into their DNA. In our experiment, we utilized P4 experimental and control mice in which we had previously observed no differences in EdU incorporation when using a standard pulse (**Fig. 1C, 1D**). The results of our analysis indicate a high abundance of rapidly dividing RPCs in the center, middle, and periphery of TET-deficient retinas (**Fig. 2B, 2C**). Conversely, in the control retinas, rapidly dividing RPCs were predominantly concentrated in the periphery (**Fig. 2B, 2C**). Since a high proliferation rate is characteristic of early RPCs, our data suggest that, even at a late stage of retinal development, TET-deficient RPCs possess the properties of early RPCs.

**Figure 2.**
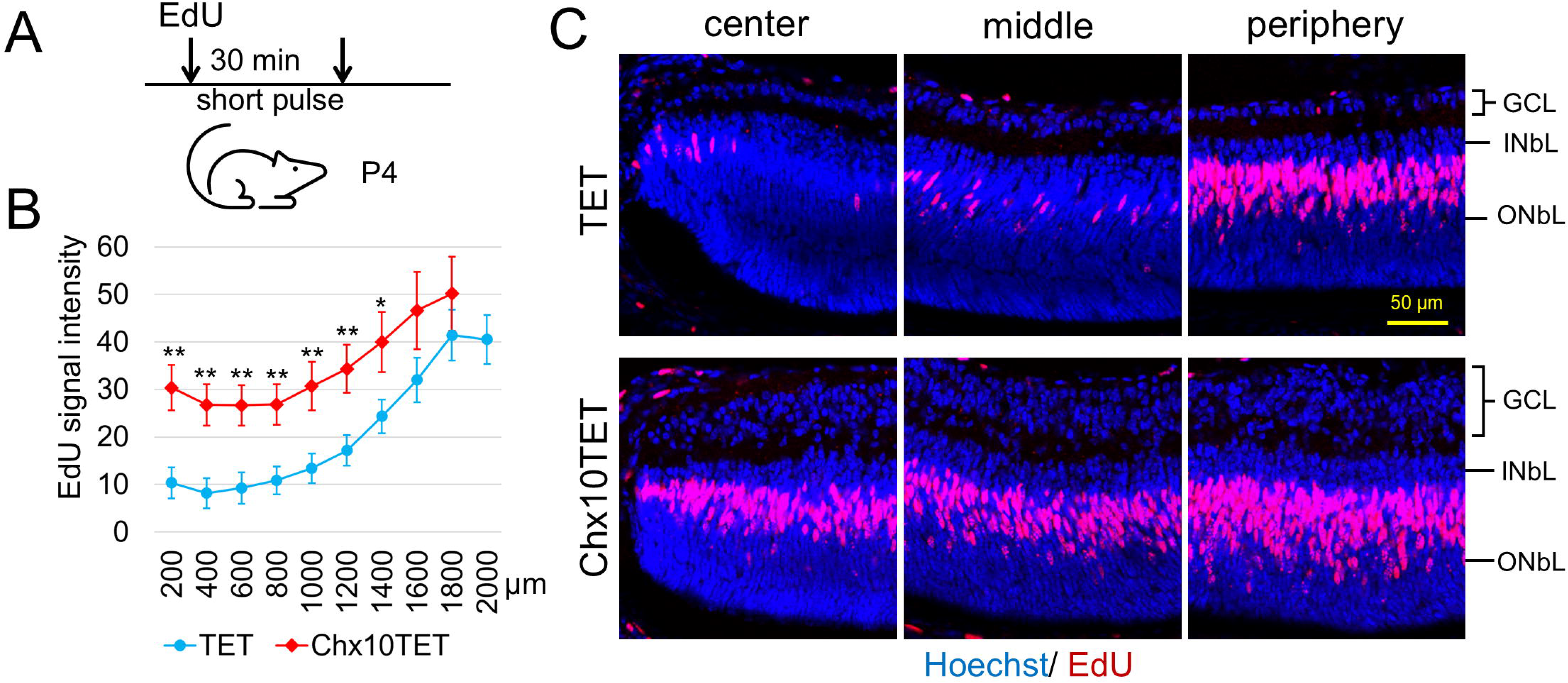
TET-deficient RPCs possess the properties of early RPCs. **A)** P4 experimental and control mice were treated with EdU for a brief period (short pulse) to ensure that it would be incorporated into the DNA of only rapidly dividing RPCs. **(B, C)** The results of measurements of EdU signal intensity (**B**) and representative confocal images (**C**) indicate that the experimental (Chx10TET) RPCs divide faster than the control (TET) RPCs (n ≥ 5 mice per group, **P value <0.01, *P value < 0.05).

### TET enzymes control the expression of genes important for RPCs and for various retinal cell types

To elucidate the molecular mechanism responsible for the observed changes, we studied gene expression in developing Chx10TET vs. TET retinas using E16, P0, and P7 mice and bulk RNA-seq analysis. We used P14 and P30 Chx10TET vs. TET bulk RNA-seq data to compare changes occurring in developing and mature retinas. Since the entire mouse retina consists of RPCs on the 11th day of embryonic development (E11), we also used wild-type E11 retinas to obtain the high-throughput gene expression profile of RPCs. Our results indicate that the gene expression profiles of E16 and P0 retinas are closer to those of RPCs, whereas the gene expression profiles of P7 retinas are closer to those of mature retinas (P14 and P30; **Fig. 3A**). Differences between the gene expression profiles in Chx10TET vs. TET retinas become more distinct with age, reaching a peak on the 7th day after birth (P7; **Fig. 3B, Supplementary Datafile S1**). In mature retinas (P14 and P30), the gene expression profiles differ not by age, but by genotype (**Fig. 3A, 3B**). One of the important observations is the increased expression of *Zic1*, a transcription factor that sustains the proliferation activities of RPCs (**Fig. 3C, Supplementary Datafile S1**)^21^. Consistent with our findings of increased RPC proliferation in P7 Chx10TET vs. TET retinas, we found increased expression of many genes involved in the cell cycle (**Fig. 3D, Supplementary Datafile S1**). It is important to note that, starting from birth, the expression of many genes necessary for the development and function of rod photoreceptors was significantly reduced in the experimental TET-deficient retinas compared to TET controls (**Fig. 3D, 3E, Supplementary Datafile S1**). On the 7th day (P7), we detected reduced expression of a substantial number of genes required for the development and function of bipolar cells (**Fig. 3E, Supplementary Datafile S1**). Bipolar cells are the very last retinal neurons to differentiate from RPCs^3^. Thus, the observed changes are expected to impact their development in TET-deficient retinas. We also detected reduced expression of genes essential for the development and function of other retinal cell types; however, the number of such genes was significantly smaller than the number of genes required for the development and function of photoreceptors and bipolar cells (**Supplementary Datafile S1**). Thus, TET enzymes influence retinal development at all levels—ranging from RPCs to the development of individual cell types.

**Figure 3.**
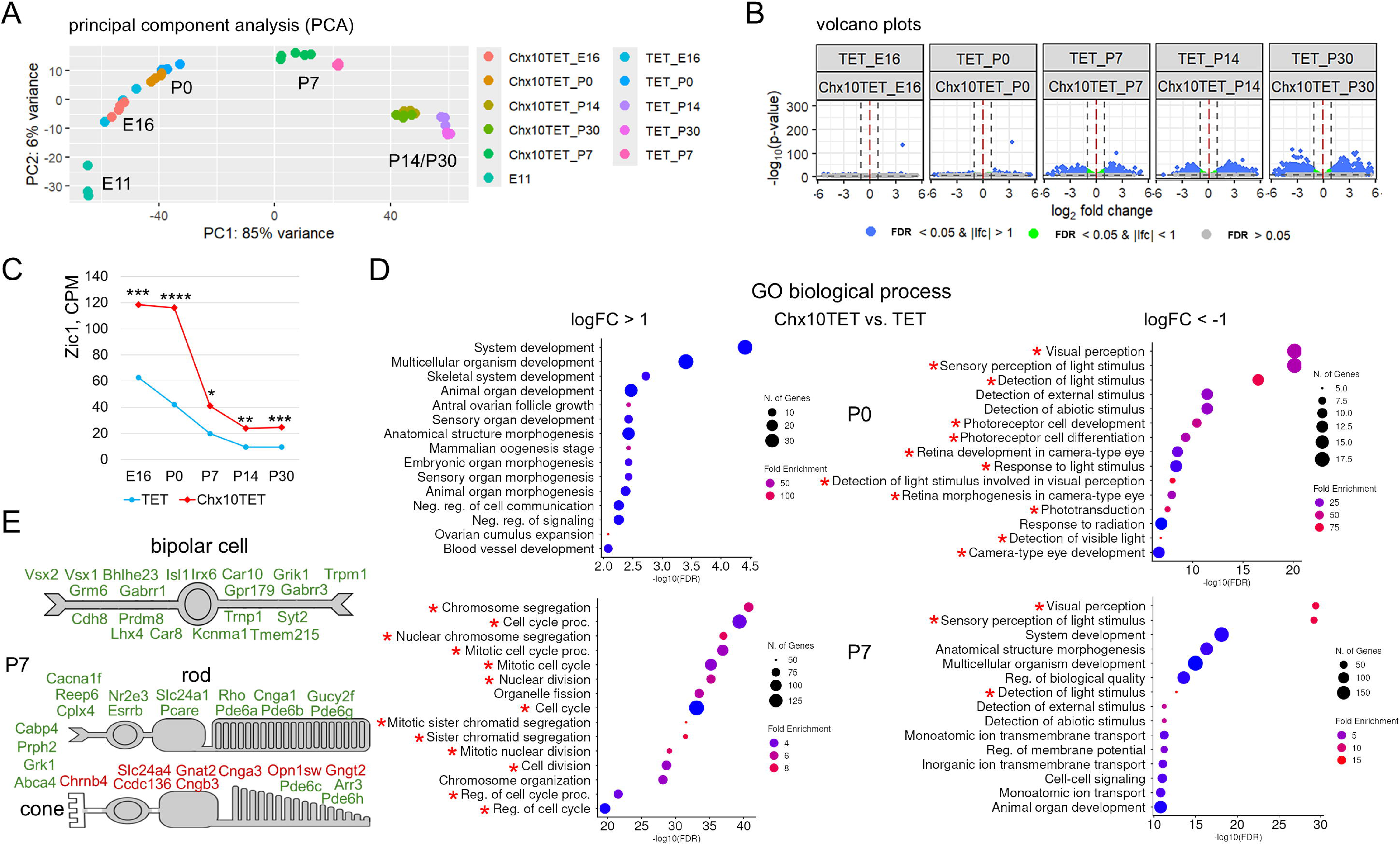
Inactivation of TET enzymes in RPCs significantly affects retinal gene expression during development. **(A, B)** The principal component analysis (PCA, **A**), and volcano plots (**B**) indicate significant differences in gene expression between developing TET-deficient (Chx10TET) and control (TET) retinas. **C)** The dynamics of *Zic1* expression during the development of Chx10TET and TET retinas may affect the expression of genes important for cell proliferation (****P value < 0.0001, ***P value <0.001, **P value < 0.01, *P value < 0.05). **D)** Genes with increased (logFC >1) or decreased (logFC < −1) expression more than twofold in Chx10TET vs. TET retinas were used in the GO biological process analysis. **E)** The panel shows retinal cell types and examples of genes important for their development and function whose expression changed twofold or more. Genes with decreased expression in Chx10TET vs. TET retinas are highlighted in green. Genes with increased expression in Chx10TET vs. TET retinas are highlighted in red.

### The TET-dependent DNA demethylation pathway controls the proper cellular composition of the retina

The changes we observe in developing TET-deficient retinas at morphological and molecular levels should be reflected in the cellular composition of mature retinas. To evaluate the changes in the cellular composition, we used P14 TET and Chx10TET retinas and single-nucleus RNA-seq (snRNA-seq). The experiment was repeated two times on TET and Chx10TET retinas using the 10X genomics platform. In each experiment, we obtained gene expression data from more than 10,000 single cell nuclei (more than 50,000 cells in total). The bioinformatics analysis of our snRNA-seq data obtained on control TET retinas enabled classification of all cells into 31 clusters (**Fig. 4A**). We manually identified which cluster corresponded to which retinal cell type using expression data of known cell markers (**Fig. 4A**, **Supplementary Fig. S3**). Two clusters, 13 and 19, contained markers of glycinergic and nGnG amacrine cells (ACs) and, therefore, we designated them as glycinergic/nGnG ACs (**Fig. 4A**). Cluster 1 contained cells expressing rod markers. However, the expression of these markers and the percentage of cells in the cluster that expressed these markers were lower than in the cluster 0 that represents Rods (**Fig. 4A, Supplementary Datafile S2**). For this reason, we have designated this cluster as Rod-like. The position of cluster 1 and the level of expression of rod markers suggest that the cells in it have not yet completed their differentiation into rod photoreceptors (**Fig. 4B, B’**). The same bioinformatics analysis of the snRNA-seq data obtained on TET-deficient Chx10TET retinas revealed only 26 clusters (**Fig. 4C**, **Supplementary Fig. S4**). While the number of clusters identified was smaller in Chx10TET vs. TET retinas, we found several clusters with cells expressing Müller glia (MG) markers (**Fig. 4D**). We designated cluster 14 with high expression of these markers as MG, while the remaining two as MG-like (**Fig. 4C, 4D, Supplementary Datafile S3**). We also detected clusters (1 and 2) whose cells express only photoreceptor (rod or cone) markers but significantly weaker than clusters 0 (Rods) and 3 (Cones).

**Figure 4.**
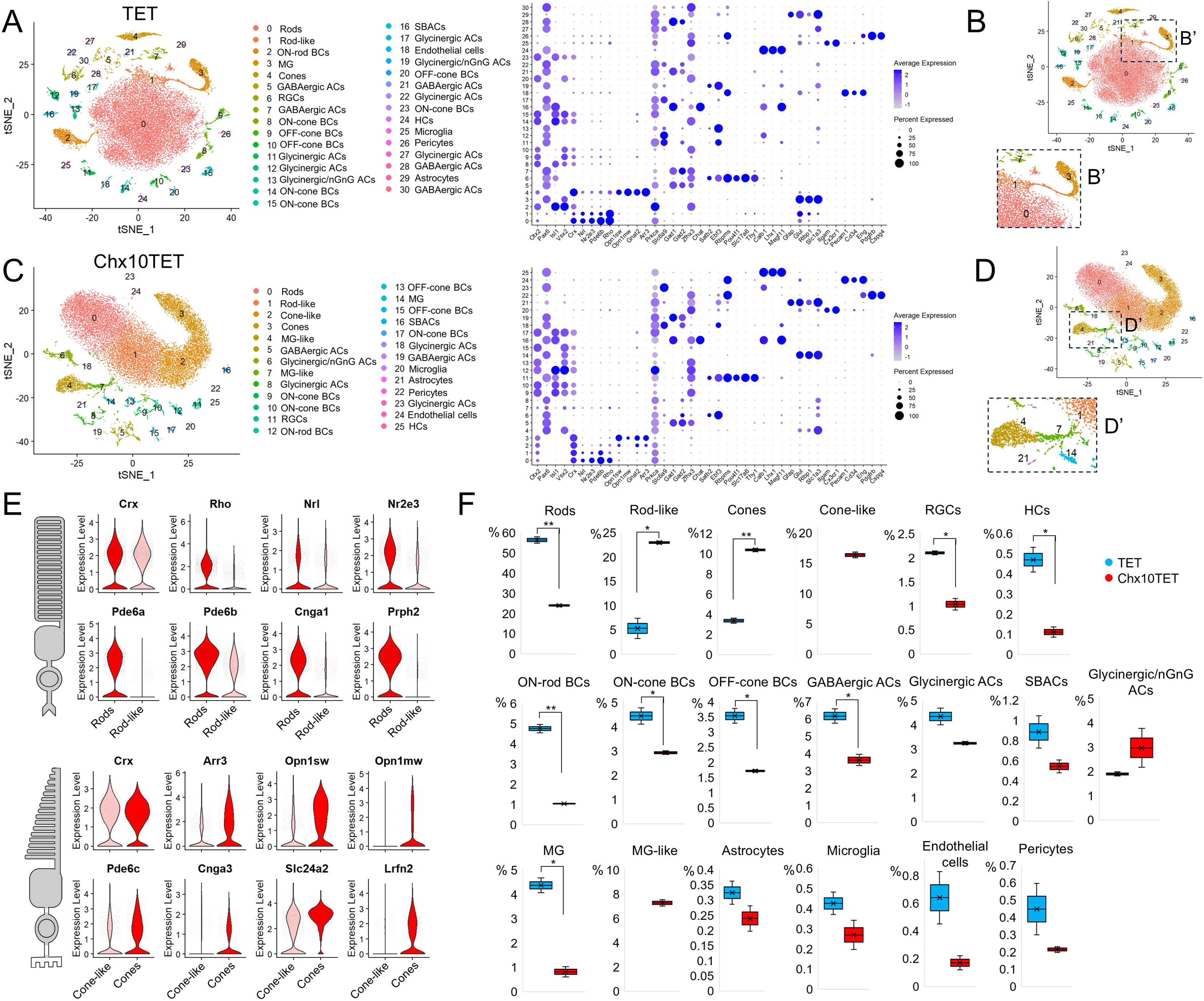
TET-deficient retinas differ significantly from control retinas in their cellular composition and complexity. **A)** Non-linear dimensional reduction t-distributed Stochastic Neighbor Embedding (tSNE) technique was used to visualize and explore identified single cell clusters in control (TET) retinas. Each cluster was associated with a specific cell type using known markers. Eight markers were particularly helpful in identifying retinal cell types: Otx2, which is a marker of cone, rod, and bipolar cells (BCs); Pax6 - a marker of amacrine cells (ACs), horizontal cells (HCs), retinal ganglion cells (RGCs), and Muller glia (MG); Isl1 – a marker of ON BCs, RGCs, and starburst amacrine cells (SBACs); Vsx2 – a marker of BCs and MG; Crx – a marker of cone and rod photoreceptors, Gad1 and Gad2 – markers of GABAergic ACs, and Slc6a9 – a marker of glycinergic ACs. **B)** While all clusters correspond to mature cell types in the control retina, cluster 1 likely contains rods that have not yet completed their development. **(C-E)** The smaller number of clusters (**C**) and the existence of similar clusters with different levels of marker activity and presence (**D, E**) indicate less complexity and underdevelopment of the TET-deficient (Chx10TET) retina compared to the control retina. **F)** The number of cells in each cluster was used to calculate the percentage of the identified cell types in Chx10TET and TET retinas. (*P-value < 0.05, **P-value < 0.01)

Therefore, we have designated them as Rod-like or Cone-like (**Fig. 4C, 4E, Supplementary Datafile S4**). After assigning clusters to retinal cell types, we further determined the percentage of cells in each cluster relative to the total number of cells and compared these percentages between Chx10TET vs. TET retinas (**Fig. 4F**). We found that the number of Rods in control retinas significantly exceeds the number of Rods in TET-deficient retinas (TET 56±2% vs. Chx10TET 24±1%, P-value = 0.003). However, the numbers are opposite for Rod-like cells (TET 5±2% vs. Chx10TET 22.9±0.3%, P-value = 0.01). The number of Cones in Chx10TET retinas significantly exceeds the number of Cones in control TET retinas (TET 3.4±0.3% vs. Chx10TET 10.4±0.2%, P-value = 0.002). Chx10TET retinas also contain Cone-like cells (16.4±0.5%), and the sum of the percentages of TET-deficient Cones and Cone-like cells is 9 times greater than the percentage of Cones in control TET retinas. Apart from photoreceptors, the percentage of many other types of retinal neurons is lower in TET-deficient retinas compared to controls (**Fig. 4F**). The percentage of bipolar cells (BCs) decreases the most (TET vs. Chx10TET: ON-rod BCs, 4.8±0.2% vs. 1.04±0.03%, P-value = 0.003; ON-cone BCs, 4.4±0.3% vs. 2.9±0.1%, P-value = 0.048; OFF-cone BCs, 3.5±0.2% vs. 1.71±0.04%, P-value = 0.02). It should also be noted that while the percentage of MG is lower in Chx10TET vs. TET retinas (0.8±0.2% vs. 4.4±0.3%, P-value = 0.01), the percentage of TET-deficient MG-like cells (7.3±0.3%) exceeds the number of control MG. To compare gene expression in Chx10TET vs. TET retinas, we integrated our snRNA-seq data (**Fig. 5A**). We found 33 clusters that included features of Chx10TET and TET retinas (**Fig. 5A, 4A, 4C**, **Supplementary Fig. S5**). It should be noted that Rod-like, Cone-like and MG-like clusters contain mostly cells from Chx10TET retinas (**Fig. 5B**). We then estimated the percentage of cells in each cluster relative to the total number of cells, taking into account which retinas they came from (TET vs. Chx10TET). The results of comparing the percentages of cells in TET vs. Chx10TET retinas after the integration of our snRNA-seq data turned out to be almost indistinguishable from those obtained before integration (**Supplementary Fig. S6**). At the same time, our data indicate that in terms of gene expression, TET-deficient Rods, Rod-like cells, Cones, and Cone-like cells are very different from those in control retinas (**Fig. 5C, 5D, Supplementary Datafile S5**). While a photoreceptor marker Crx is equally highly expressed in Chx10TET and TET cells comprising clusters 0, 1, 2, and 4, the expression of many other photoreceptor markers is significantly reduced in TET-deficient photoreceptors (**Fig. 5C-5F, Supplementary Fig. S7**). The significant decrease in M-opsin (Opn1mw) expression compared to S-opsin (Opn1sw) expression in Chx10TET retinas suggests that the majority of Cones and Cone-like cells in TET-deficient retinas are S-Сones or S-Сone-like (**Fig. 5D**). We found a significant decrease in the expression of genes responsible for phototransduction, the formation of outer segments and synapses of photoreceptors (**Fig. 5E, 5F, Supplementary Fig. S7, Supplementary Datafile S5**). This decrease in expression prevents the normal development of outer segments and synaptic patterns in TET-deficient retinas (**Fig. 5F**). We also found reduced expression of genes responsible for synaptogenesis in TET-deficient retinal cells that make up clusters corresponding to BCs, HCs, and ACs (**Fig. 5F, Supplementary Fig. S7, Supplementary Datafile S6**). Decreased expression of some genes responsible for lamination in the retina may partly explain the disturbances in the lamination of the IPL of the Chx10TET retina that we observed (cluster 17, **Fig. 5G**).

**Figure 5.**
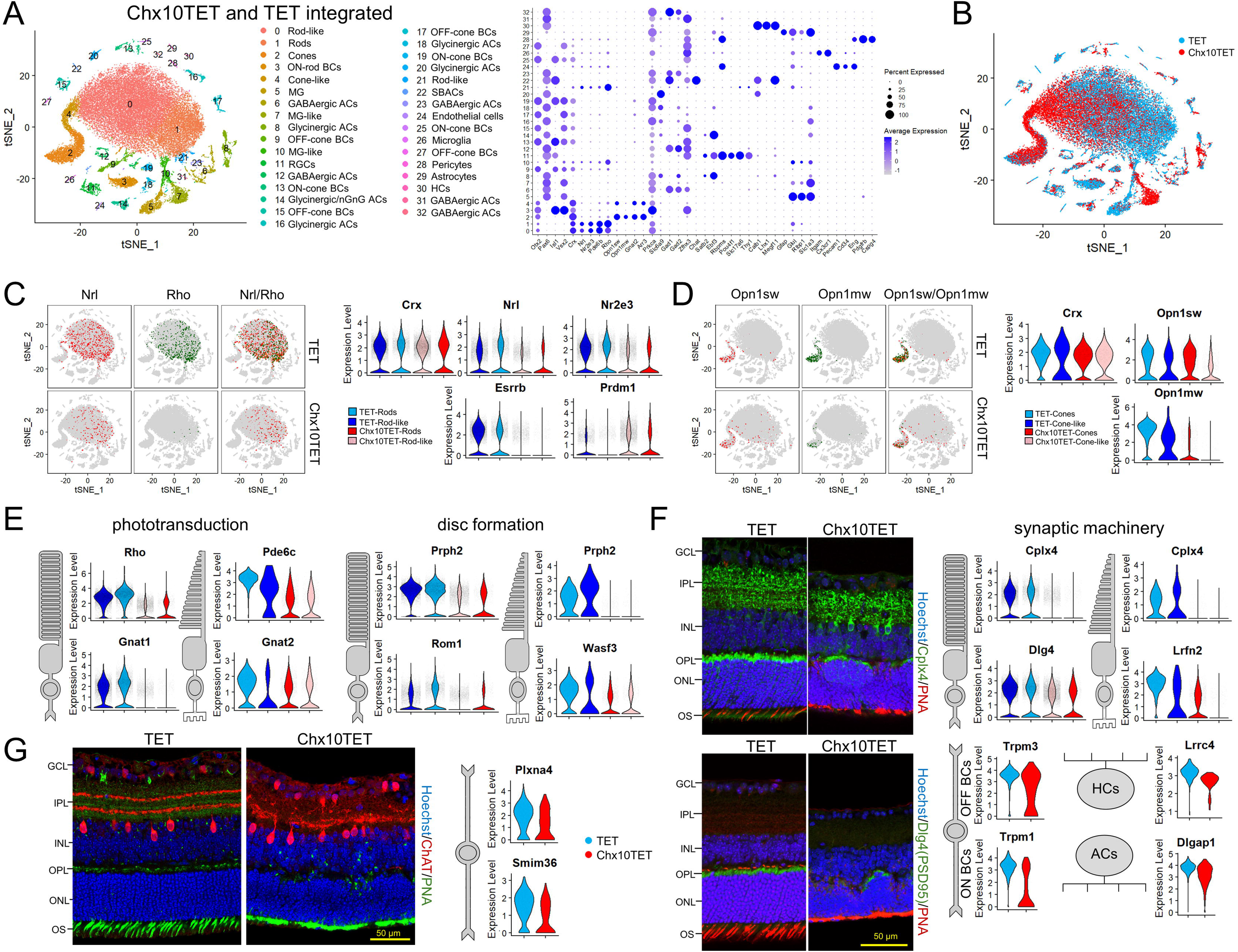
The increase in the number of S-cones in TET-deficient retinas occurs at the expense of all retinal cell types. **A)** The panel shows the results of the snRNA-seq data integration. The cell type corresponding to each cluster was determined based on the expression of specific markers. **B)** Integration of the snRNA-seq data indicates that Rod-like, Cone-like, and MG-like cells are predominantly contained in clusters where TET-deficient cells (Chx10TET) predominate. **C)** Low but detectable expression of some rod specification markers in cells of the TET-deficient retina may result in the total percentage of Rods and Rod-like cells differing slightly between the Chx10TET and TET retinas when the snRNA-seq data are clustered. **D)** The peculiarity of TET-deficient retinas is not only that they have a higher content of Cones and Cone-like cells, but also that these photoreceptors are mainly S-Cones and S-Cone-like cells. **(E, F)** Reduced expression of genes responsible for phototransduction (**E**), outer segment formation (**E**), and synaptogenesis (**F**) results in the abnormalities seen in the confocal images (**F**) of Chx10TET vs. TET retinas. **G)** Altered gene expression in TET-deficient bipolar cells (BCs) may result in abnormal lamination of the inner plexiform layer (IPL).

### Photoreceptor genes constitute a significant proportion of highly methylated genes in RPCs that must be demethylated to be activated in the mature retina

Bulk RNA-seq and snRNA-seq data indicate that the TET dependent DNA demethylation pathway increases the expression of genes essential for the development and function of many retinal cell types. A direct mechanism for such increased expression would be demethylation of the transcription start sites (TSSs) of these genes during differentiation of RPCs into retinal cell types (**Fig. 6A**). Consequently, the TSSs of these genes should be located within highly methylated regions of RPC DNA. To find the TSSs, we used whole genome bisulfite sequencing (WGBS) data obtained from embryonic 11 (E11) retinas. We also used our previously published WGBS data obtained on Chx10TET and TET retinas^17^. Average methylation levels of DNA regions were obtained using the methylKit Bioconductor package. We selected only those genes whose TSSs (±500 bp) are located within highly methylated (≥75%) regions of the RPC DNA. The gene expression level is required to be reduced in Chx10TET vs. TET retinas and to exceed a threshold of 10 CPM. We found that most of the highly methylated genes are required for photoreceptor development and function, and only a small number are required for the development and function of other retinal cell types (**Fig. 6B**, **Supplementary Datafile S7**). A significant decrease in methylation in TET retinas was found only for photoreceptor genes, which is understandable since photoreceptors make up 70% of all retinal cells. Methylation of the TSSs of these genes remained high in Chx10TET retinas (**Fig. 6B**). The approach we have described is, however, more qualitative in nature than quantitative. To overcome the limitations of this approach, we used the DMRseq Bioconductor package to search for differentially methylated regions (DMRs) when comparing the methylomes of E11, Chx10TET, and TET retinas. We selected only those genes whose TSSs (±500 bp) are located in DMRs with reduced methylation in TET vs. E11 and TET vs. Chx10TET (the gene expression level is required to be reduced in Chx10TET vs. TET retinas; **Fig. 6C**, **Supplementary Datafile S7**). Consistent with our previous findings, the majority of the genes identified through this alternative approach proved to be essential for the development and function of photoreceptors (**Fig. 6C, 6D**, **Supplementary Datafile S7, S8**). By combining the results obtained from both approaches, we compiled a list of genes that are highly methylated in RPCs and must undergo demethylation to become transcriptionally active within the retina (**Fig. 6E, Supplementary Datafile S7**). These genes controlled by the TET-dependent DNA demethylation pathway play a crucial role in the development and function of the retina, and particularly of photoreceptors (**Fig. 6E**). It is important to note that a number of these genes are essential for the development and function of many retinal cell types, which likely explains the pathological phenotype observed in TET-deficient retinas. Two genes, Lrrtm2 and Sncg, however, stand apart; they are likely critical solely for the development and function of RGCs^22,23^. The expression of these genes was already reduced in embryonic (E16) TET-deficient retinas (**Supplementary Datafile S1**). Our results suggest that this reduction was likely caused by high levels of methylation of the promoters of these genes (**Fig. 6B, 6E**, **Supplementary Datafile S7, S8**).

**Figure 6.**
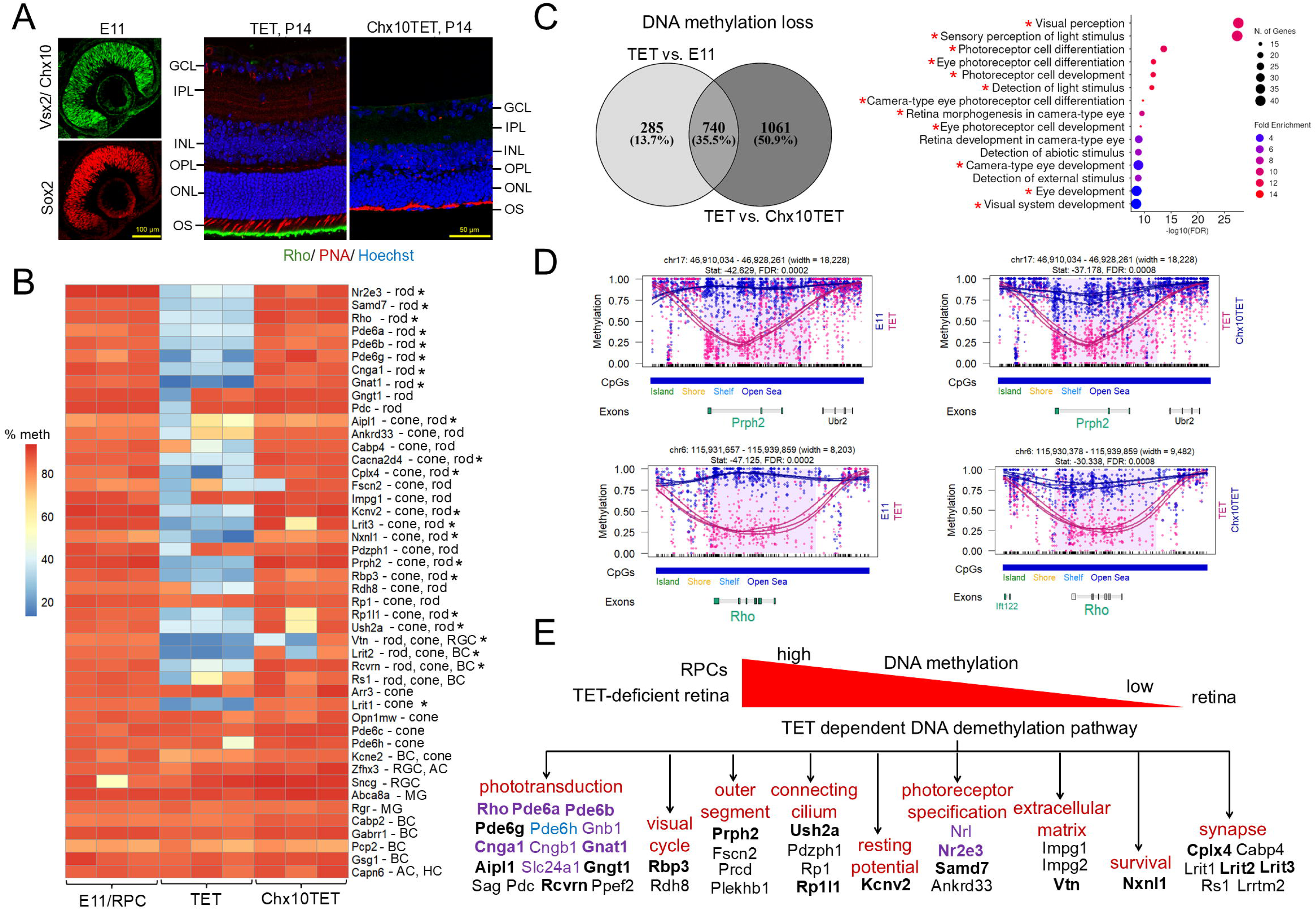
Among the hypermethylated genes in RPCs that are essential for retinal development and function, photoreceptor genes predominate. **A)** Confocal images of Sox2- and Vsx2 (Chx10) -labeled RPCs in embryonic retinas indicate that by day 11 (E11) of development, virtually all cells in the retina are RPCs. Genetic ablation of TET enzymes in RPCs (Chx10TET mice) results in retinas with abnormal morphology and lacking rod (Rho as a marker) and cone (PNA as a marker) outer segments. (P14 – postnatal-day 14). **B)** The heatmap shows the average level of gene TSS±500 bp methylation in embryonic retinas (E11) containing only RPCs, and in matured experimental (Chx10TET) and control (TET) retinas. **C)** A Venn diagram enabled the identification of genes whose TSS±500 bp methylation levels are low in control TET retinas but high in TET-deficient Chx10TET retinas and RPCs. These genes were used in the GO biological process analysis. **D)** Examples of DMRs identified using the DMRseq package are shown in the panel. **E)** The panel shows the genes whose methylation and expression are directly controlled by the TET dependent DNA demethylation pathway during retinal development. Rod-specific genes are highlighted in purple. Cone-specific genes are highlighted in blue. Genes in bold were confirmed using both the methylKit and DMRseq packages.

### The process of DNA demethylation differs in rods and cones

Since photoreceptor genes are predominantly hypermethylated in RPCs, we investigated their demethylation process using publicly available WGBS data derived from isolated rods and cones^24–26^. This approach allowed us to eliminate the background noise generated by other cell types when analyzing WGBS data obtained from whole retinas. As described previously, we utilized the methylKit Bioconductor package to identify genomic regions exhibiting distinct average levels of DNA methylation within the same genome. By analyzing WGBS data derived from DNA isolated specifically from rods and cones, we were able to enhance the statistical significance of the observed demethylation events within the TSSs±500 bp of the photoreceptor genes (**Fig. 7A**, **Supplementary Datafile S9**). However, this approach does not entirely mitigate the substantial variability arising from differing levels of genomic coverage across various samples. This issue is effectively addressed by the DMRseq Bioconductor package, which was specifically designed to resolve such technical challenges. We found that the TSSs (±500 bp) of all photoreceptor genes located in DMRs when comparing the methylomes of RPCs and retinas are also located in DMRs when comparing the methylomes of RPCs, rods, and cones (**Supplementary Datafile S9, S10**). However, one notable distinction emerged: while a significant number of genes underwent demethylation in both rods and cones, two distinct groups of genes were identified that underwent demethylation exclusively in either rods or cones (**Fig. 7B**, **Supplementary Datafile S9, S10**). The number of such genes was relatively small in cones, whereas in rods, we identified 13 genes exhibiting this specific pattern (**Fig. 7C**, **Supplementary Datafile S9, S10**). Thus, during the differentiation of RPCs, the process of DNA demethylation in developing rods and cones exhibits distinct cell-type-specific characteristics (**Fig. 7C**). Given the large number of critical photoreceptor genes that undergo demethylation during photoreceptor differentiation, we anticipated that this process would be the dominant epigenetic event in rods and cones. To our surprise, during the development of rods and cones, a greater number of gene TSSs (±500 bp) undergo methylation than undergo demethylation (**Fig. 7D**). However, the number of gene TSSs (±500 bp) that undergo demethylation predominates in rods compared to cones (**Fig. 7D**).

**Figure 7.**
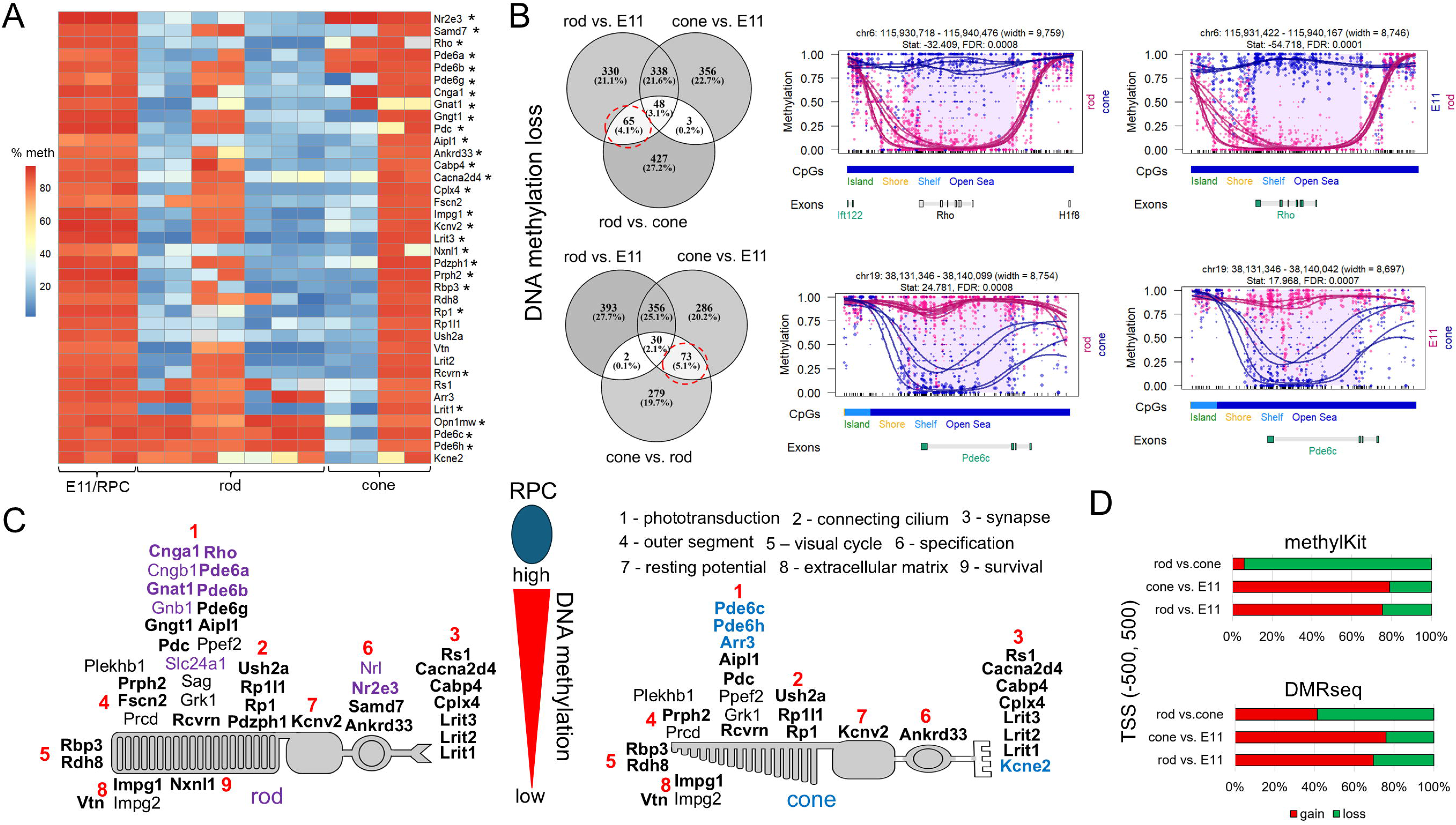
The methylation pattern of RPCs during their differentiation into rods and cones transitions into two distinct methylation patterns. **A)** Average methylation levels of genomic regions identified in the segmentation analysis (methylKit) and photoreceptor genes (hypermethylated in RPCs) were used to generate the heatmap. **B)** Venn diagrams were used to identify genes whose TSSs±500 bp are located in regions (DMRs) with decreased methylation levels specific to rods or cones (these are circled). Examples of DMRs are shown in the panel. **C)** The panel shows genes whose TSSs±500 bp methylation is reduced when RPCs differentiate into rods only (highlighted in purple), cones only (blue), or rods and cones (black). These genes are critical for the development and function of rod and cone photoreceptors. Genes in bold were confirmed using both the methylKit and DMRseq packages. **D)** Percentages of genes whose TSSs±500 bp statistically significantly increase or decrease methylation in pairwise comparisons of RPCs, rod, and cone methylomes were calculated using the methylKit and DMRseq packages.

### Methylation of photoreceptor gene TSSs inhibits their expression both directly and indirectly

Our results indicate that in RPCs, photoreceptor gene TSSs are located in highly methylated regions, the methylation of which must be reduced for these genes to be activated in rods and cones. However, to fully understand the mechanisms governing photoreceptor gene expression, we must pinpoint the genomic locations of specific cytosines whose methylation status may influence the activity of these genes. Methylation of cytosines located near a gene’s TSS can lead to a significant reduction in gene expression, either by inducing chromatin compaction (indirect mechanism) or by hindering the binding of specific transcription factors (direct mechanism)^27,28^. We utilized the DSS Bioconductor package to identify all cytosines exhibiting statistically significant changes in methylation levels near the TSSs of rod photoreceptor genes in a comparative analysis of rods vs. RPCs (E11), rods vs. Chx10TET, TET vs. RPCs, and TET vs. Chx10TET. We focused exclusively on rod-specific genes, as the cone methylome was not discernible in the comparative analysis (**Fig. 6B**). We found multiple cytosines located in the TSSs±1,000 bp (**Supplementary Datafile S11**). Those cytosines that are highly methylated in RPCs (E11) remain highly methylated in TET-deficient (Chx10TET) retinas (**Fig. 8A**, **Supplementary Datafile S11**). The differences in methylation—and the abundance of specific cytosines—were particularly evident when comparing the methylomes of rods vs. RPCs (E11) and rods vs. Chx10TET (**Fig. 8A**, **Supplementary Datafile S11**). The genomic localization of these significant cytosines suggests that their methylation status may influence photoreceptor gene expression through both direct and indirect mechanisms. To elucidate which mechanism inhibits photoreceptor gene expression in RPCs and TET-deficient (Chx10TET) retinas, we assessed the level of chromatin compaction using ATAC-seq. To this end, we utilized P14 Chx10TET (n=3) and TET (n=3) retinas, as well as previously published ATAC-seq data obtained from RPCs and rods^26,29^. Pairwise comparison of ATAC-seq data using the DiffBind Bioconductor package revealed minimal differences between Chx10TET and TET retinas and maximal differences between rods and RPCs (**Fig. 8B, 8C**, **Supplementary Datafile S11**). It is interesting to note that there are more genomic regions in which the chromatin is more open in RPCs than in rods, Chx10TET, and TET (up, **Fig. 8C**). At the same time, there are more TSSs±3,000 bp in which the chromatin is more open in rods, Chx10TET, and TET than in RPCs (down, **Fig. 8C**). Using the ATAC-seq and DiffBind data, we identified those rod photoreceptor genes for which the level of chromatin compaction at TSSs±1,000 bp differs pairwise in RPCs, rods, Chx10TET, and TET (**Supplementary Datafile S11**). We found that chromatin is more compact for a small subset of the genes in TET-deficient retinas compared to controls (**Fig. 8D**, **Supplementary Datafile S11**). Given that the retina contains more than just photoreceptors, we found that chromatin is more open at TSSs (±1,000 bp) of these genes in rods vs. Chx10TET and rods vs. TET (**Fig. 8D**). However, the most significant finding is that the chromatin in RPCs is more compact at TSSs (±1,000 bp) of the photoreceptor genes compared to the chromatin in rods, Chx10TET, and TET (**Fig. 8D**, **Supplementary Datafile S11**). Our results suggest that cytosine methylation near TSSs of rod photoreceptor genes leads to chromatin compaction in RPCs (closed chromatin). Chromatin is open at TSSs (±1,000 bp) of photoreceptor genes in TET-deficient (Chx10TET) retinas. However, the presence of methylated cytosines in these regions prevents the expression of the rod photoreceptor genes. An important observation is that the inactivation of TET enzymes in RPCs does not prevent the opening of chromatin at TSSs (±1,000 bp) of rod photoreceptor genes in TET-deficient (Chx10TET) retinas.

**Figure 8.**
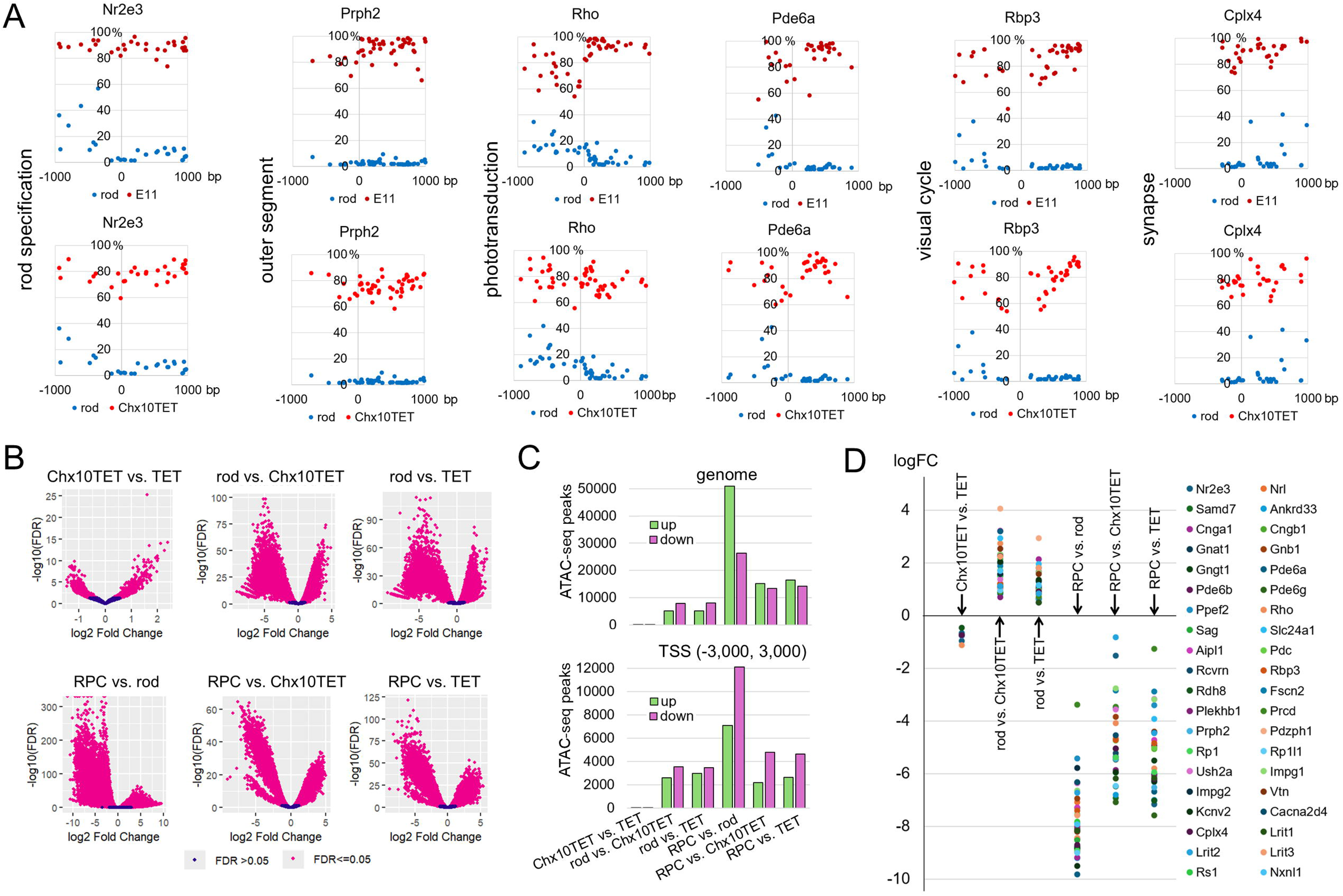
Chromatin in hypermethylated photoreceptor gene TSSs is in a closed state in RPCs, while chromatin in the same DNA regions is in an open state in TET-deficient retinas. **A)** The panels show examples of photoreceptor genes with differentially methylated cytosines (FDR < 0.001) at their TSSs (±1,000 bp). The methylomes of RPCs (E11), rods, Chx10TET, and TET were compared using the DSS package. **(B, C)** The ATAC-seq peaks were obtained using MACS2 software during the analysis of raw ATAC-seq data. The volcano plots (**B**) were generated by the DiffBind package during a search for genomic regions differing in chromatin compaction levels (**C**) in RPCs (E11), rods, Chx10TET, and TET. When constructing graphs (**C**), only those ATAC-seq peaks were used for which logFC > 1 (up, chromatin is open) or logFC <-1 (down, chromatin is closed). (FDR <0.05, logFC – Log Fold Change) **D)** The results of the ATAC-seq data analysis, utilizing the DiffBind package, were applied to rod photoreceptor genes.

## Discussion

The retina is a multi-layered neural tissue located in the eye that is responsible for coding light information into electrical signals that the brain uses to form the sensation of vision^1–4^. The highly organized structure and cellular composition of the retina are critical to its function, and any disruptions due to impaired development lead to blindness^1–5^. In this study, we investigated how the TET-dependent DNA demethylation pathway controls retinal development. We found that developmental abnormalities in TET-deficient retinas emerge as early as the embryonic stage. These abnormalities are marked by delayed retinal growth, the low thickness of the ONbL, which contains RPCs, and the enormous thickness of the GCL, which contains non-dividing precursors at this stage of development. TET-deficient RPCs exhibit high rates of proliferation and continue to divide even during late stages of retinal development—long after division has ceased in control retinas. These results could be explained by the fact that TET-deficient RPCs possess characteristics of both early and late RPCs—a factor that could account for the observed cellular composition of TET-deficient mature retinas whose development was driven by these RPCs. Beyond the level of RPCs themselves, the TET-dependent DNA demethylation pathway regulates the development and function of various retinal cell types, a role that is particularly evident in the case of rod and cone photoreceptors. Our results indicate that this broad regulatory control exerted by the TET-dependent DNA demethylation pathway over retinal development can be attributed to the existence of many critical for retinal development genes whose high levels of methylation in RPCs directly and indirectly suppress their expression.

Early progenitors exhibit a high rate of proliferation and symmetric self-renewal^30–32^. This leads to an exponential increase in cell number and rapid tissue growth. Early progenitors are also capable of differentiating into all types of tissue cells directly or indirectly by differentiating into late progenitors, which in turn differentiate into cell types that appear later in development^30–32^. A defining characteristic of late progenitors is that they divide slowly and asymmetrically, generating one progenitor and one non-dividing precursor of a certain cell type^30–32^. RPCs possess all these properties (**Fig. 9A**)^3,20,33,34^. In our study, we found that inactivation of the TET-dependent DNA demethylation pathway in RPCs results in slow retinal growth, a small number of RPCs in the ONbL, and a large number of non-dividing cells in the GCL, starting from the earliest stage of retinal development. This phenomenon could be explained by the predominantly asymmetric division of TET-deficient RPCs already at an early stage of retinal development, which is characteristic of late RPCs (**Fig. 9A**). Such asymmetric division—yielding one dividing RPC and one non-dividing precursor—would result in a more linear (rather than exponential) growth of the retina and the subsequent accumulation of non-dividing cells within the GCL. At the same time, however, our EdU tests and RNA-seq data indicate that TET-deficient RPCs maintained a high rate of proliferation, continuing to divide even during late stages of retinal development — a phase during which cell division could no longer be detected in the control samples. The high rate of proliferation is a characteristic typically associated with early RPCs. Increased expression of *Zic1*, a transcription factor which supports RPC proliferation, could explain, in part, these observations^21^. These findings strongly suggest that TET-deficient RPCs exhibit characteristics associated with both early and late RPCs. Thus, the TET-dependent DNA demethylation pathway prevents the mixing of early and late RPC phenotypes and facilitates their manifestation in accordance with the stage of retinal development.

**Figure 9.**
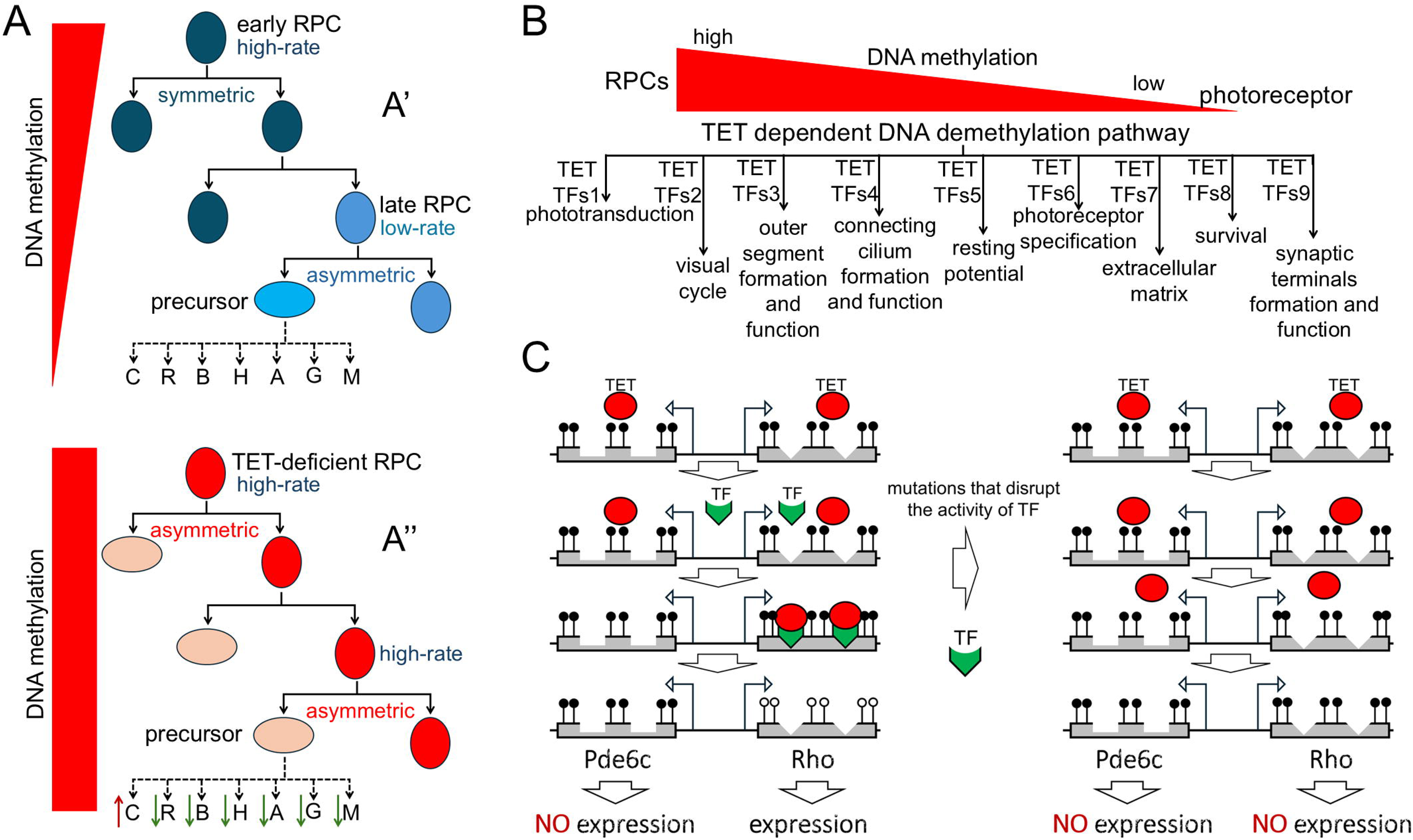
Many groups of still unknown transcription factors (TFs) orchestrate TET-dependent demethylation of hypermethylated in RPCs genes critical for retinal development. **A)** Early RPCs divide predominantly symmetrically and exhibit a high proliferation rate (**A’**). This leads to rapid retinal growth. Late RPCs divide predominantly asymmetrically and exhibit a low proliferation rate (**A’**). TET-deficient RPCs divide predominantly asymmetrically but exhibit a high proliferation rate (**A’’**). This results in slow retinal growth, yet ensures the presence of dividing cells even at the very latest stage of retinal development. (C- cones, R – rods, B – bipolar cells, H – horizontal cells, A – amacrine cells, G – RGCs, M – Muller glia) **B)** The activity of the TET-dependent DNA demethylation pathway requires several groups of TFs responsible for photoreceptor gene demethylation. For instance, one such group of TFs assists TET enzymes in demethylating genes responsible for photoreceptor specification; another group is responsible for demethylating genes involved in phototransduction; and so on. **C)** TET enzymes require assistance from TFs that bind to TET enzymes and deliver them to specific hypermethylated genes for demethylation. If a mutation disrupts the activity of a TF responsible for the demethylation of a gene critical for retinal development and function, it will lead to a lack of its expression and, as a result, to IRD.

Our results also indicate that the TET-dependent DNA demethylation pathway regulates the differentiation of RPCs into various retinal cell types. We observed a high abundance of early-born S-cone photoreceptors in TET-deficient retinas. Conversely, the numbers of other retinal cell types—both early-born and late-born—were reduced. This unbalanced distribution of early- and late-born retinal cell types can likely be attributed to the mixed phenotype of TET-deficient RPCs. However, the influence of the TET-dependent DNA demethylation pathway is not limited merely to altering the composition of retinal cell populations; the inactivation of TET enzymes in RPCs impacts the development of all retinal cell types^17–19^. This effect is particularly pronounced in the development of rod and cone photoreceptors. Our results explain this phenomenon by high levels of methylation in RPCs of many genes required for the development and function of rods and cones. The TET-dependent DNA demethylation pathway is responsible for the demethylation of these genes in the developing retina, leading to their activation in the photoreceptors. A subset of these genes is also essential for the development of other retinal cell types (e.g., bipolar cells); consequently, the absence of demethylation of these genes in TET-deficient retinas likely impairs the development of these cell types. We also found that there were differences in gene demethylation depending on the retinal cell type. While many genes that are highly methylated in RPCs undergo demethylation in both rods and cones, there exist distinct groups of genes that are demethylated exclusively in rods or exclusively in cones. This implies that there is a degree of specificity in the activity of the TET-dependent DNA demethylation pathway within the precursors of rods and cones. Such specificity likely exists both within RPCs themselves and within other retinal cell types differentiating from them.

One of the key questions we sought to address was how high levels of gene methylation in RPCs and TET-deficient retinas impede gene expression. We focused on highly methylated rod-specific genes, which are easier to study given the high abundance of rods within the retina. We identified many methylated cytosines located near TSSs of genes responsible for rod development and function. In this context, two scenarios are plausible. First, methyl-binding proteins (MBPs) could bind to the methylated cytosines, thereby stimulating chromatin compaction and consequently inhibiting gene expression^27^. The second scenario is more specific, positing that cytosine methylation may directly interfere with the binding of transcription factors required to activate gene expression^28^. We investigated the degree of chromatin compaction in RPCs, rods, TET-deficient retinas, and control retinas. We discovered that in RPCs, chromatin exists in a “closed” state at the highly methylated TSSs of 40 genes essential for rod (and cone) photoreceptor development and function. Conversely, in TET-deficient retinas, chromatin at these same highly methylated TSSs was found to be in an “open” state. In the genomes of rods and control retinas, chromatin was in an “open” state, and methylation levels were low at TSSs of the same genes. These findings indicate that high levels of photoreceptor gene methylation in RPCs lead to chromatin compaction. During retinal development, chromatin opens and the TET-dependent DNA demethylation pathway has no influence on this process. Subsequently, TET-dependent demethylation of the TSSs is required to enable specific transcription factors to bind to them and initiate the expression of rod (and cone) photoreceptor genes^28^. Thus, there exists a multi-step process governing the initiation of photoreceptor gene expression, in which the participation of the TET-dependent DNA demethylation pathway is necessary but not sufficient. However, it is crucial to note that the methylation of photoreceptor genes controls their expression both directly and indirectly.

The totality of data collected in this study, combined with previously published findings, indicates that the TET-dependent DNA demethylation pathway exerts global control over retinal development and function^17–19^. However, there is a paradox: TET enzymes are unable to bind to hypermethylated DNA^35–41^. Consequently, TET enzymes require the assistance of transcription factors (TFs) to help them bind to the TSS of a specific gene for its demethylation. These TFs are responsible for the specificity of gene demethylation. This mechanism has already been demonstrated in many tissues but requires further study in the developing retina^42–52^. This mechanism is of particular importance during the development of rods and cones. We have demonstrated that, in RPCs, the genes responsible for photoreceptor specification, phototransduction, visual cycle as well as for the formation and function of connecting cilium, synapses, and outer segments, are highly methylated (**Fig. 6E, 7C**). Thus, the development of functional photoreceptors is impossible without the demethylation of these genes—a conclusion strongly supported by our data^17^. However, these genes can undergo demethylation only in the presence of as-yet-unidentified TFs that assist TET enzymes in binding to the TSSs of these genes (**Fig. 9B**). Another reason why it is important to identify these TFs is that they are most likely involved in inherited retinal diseases (IRD). For example, if mutations disrupt the activity of the TF responsible for TET-dependent demethylation of the rhodopsin (Rho) gene TSS, this will lead to a lack of Rho expression and, as a result, to IRD (retinitis pigmentosa, **Fig. 9C**). However, in this case, the IRD is caused not by mutations in a gene (e.g., Rho) but by high levels of its methylation. We call this form of IRD the epigenetic form of IRD. So, we can distinguish two forms of IRD: genetic and epigenetic. Mutations that disrupt the activity of IRD gene (e.g., Rho, Prph2, Pde6a, Pde6b, etc.) lead to genetic forms of IRD. Mutations that disrupt the activity of TF responsible for TET-dependent IRD gene demethylation lead to epigenetic form of IRD. Genetic IRD and epigenetic IRD are likely to be phenotypically similar. The reason these forms of IRD should be phenotypically similar is that the IRD gene is inactive in both cases. The difference is that in the genetic form of IRD, the IRD gene is inactive due to a mutation. In the epigenetic form of IRD, the IRD gene is inactive due to high levels of methylation of its TSS.

In conclusion, we have demonstrated that the TET-dependent DNA demethylation pathway exerts global control over retinal development at all levels. It maintains the phenotype of early and late RPCs and further promotes RPC differentiation into various retinal cell types—most notably rods and cones. Mutations in genes involved in the TET-dependent DNA demethylation pathway are expected to result in epigenetic forms of IRD. A particularly significant role in this process is likely played by TFs, which assist TET enzymes in demethylating genes critical for retinal development and function.

## Materials and Methods

### Ethical approvals

The procedures were performed in compliance with the NIH Guide for the Care and Use of Laboratory Animals, ARVO statement for the Use of Animals in Ophthalmic and Vision Research, ARRIVE (Animal Research: Reporting of In Vivo Experiments) guidelines, and according to the University of Miami Institutional Animal Care and Use Committee (IACUC) approved protocol #: 23-058. Animals were euthanized according to the recommendations of the Panel on Euthanasia of the AVMA.

### Animals

Tet1-floxed, Tet2-floxed, and Tet3-floxed mice were obtained from Dr. Anjana Rao (La Jolla Institute for Immunology) and were crossbred to produce TET triple floxed mice (they are designated in the text as TET). These animals were used as controls and also to generate TET conditional knockouts. To this end, Chx10-Cre mice (strain 005105, the Jackson Laboratory, Bar Harbor, ME, USA) and TET triple floxed mice were crossbred to produce TET triple conditional knockout mice (they are designated in the text as Chx10TET). Chx10TET mice were heterozygous for Chx10-Cre allele and homozygous for Tet1-floxed, Tet2-floxed, and Tet3-floxed alleles. Chx10TET and TET mice have the C57BL/6J genetic background. Embryonic retinas were collected from embryonic day 11 (E11) C57BL/6J mice (strain 000664, the Jackson Laboratory, Bar Harbor, ME, USA). Both females and males were used to address sex as a biological variable. The animals were kept in standard housing conditions on a 12-hour light/dark cycle with unrestricted access to food and water.

### Tissue collection

To collect retinas without contamination by blood cells, mice were perfused with phosphate buffered saline (PBS, pH 7.4; #10010023, ThermoFisher Scientific, Waltham, MA, USA). To this end, the mice were IP injected with ketamine (80 mg/kg) and xylazine (10 mg/kg). When the mice were under deep anesthesia, the chest cavity was opened, and a syringe needle was placed into the heart. The syringe needle was attached to a pump to allow PBS to circulate through the mouse. Eyes were then dissected out and retinas were isolated. The retinas were collected in the appropriate buffers for RNA and DNA purification, and immunohistochemistry.

### Immunohistochemistry

Enucleated eyes were fixed in 4% paraformaldehyde (PFA) overnight at 4°C. The fixed eyes were placed in 30% sucrose/phosphateLbuffered saline (PBS). After 24 hours, the eyes were cryoLembedded in OCT and cryoLsectioned at 10Lμm thickness. We then permeabilized the sections with 0.3% Triton XL100/PBS for 1Lhour, washed with PBS, and blocked in a buffer containing 5% donkey serum, 2% BSA, and 0.15% TweenL20 in PBS. After 1-hour, primary antibodies and/or PNA (L32458, ThermoFisher Scientific, Waltham, MA, USA) in the blocking buffer were added to the sections. We used the following antibodies: anti-Sox2 (1:600, ab97959, Abcam, Waltham, MA, USA), anti-Chx10 (1:200, AB9016, MilliporeSigma, Burlington, MA, USA), anti-Rho (1:100, PA5-85608, ThermoFisher Scientific, Waltham, MA, USA), anti-ChAT (1:200, AB114P, MilliporeSigma, Burlington, MA, USA), anti-Cplx4 (1:200, 122 402, Synaptic Systems GmbH, Goettingen, Germany), anti-Cplx3 (1:500, 122 302, Synaptic Systems GmbH, Goettingen, Germany), anti-Dlg4 (1:500, GTX133255, GeneTex, Irvine, CA, USA). The sections were incubated overnight at 4°C. The next day, the retinal sections were washed with PBS and incubated with secondary fluorescent antibodies (ThermoFisher Scientific, Waltham, MA, USA). Control sections were incubated without primary antibodies. Hoechst 33342 (H3570, ThermoFisher Scientific, Waltham, MA, USA) was applied to stain the cell nuclei. Images were collected and analyzed using Leica STELLARIS confocal microscope (Leica Microsystems, Deerfield, IL, USA) and its software.

### EdU cell proliferation assay in vivo

To label proliferating cells in the developing retina we used EdU-Click kit (BCK-EDU647, MilliporeSigma) according to the manufacturer’s instructions. To this end, the EdU (50mg/kg in 1X PBS) solution was injected intraperitoneally into pregnant mice (E16) or pups (P0-P10). After 4 hours (pulse) or 30 min (short pulse), the retinas were collected and used in the immunohistochemistry analysis.

### Whole genome bisulfite sequencing (WGBS) library preparation

The genomic DNA (gDNA) was purified from retinas using the DNeasy Blood and Tissue Kit (#69504, Qiagen, Hilden, Germany). The quantity and quality of gDNA was assessed using the NanoDrop One Spectrophotometer, Qubit 4 Fluorometer (ThermoFisher Scientific, Waltham, MA, USA), and 4200 TapeStation system (Agilent Technologies, Santa Clara, CA, USA). To prepare WGBS libraries, we used xGen MethylLSeq DNA Library Prep Kit (#10009860, Integrated DNA Technologies, Coralville, IA, USA) according to manufacturer’s instructions. The WGBS libraries were quantified using Qubit 4, and sizes were assessed using the 4200 TapeStation system. The WGBS libraries were sequenced on the Illumina NextSeq2000 using 2L×L150 paired end (PE) configuration. The generated FASTQ files have been deposited in the BioProject database (ncbi.nlm.nih.gov/bioproject/) and can be accessed using the accession number PRJNA1286813.

### WGBS data analysis

FastQC (v0.11.8) quality control tool was used to assess the overall quality of each sequenced sample. TrimGalore (v0.4.5) with the following parameters: –adapter AGATCGGAAGAGC Le 0.1 –stringency 6 –length 20 –nextseq 20 –three_prime_clip_R1 10 –clip_R2 10 was used for preprocessing our FASTQ files. To align trimmed PE reads and calculate the percentage of cytosine methylation, the Bismark program and GENCODE M25 (GRCm38.p6) Mus musculus reference (mm10) genome were used^53^. We have applied the DSS Bioconductor package, to calculate the average base coverage of the mouse genome^54^. The average base coverage of the mouse genome was 20X for E11_1, 20X for E11_2, 22X for E11_3, 10X for TET_1, 11X for TET_2, 11X for TET_3, 10X for Chx10TET_1, 12X for Chx10TET_2, 16X for Chx10TET_3, 8X for SRR9833662_rod3m1, 9X for SRR9833663_rod3m2, 9X for SRR9833664_rod3m3, 5X for SRR3933743_rod1_1, 5X for SRR3933744_rod2_1, 19X for SRR2722846_rod1, 11X for SRR2722847_rod2, 4X for SRR3933741_cone1_1, 4X for SRR3933742_cone2_1, 16X for SRR2722850_cone1, 13X for SRR2722851_cone2. We utilized the methylKit Bioconductor package to identify genomic regions exhibiting distinct average levels of DNA methylation within the same genome^55^. To detect differentially methylated cytosines, we used the DSS Bioconductor package. To detect differentially methylated regions (DMRs), we used DMRseq Bioconductor package^56^. To find promoters located in DMRs, we used EPD and EPDnew databases^57^. To generate Venn diagrams, we used the Venny2.1 interactive tool (https://bioinfogp.cnb.csic.es/tools/venny/). We used the ShinyGO graphical gene-set enrichment tool for gene ontology enrichment (GO) analysis (ver. 0.82, http://bioinformatics.sdstate.edu/go/).

### Bulk RNA-seq library preparation and sequencing

Total RNA was purified from retinas using RNeasy Plus Mini Kit (#74134, Qiagen, Hilden, Germany)^58^. The quantity and quality of RNA was assessed using the NanoDrop One Spectrophotometer and Qubit 4 Fluorometer. To assess RNA integrity, we used the 2100 Bioanalyzer Instrument (Agilent Technologies, Santa Clara, CA, USA). RNA samples with a RIN (RNA Integrity Number) score of 8 or higher were used to prepare RNALseq libraries. The mRNA-seq library preparation Kit (RK20302, ABclonal, Woburn, MA, USA) was utilized according to manufacturer’s instructions to prepare RNALseq libraries, which were sequenced on the Illumina NovaSeq X Plus using 2L×L150 PE configuration. The FASTQ files have been deposited in the BioProject database (ncbi.nlm.nih.gov/bioproject/) and can be accessed using the accession number PRJNA1286813.

### Bulk RNA-seq data analysis

We used the STAR RNA-seq aligner, HTseq package, and GENCODE M25 (GRCm38.p6) Mus musculus reference genome (mm10) to compute how many reads overlap each of the mouse genes^59,60^. The edgeR Bioconductor package was applied to perform the differential gene expression analysis^61^. We applied the DESeq2 Bioconductor package to perform the principal component analysis (PCA) and sample clustering^62^.

### Single-nucleus RNA library preparation, sequencing and data analysis

Retinas were collected from P14 TET and Chx10TET mice and immediately frozen. Cell nuclei from the frozen retinas were isolated using the Nuclei Extraction Buffer (130-128-024, Miltenyi Biotec, Gaithersburg, MD, USA) and used to generate the 10X snRNA-seq library according to the manufacturer’s instruction (Chromium GEM-X Single Cell 3’ Reagent Kits v4, 10x Genomics, Pleasanton, CA, USA). Briefly, the GEM generation and barcoding were done with a 10x Chromium X series. mRNA from the uniquely barcoded cell nuclei was subsequently reverse transcribed into cDNA, which was then amplified to generate sufficient amount for snRNA-seq library construction. Next the cDNA was used for the bulk library preparation, including enzymatic fragmentation, size-selection, end-repair, A-tailing, adaptor ligation and dual index PCR. Finally, the library was ready for quality control and followed sequencing on Illumina Novaseq X plus platform using PE150 strategy. The FASTQ files generated in the study have been deposited in the BioProject database (ncbi.nlm.nih.gov/bioproject/) and can be accessed using the accession number PRJNA1286813. To demultiplex next-generation sequencing (NGS) data (FASTQ files), process barcodes, align reads to GRCm38.p6/mm10 (GENCODE M25 release) reference genome, and generate feature-barcode matrices, we used the Cell Ranger software suite developed by 10x Genomics. The output from Cell Ranger was used in R toolkit Seurat (v5.2.0) for clustering, cell type annotation, differential expression analysis, and visualization. During quality control, only those cells that met the following conditions were retained: nCount_RNA > 500 & nFeature_RNA > 500 & mitoPercent < 5 & riboPercent < 5. In order to identify doublets and remove them we used the scDblFinder R Bioconductor package^63^. We used the SoupX R package to remove ambient RNA contamination from our snRNA-seq data^64^. The R code that was used in snRNA-seq analysis can be found in Supplementary Datafile S12.

### ATAC-seq library preparation and sequencing

ATAC-seq library construction was performed as previously reported^65,66^. Briefly, nuclei were extracted from frozen retinas, and the nuclei pellet was resuspended in the Tn5 transposase reaction mix. The transposition reaction was incubated at 37°C for 30 min. Adapters were added after transposition and PCR was performed to amplify the library. After the PCR amplification, libraries were purified with AMPure beads. The libraries were checked with Qubit and quantitative PCR for quantification and Bioanalyzer for size distribution detection. Quantified libraries were pooled and sequenced on the Illumina NovaSeq X Plus using 2L×L150 PE configuration. The FASTQ files have been deposited in the BioProject database (ncbi.nlm.nih.gov/bioproject/) and can be accessed using the accession number PRJNA1286813.

### ATAC-seq data analysis

TrimGalore (v0.4.5) was used for preprocessing the FASTQ files. To align trimmed PE reads, the Bowtie 2 software and GENCODE M25 (GRCm38.p6) Mus musculus reference (mm10) genome were used. The Picard (MarkDuplicates) software was used for PCR duplicate removal. To call peaks, we used the MACS2 software^67^. The bam and narrowPeak files generated by the MACS2 software were used in a comparative analysis utilizing the DiffBind Bioconductor package^68^. Prior to the comparative analysis using the DESeq2 algorithm, the data were normalized using these parameters: method = DBA_DESEQ2, normalize = DBA_NORM_RLE. The comparison results (FDR < 0.5) were annotated using the annotatePeak function (ChIPseeker, version 1.8.6)^69^.

### Statistical analysis

The Student’s tLtest was applied for experiments containing one variable. We considered PLvalues and FDR equal to or less than 0.05 statistically significant. Generation and analysis of nextLgeneration sequencing data were performed inLhouse according to ENCODE standards and pipelines. The relevant details can be found above.

## Supporting information

Supplementary Datafile S1

Supplementary Datafile S2

Supplementary Datafile S3

Supplementary Datafile S4

Supplementary Datafile S5

Supplementary Datafile S6

Supplementary Datafile S7

Supplementary Datafile S8

Supplementary Datafile S9

Supplementary Datafile S10

Supplementary Datafile S11

Supplementary Datafile S12

## Data availability

The datasets collected and analyzed in the study are available in the BioProject database (accession numbers PRJNA1286813 and PRJNA1121391) and in the article/Supplementary Datafiles. We also used NCBI Sequence Read Archive (SRA) datasets: 1) WGBS data: rods (SRR9833662, SRR9833663, SRR9833664, SRR3933743, SRR3933744, SRR2722846, SRR2722847) and cones (SRR3933741, SRR3933742, SRR2722850, SRR2722851); 2) ATAC-seq data: RPCs (SRR7734014, SRR7734015, SRR7734016, SRR7734017) and rods (SRR2721946, SRR2721948).

## Code availability

The R code that was used in snRNA-seq analysis can be found in Supplementary Datafile S12.

## Acknowledgments

This study was supported in part by the National Institutes of Health/National Eye Institute grant R01 EY035235 (D.I.), National Institutes of Health/National Eye Institute Center Core grant P30 EY014801, Research to Prevent Blindness/Unrestricted Grant GR004596-1, and the Mark J. Daily Inherited Retinal Diseases Research Center at Bascom Palmer Eye Institute. We are deeply grateful to Patricia and Jim Derryberry for their generous support of this research. We are sincerely grateful to Dr. Anjana Rao (La Jolla Institute for Immunology) for the TET triple-floxed mice she provided to us. We thank Dr. Kurtenbach and Dr. Dollar for their advice on snRNA-seq data analysis. The authors thank Charles K. Yaros for his expert assistance.

## Conflicts of interest

The authors declare no conflicts of interest.

## Author contributions

D.I. conceived and supervised the project. G.D., M.F., and D.I. performed the experiments and the data analyses. G.D., and D.I. assisted with the bioinformatic analysis. G.D., M.F., B.L.L., and D.I. assisted with the research design, data interpretation, manuscript writing and editing. All authors have read and agreed to the published version of the manuscript.

## Notes

### Competing Interest Statement

The authors have declared no competing interest.

