## Supplementary figures and images for "Retinal development driven by TET-dependent DNA demethylation"

### Supplementary Datafile S8

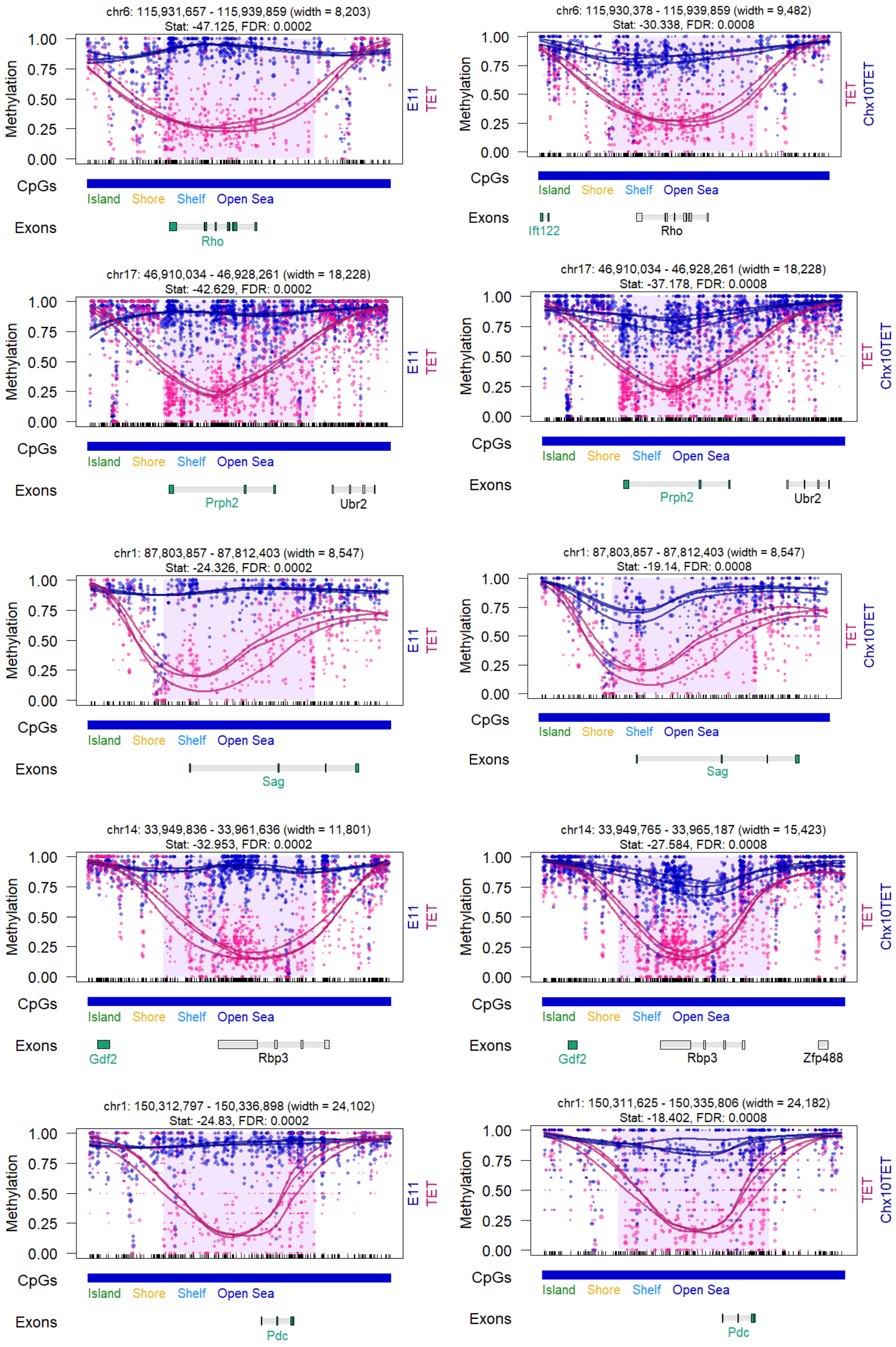

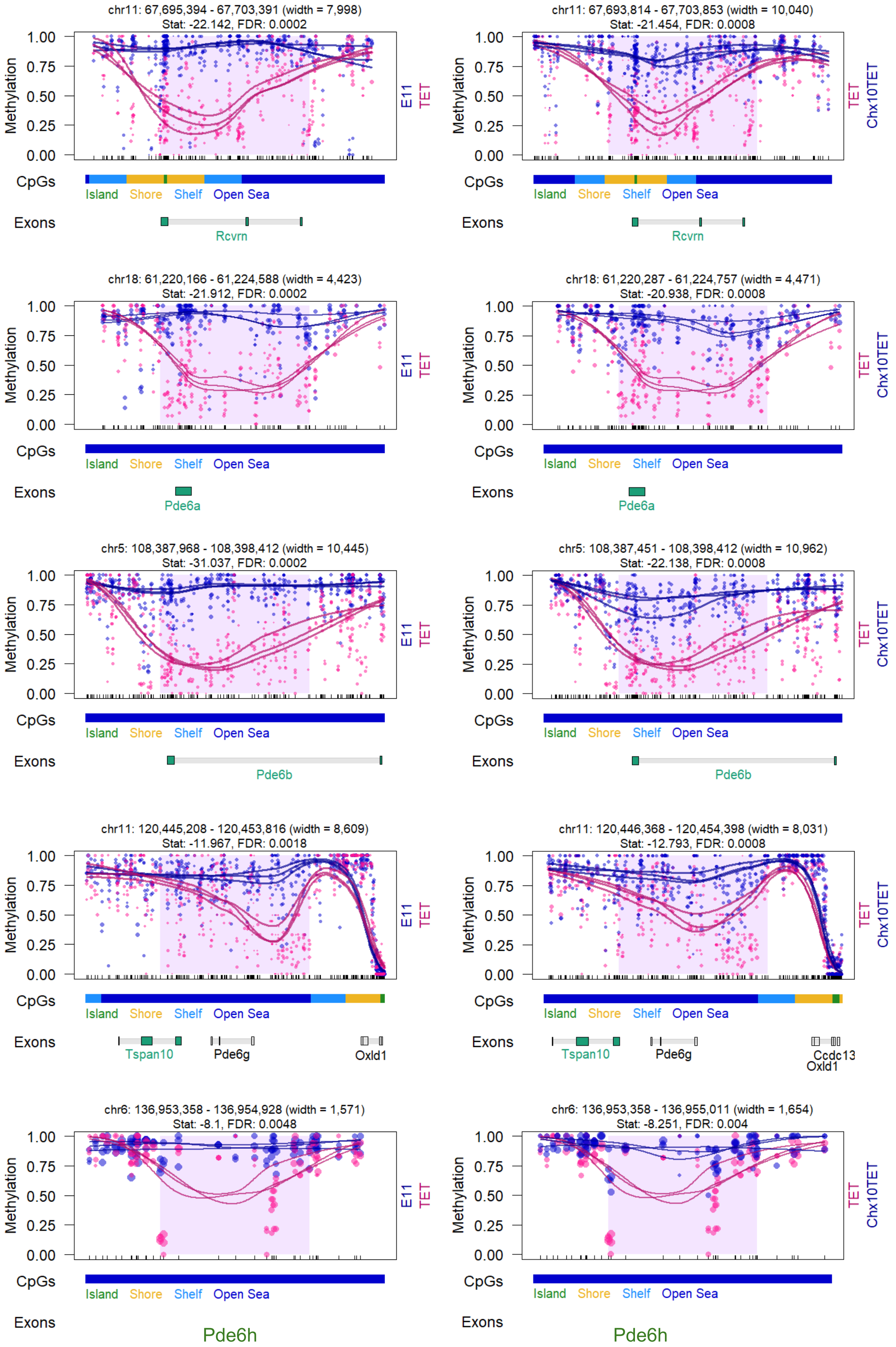

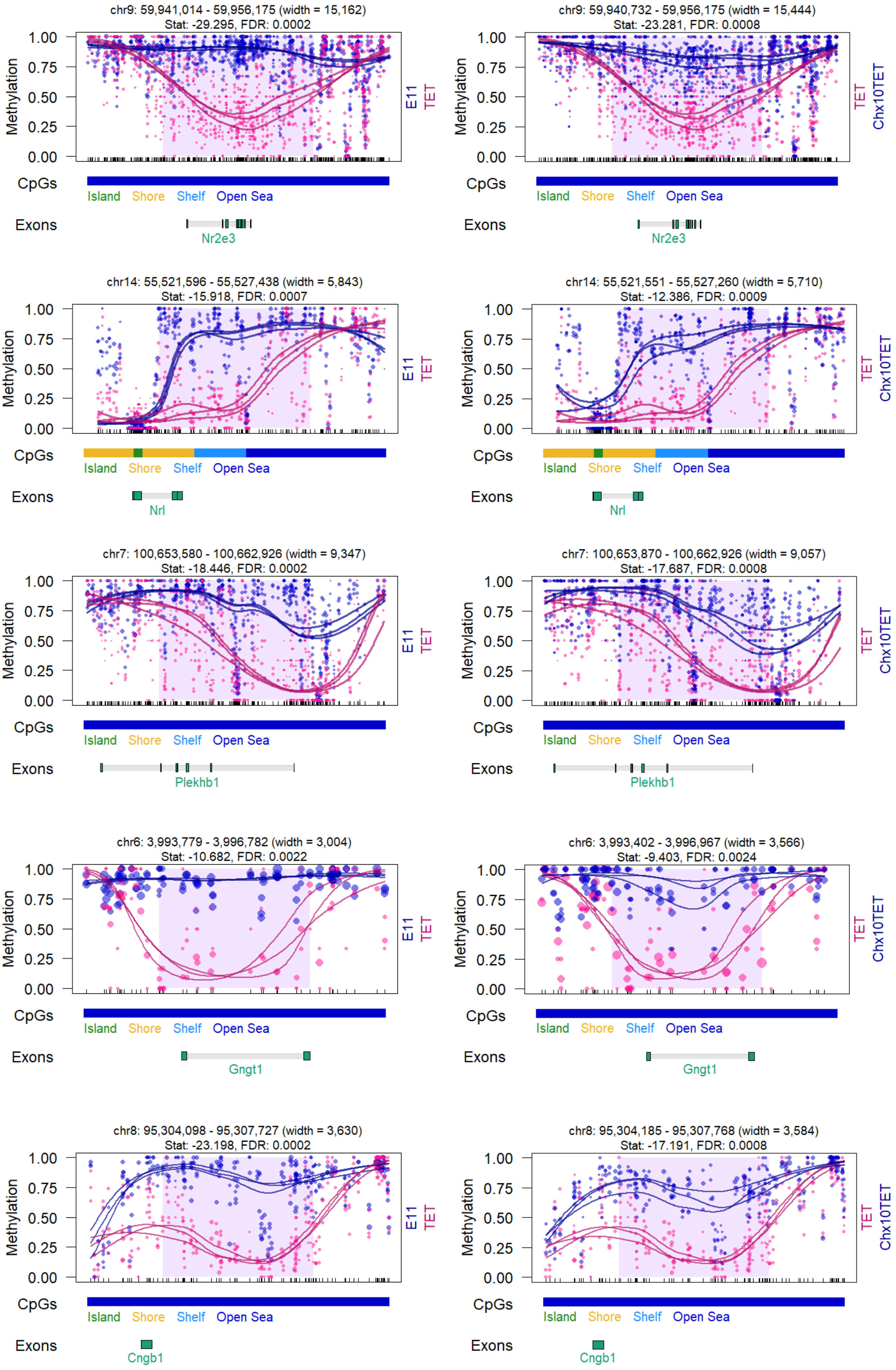

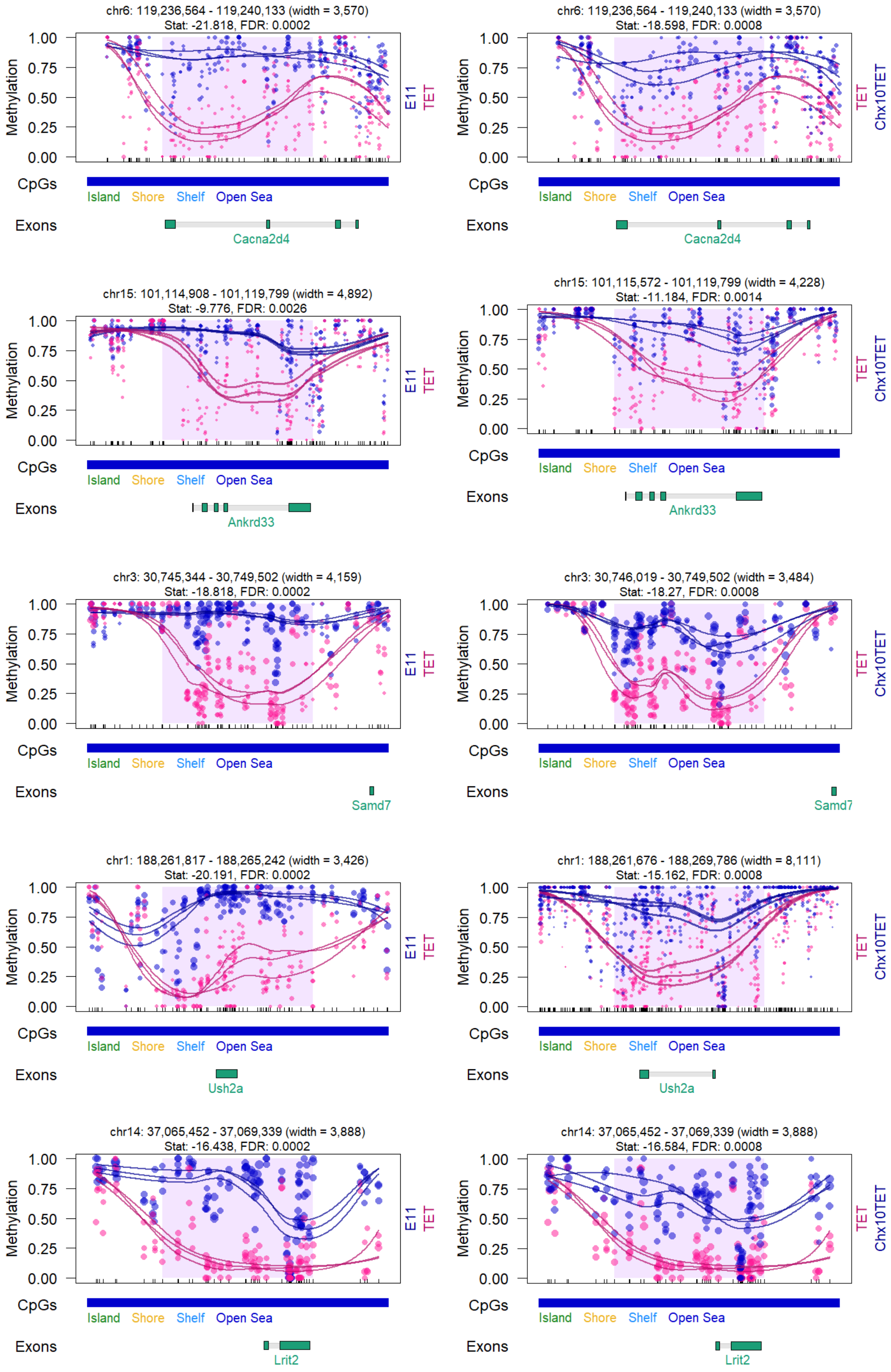

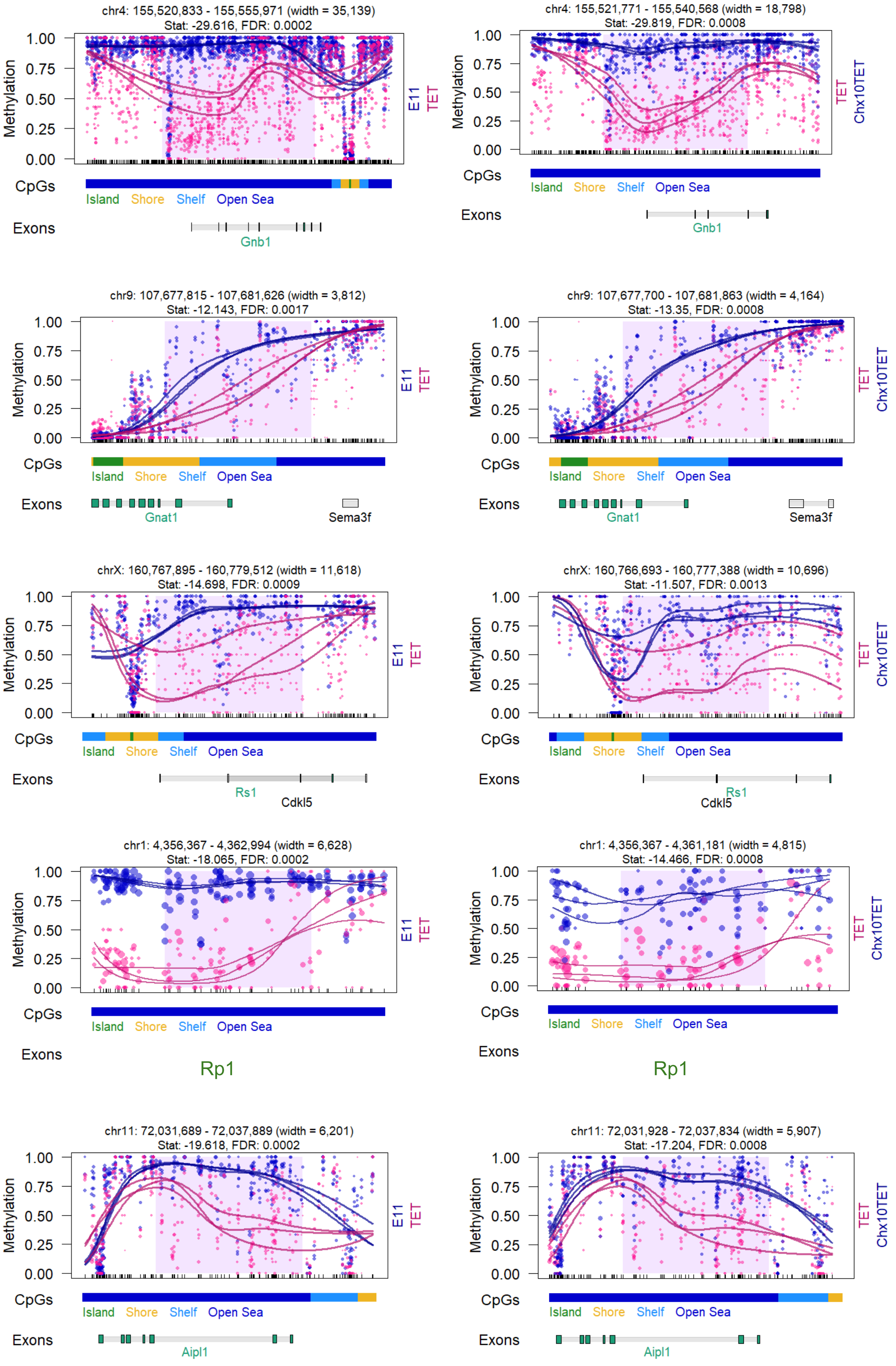

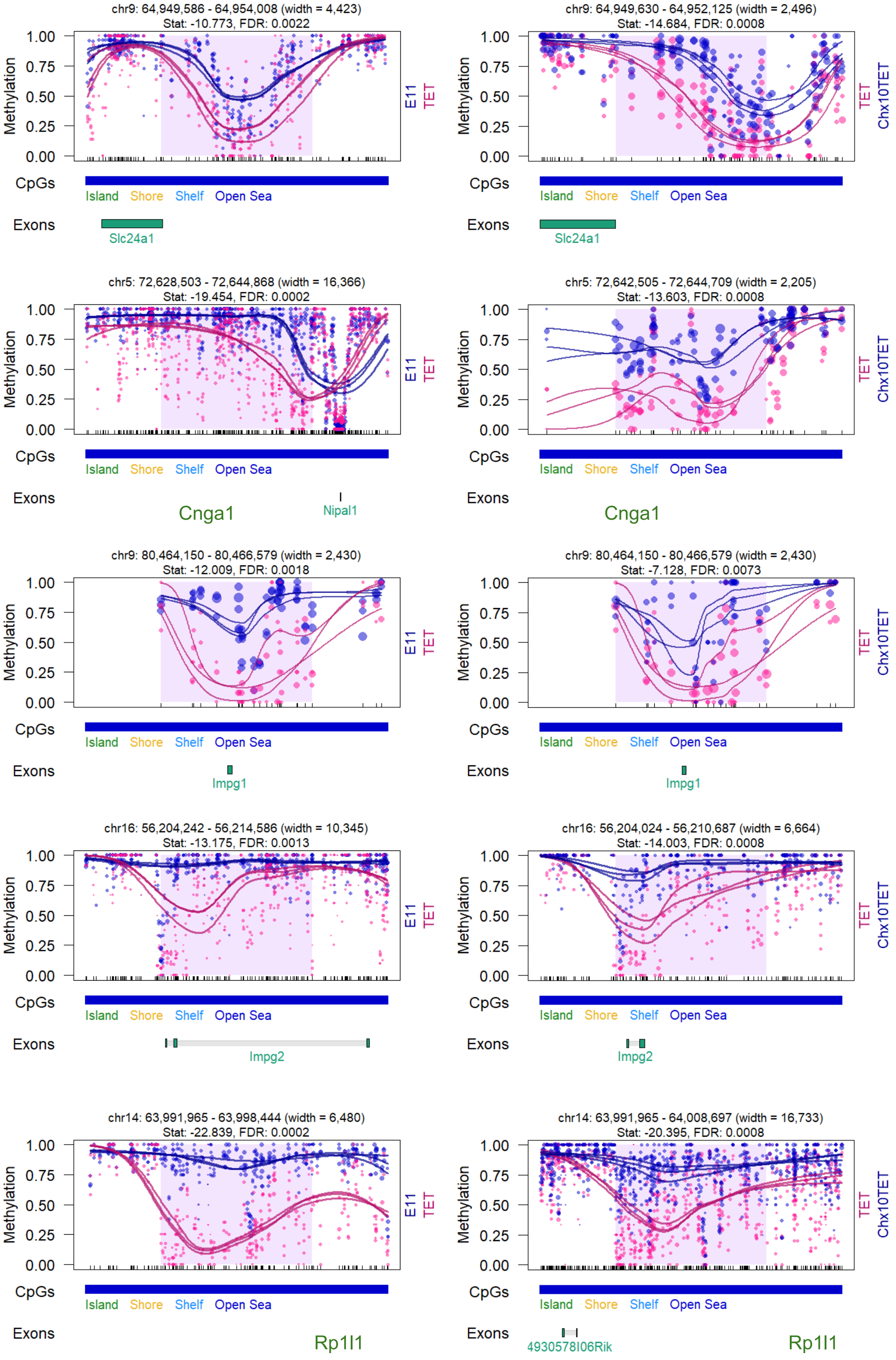

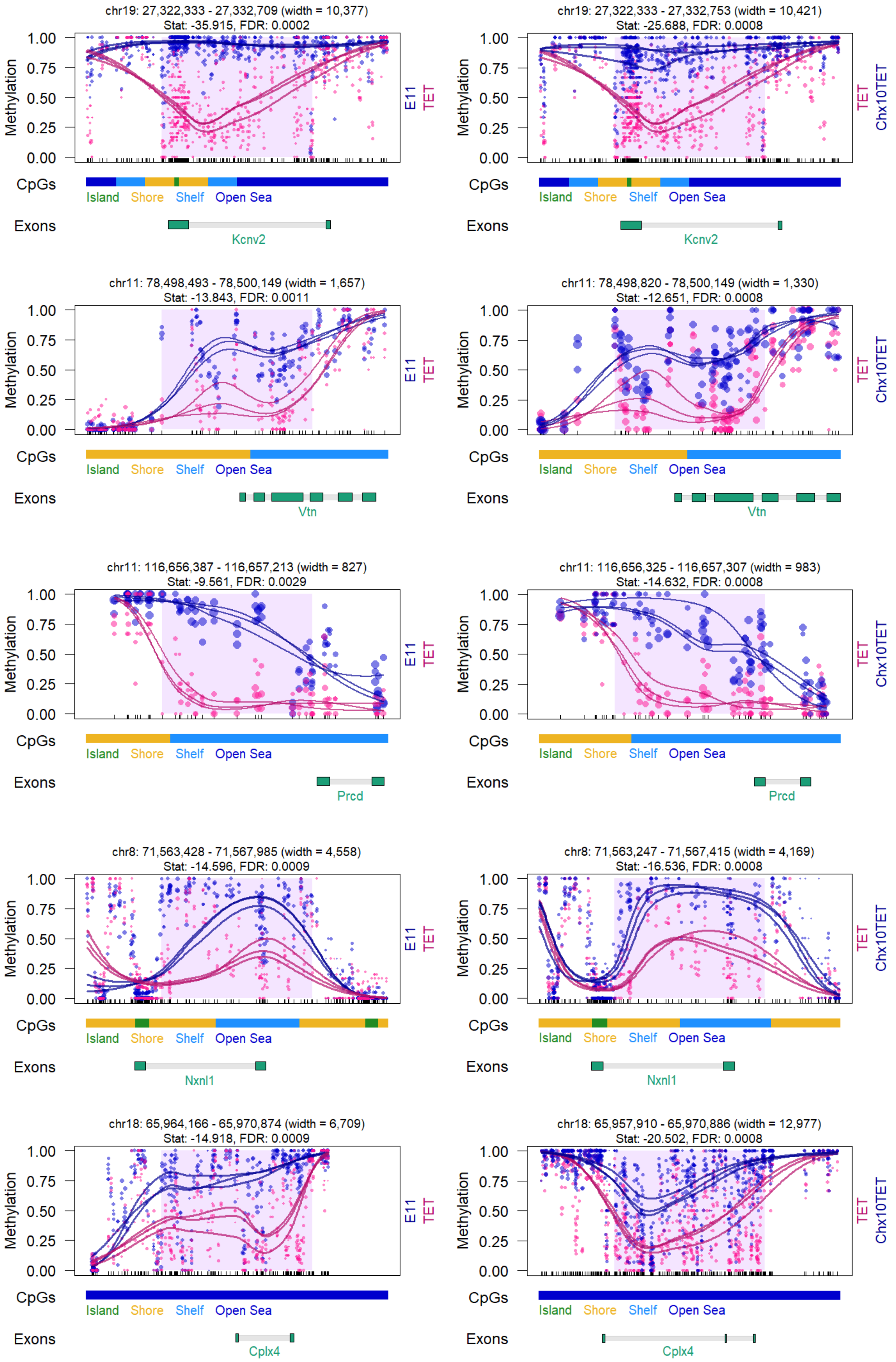

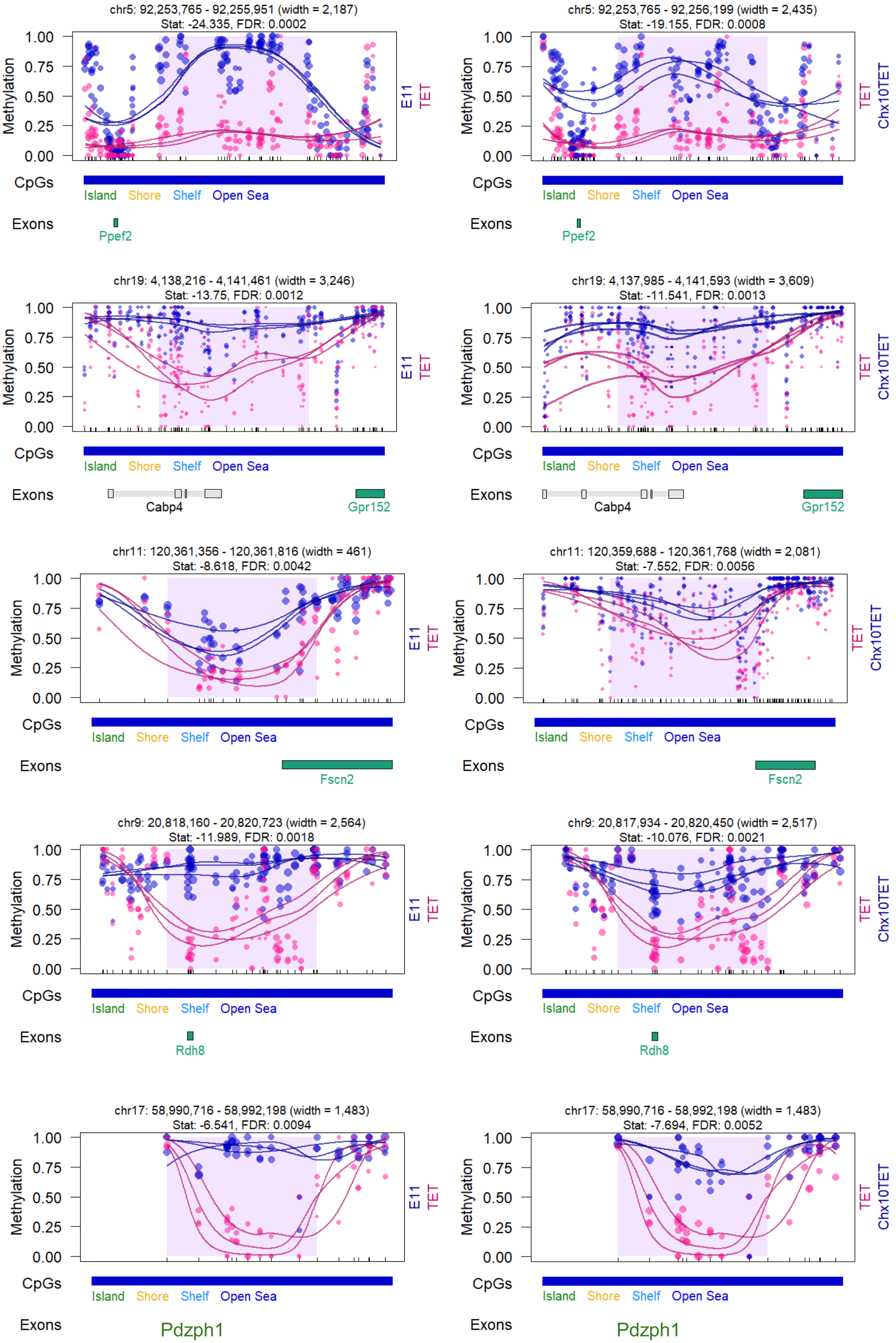

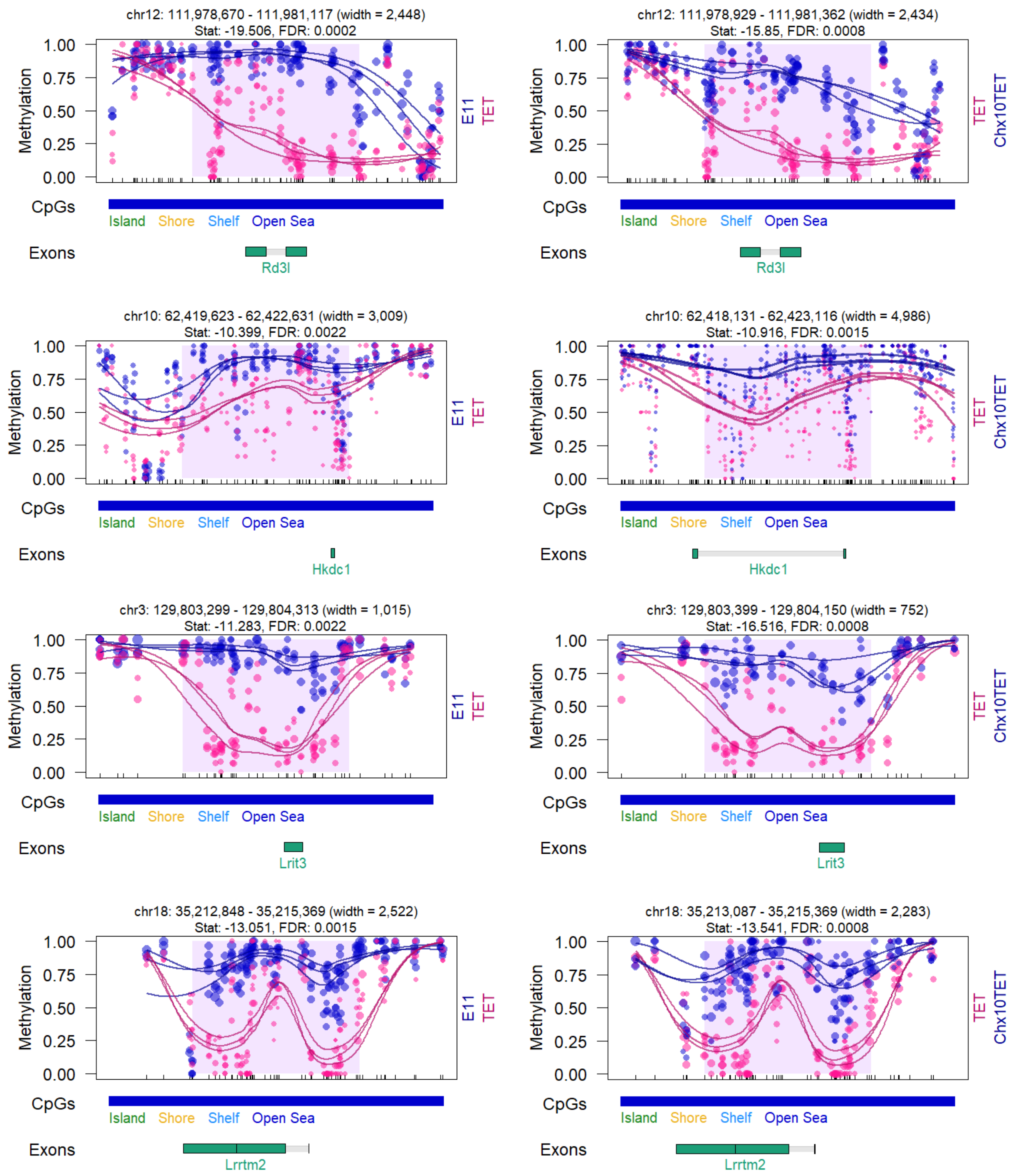

### Supplementary Figure S1

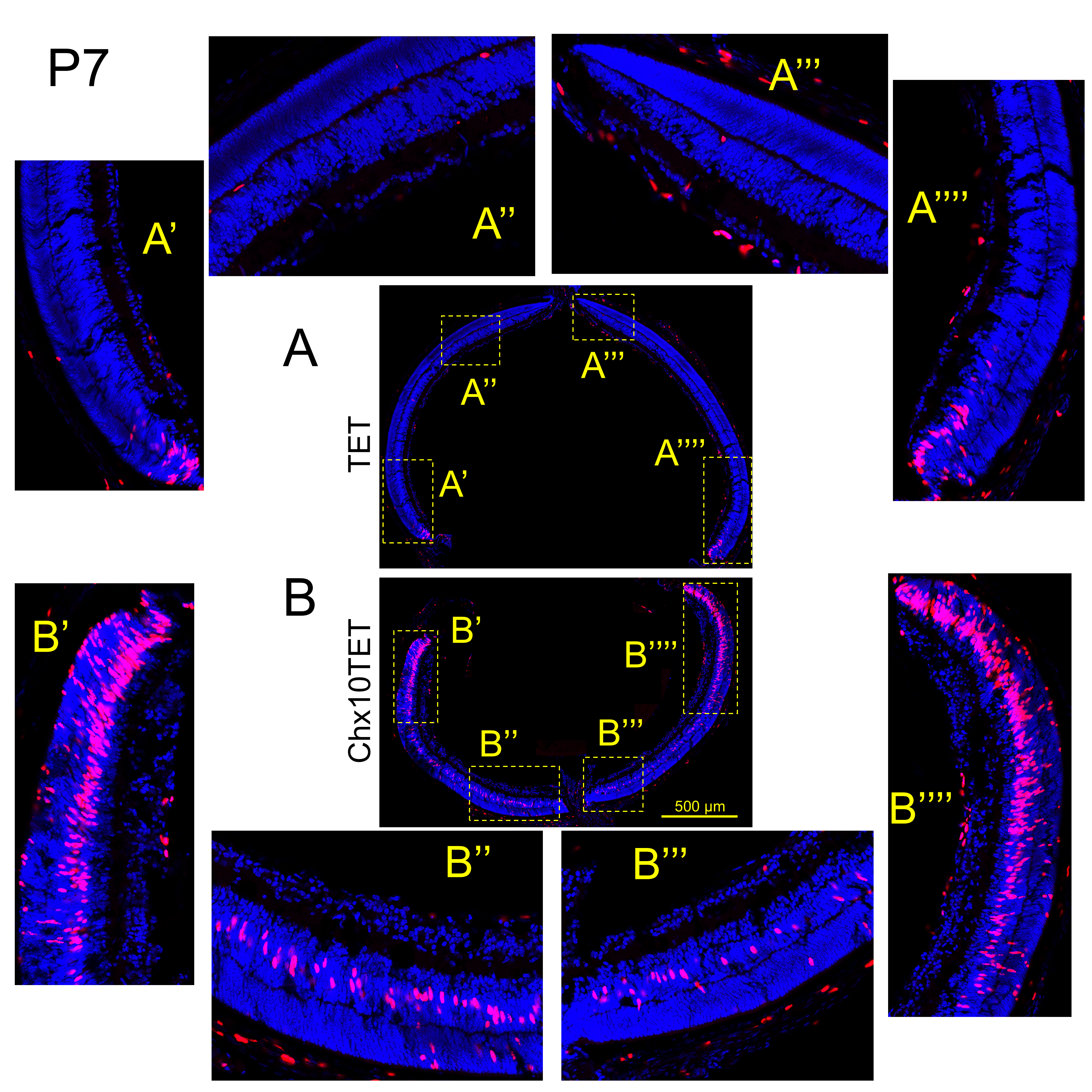

### Supplementary Figure S2

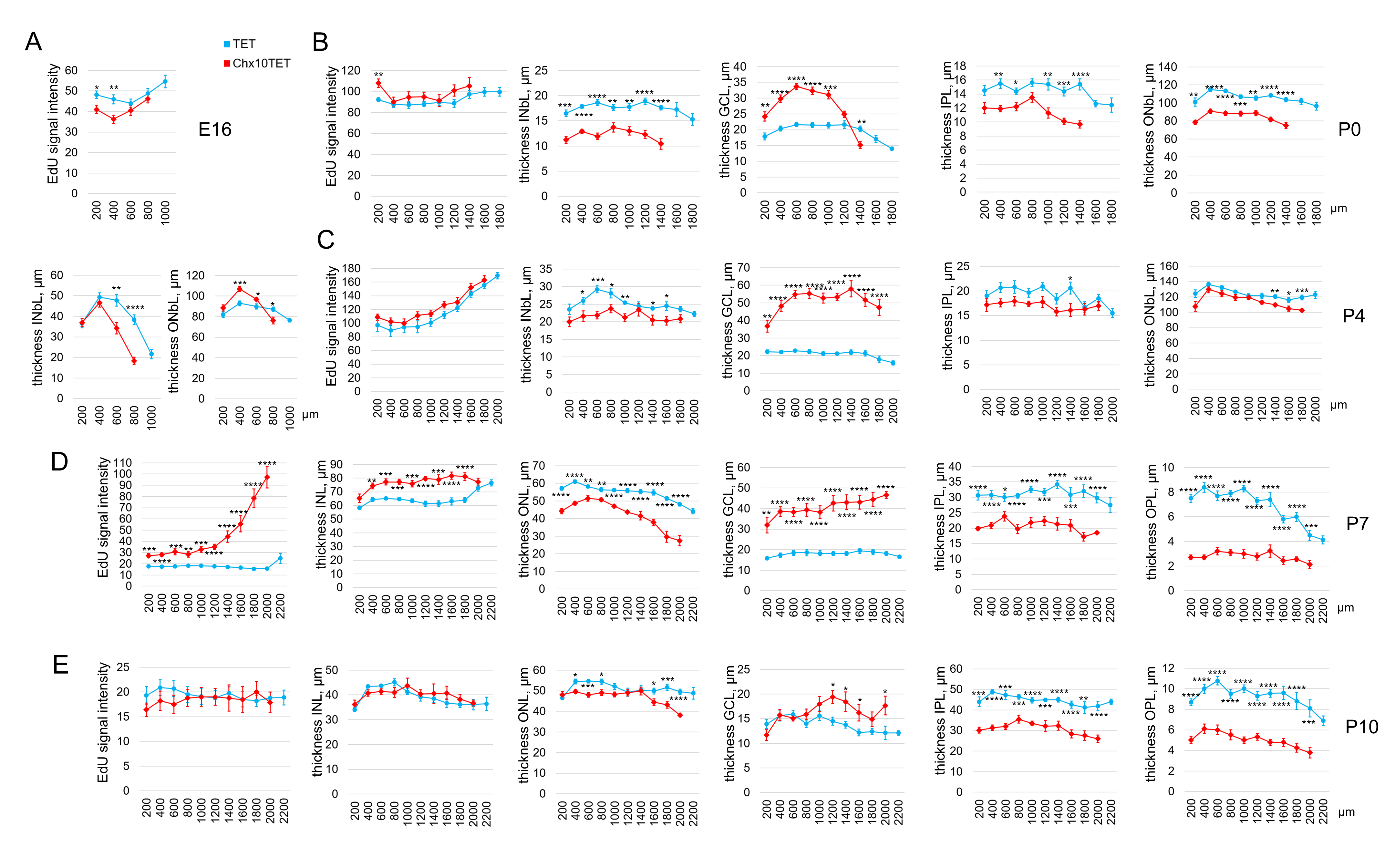

### Supplementary Figure S3

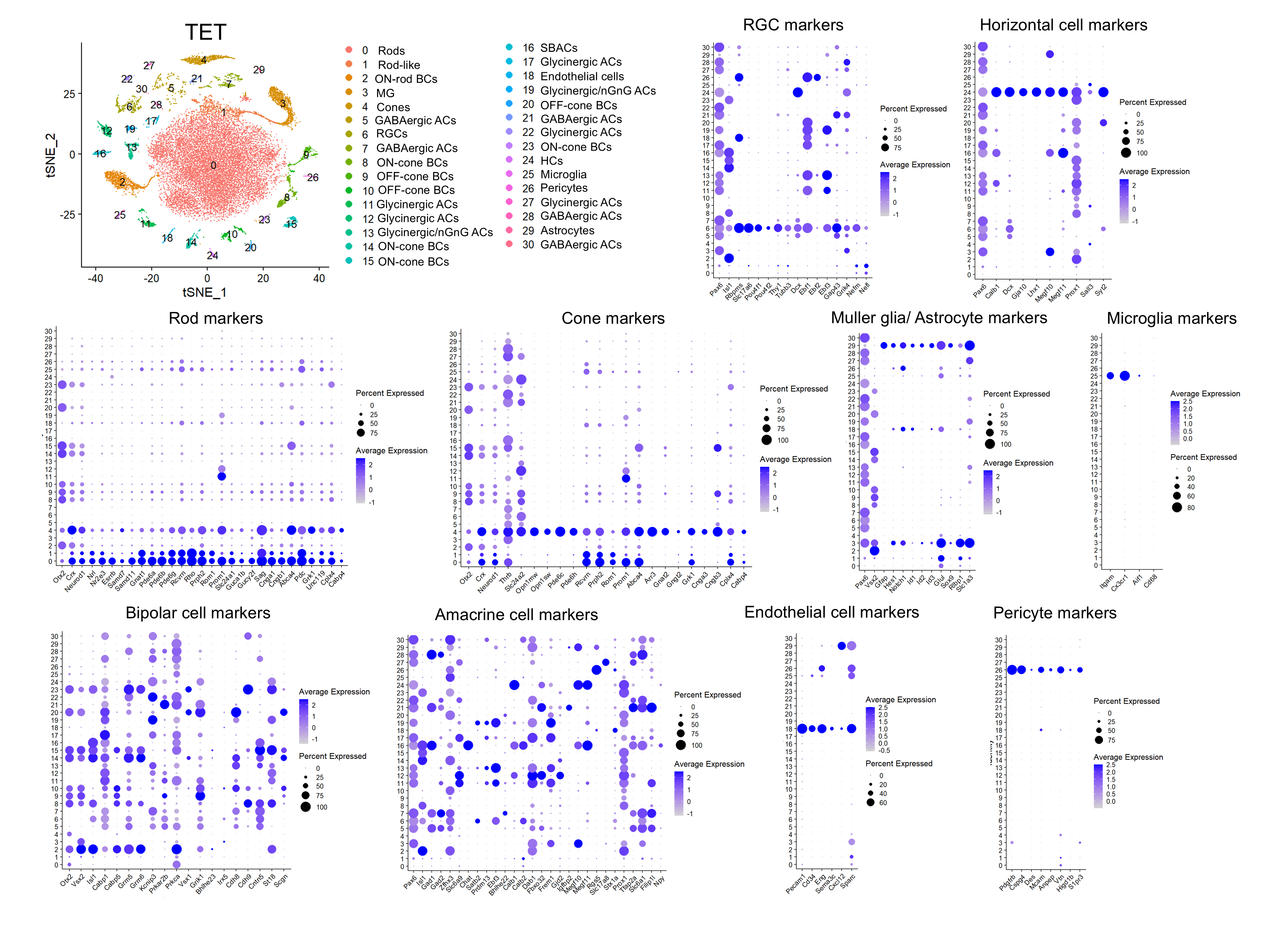

### Supplementary Figure S4

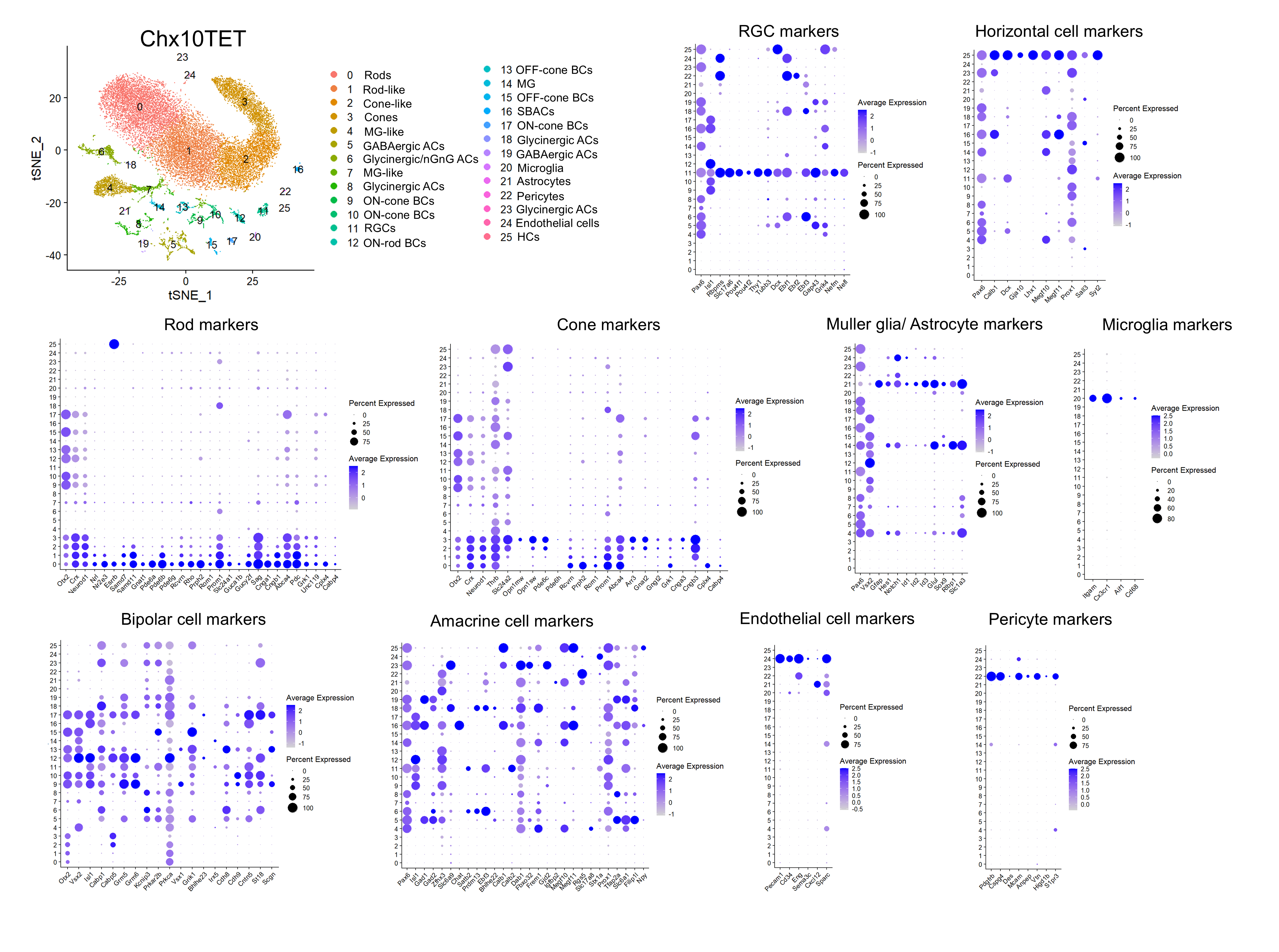

### Supplementary Figure S5

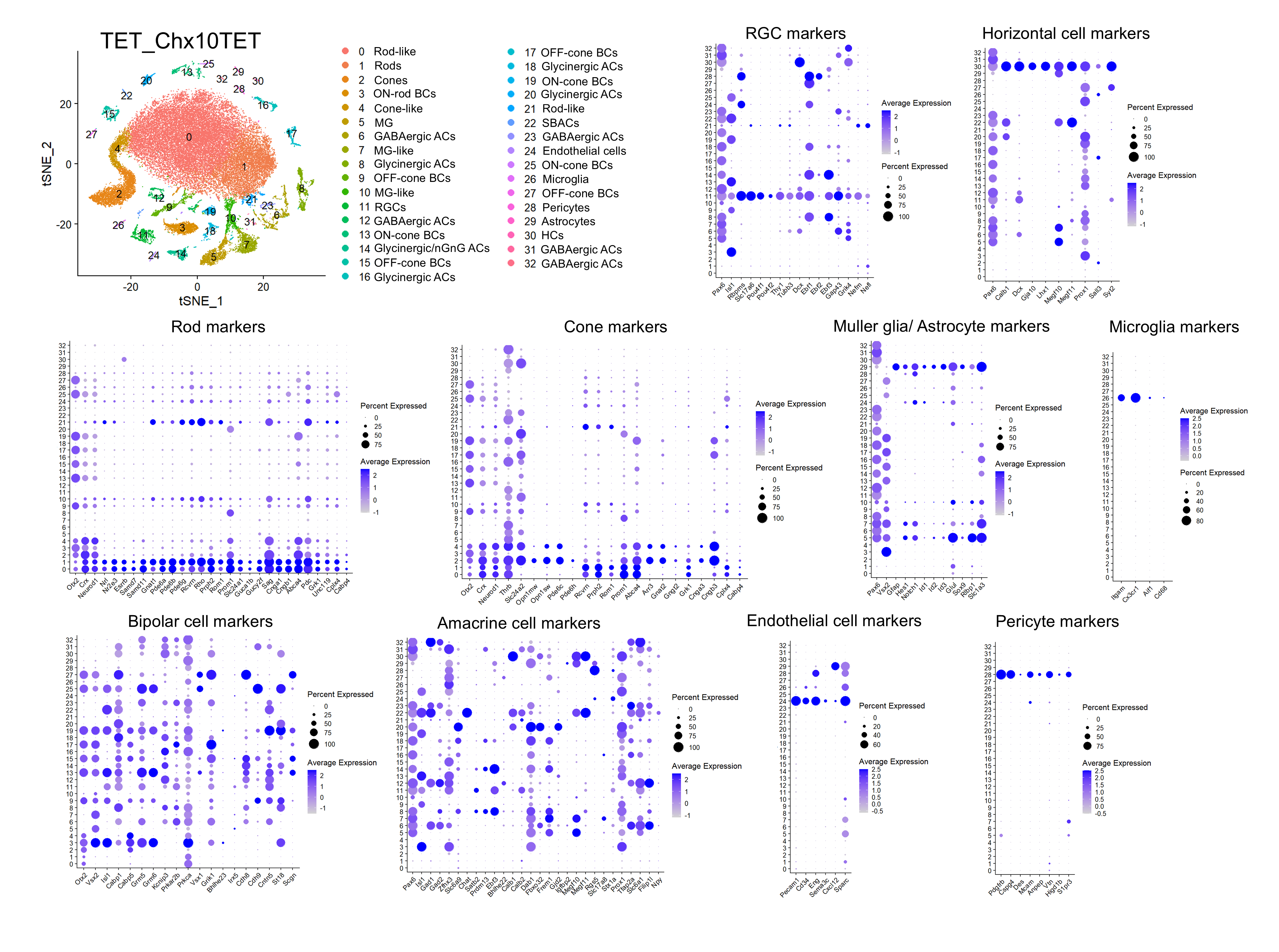

### Supplementary Figure S6

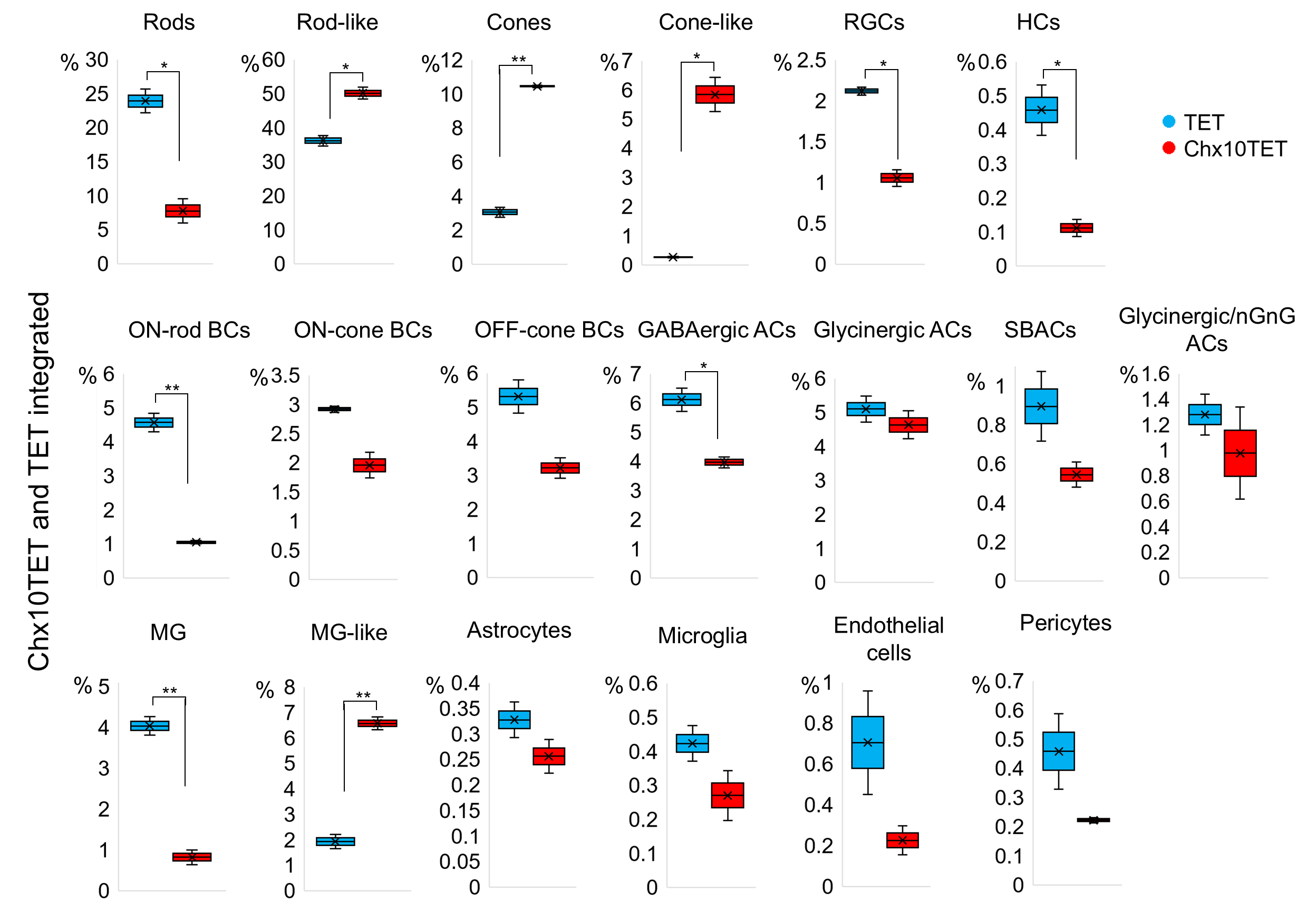

### Supplementary Figure S7

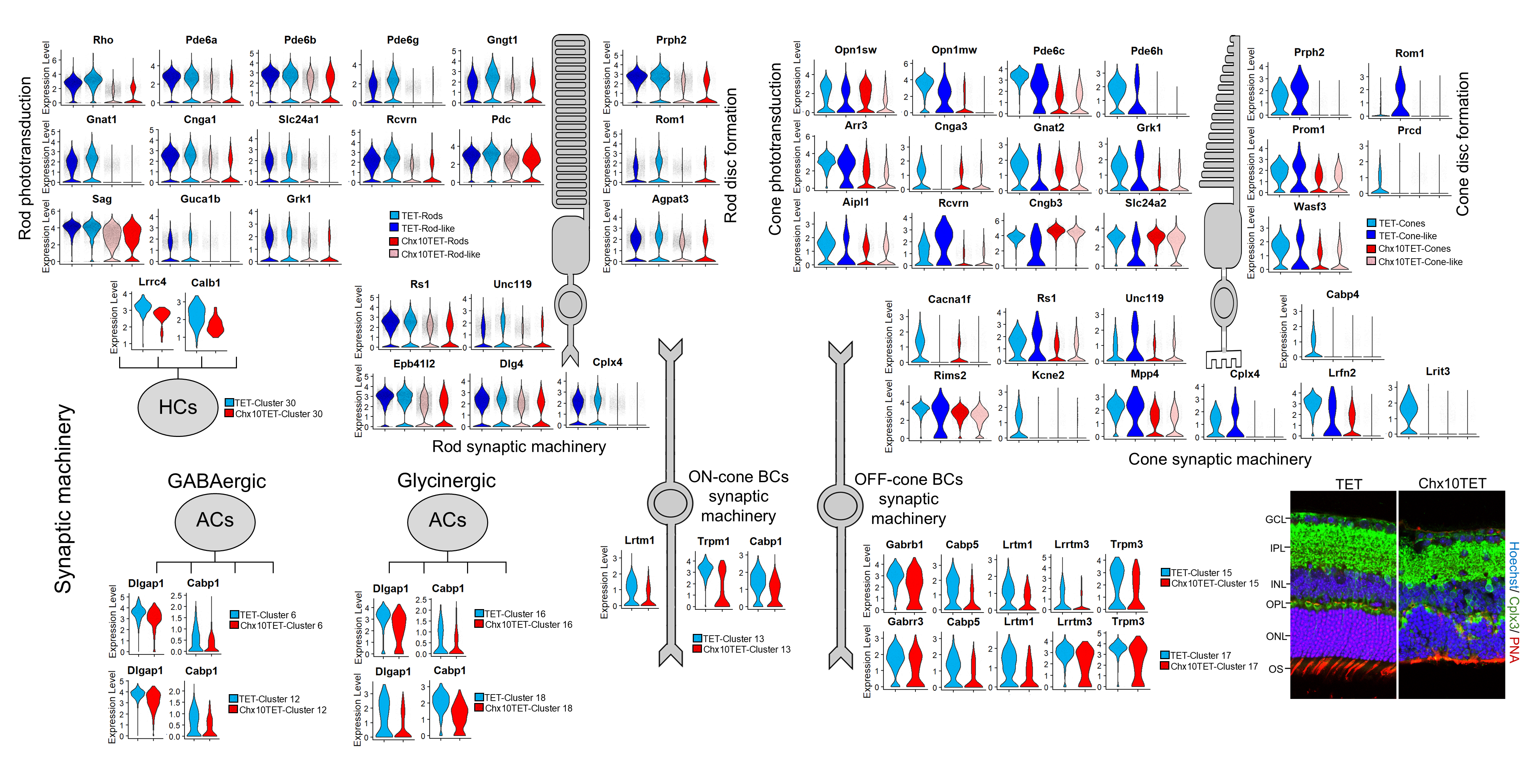
