## Supplementary Datafile S10 for "Retinal development driven by TET-dependent DNA demethylation"

### Rods

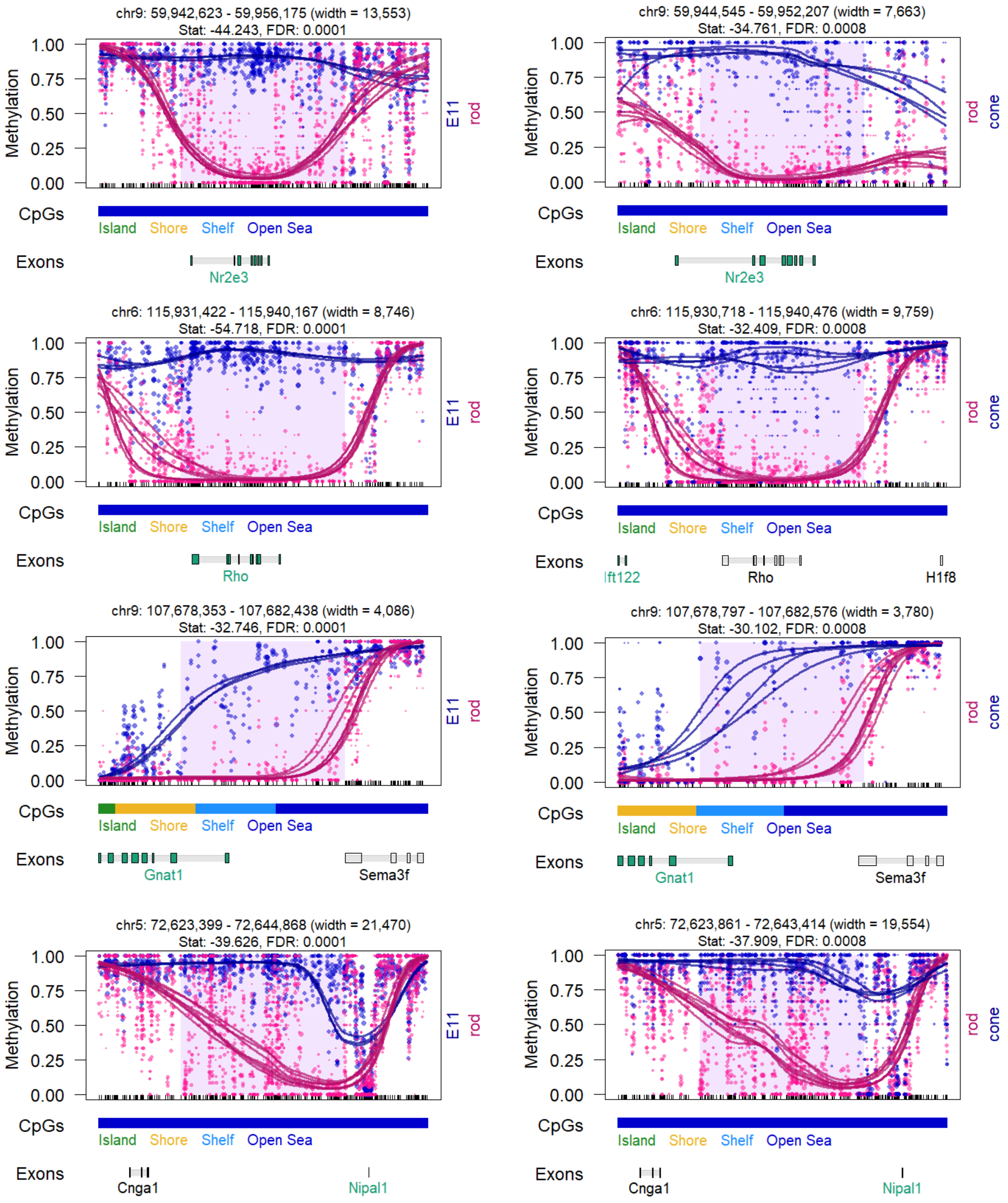

chr9: 59,942,623 - 59,956,175 (width = 13,553)
Stat: -44.243, FDR: 0.0001
chr9: 59,944,545 - 59,952,207 (width = 7,663)
Stat: -34.761, FDR: 0.0008
Nr2e3
chr6: 115,931,422 - 115,940,167 (width = 8,746)
Stat: -54.718, FDR: 0.0001
chr6: 115,930,718 - 115,940,476 (width = 9,759)
Stat: -32.409, FDR: 0.0008
Rho
Ift122
H1f8
chr9: 107,678,353 - 107,682,438 (width = 4,086)
Stat: -32.746, FDR: 0.0001
chr9: 107,678,797 - 107,682,576 (width = 3,780)
Stat: -30.102, FDR: 0.0008
Gnat1
Sema3f
chr5: 72,623,399 - 72,644,868 (width = 21,470)
Stat: -39.626, FDR: 0.0001
chr5: 72,623,861 - 72,643,414 (width = 19,554)
Stat: -37.909, FDR: 0.0008
Cnga1
Nipal1
Methylation
1.00
0.75
0.50
0.25
0.00
E11
rod
cone
CpGs
Island
Shore
Shelf
Open Sea
Exons

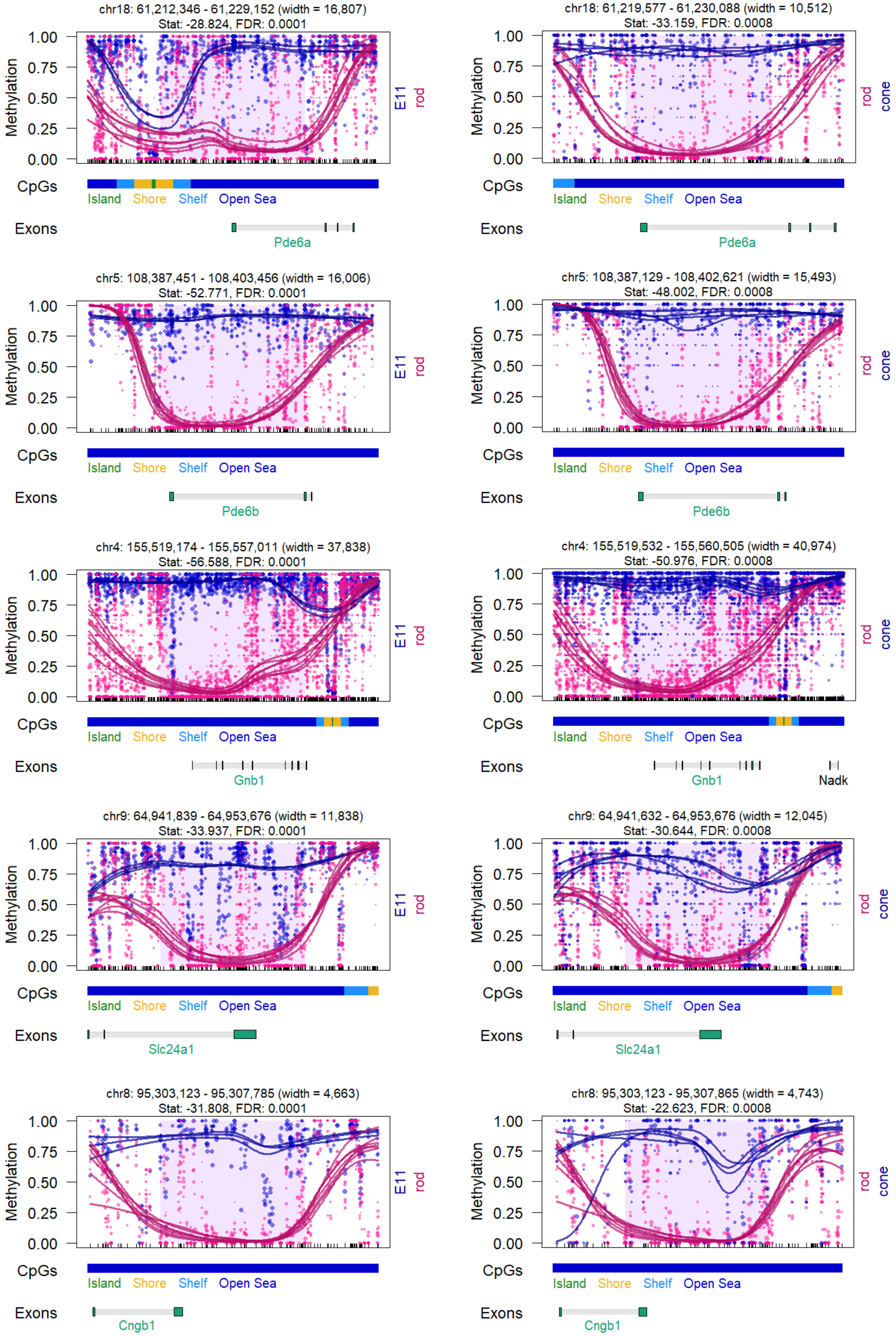

chr18: 61,212,346 - 61,229,152 (width = 16,807)
Stat: -28.824, FDR: 0.0001
chr18: 61,219,577 - 61,230,088 (width = 10,512)
Stat: -33.159, FDR: 0.0008
Pde6a
chr5: 108,387,451 - 108,403,456 (width = 16,006)
Stat: -52.771, FDR: 0.0001
chr5: 108,387,129 - 108,402,621 (width = 15,493)
Stat: -48.002, FDR: 0.0008
Pde6b
chr4: 155,519,174 - 155,557,011 (width = 37,838)
Stat: -56.588, FDR: 0.0001
chr4: 155,519,532 - 155,560,505 (width = 40,974)
Stat: -50.976, FDR: 0.0008
Gnb1
Nadk
chr9: 64,941,839 - 64,953,676 (width = 11,838)
Stat: -33.937, FDR: 0.0001
chr9: 64,941,632 - 64,953,676 (width = 12,045)
Stat: -30.644, FDR: 0.0008
Slc24a1
chr8: 95,303,123 - 95,307,785 (width = 4,663)
Stat: -31.808, FDR: 0.0001
chr8: 95,303,123 - 95,307,865 (width = 4,743)
Stat: -22.623, FDR: 0.0008
Cngb1
Methylation
1.00
0.75
0.50
0.25
0.00
E11
rod
cone
CpGs
Island
Shore
Shelf
Open Sea
Exons

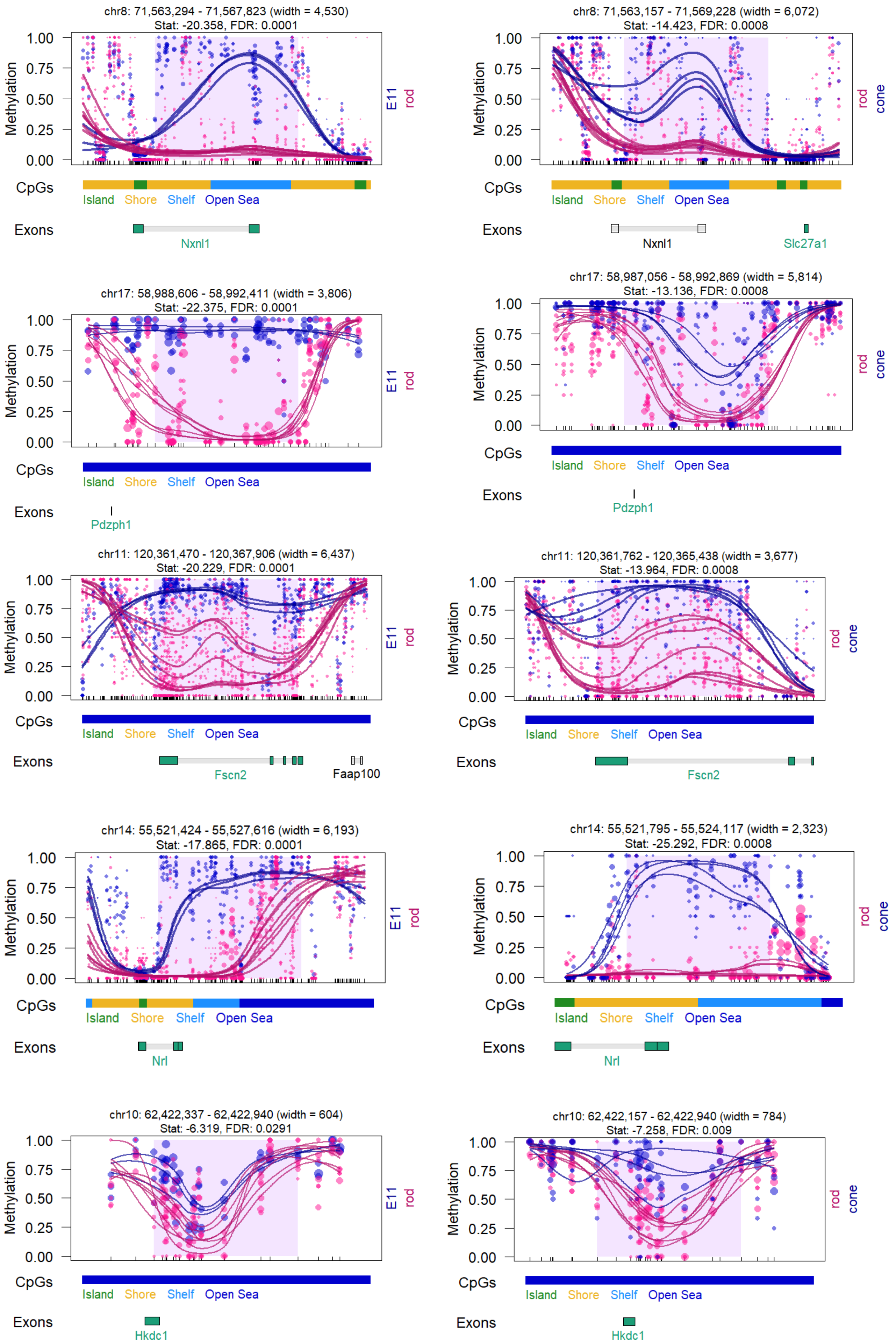

chr8: 71,563,294 - 71,567,823 (width = 4,530)
Stat: -20.358, FDR: 0.0001
Methylation
1.00
0.75
0.50
0.25
0.00
E11
rod
CpGs
Island
Shore
Shelf
Open Sea
Exons
Nxnl1
chr8: 71,563,157 - 71,569,228 (width = 6,072)
Stat: -14.423, FDR: 0.0008
rod
cone
Nxnl1
Slc27a1
chr17: 58,988,606 - 58,992,411 (width = 3,806)
Stat: -22.375, FDR: 0.0001
Pdzph1
chr17: 58,987,056 - 58,992,869 (width = 5,814)
Stat: -13.136, FDR: 0.0008
Pdzph1
chr11: 120,361,470 - 120,367,906 (width = 6,437)
Stat: -20.229, FDR: 0.0001
Fscn2
Faap100
chr11: 120,361,762 - 120,365,438 (width = 3,677)
Stat: -13.964, FDR: 0.0008
Fscn2
chr14: 55,521,424 - 55,527,616 (width = 6,193)
Stat: -17.865, FDR: 0.0001
Nrl
chr14: 55,521,795 - 55,524,117 (width = 2,323)
Stat: -25.292, FDR: 0.0008
Nrl
chr10: 62,422,337 - 62,422,940 (width = 604)
Stat: -6.319, FDR: 0.0291
Hkdc1
chr10: 62,422,157 - 62,422,940 (width = 784)
Stat: -7.258, FDR: 0.009
Hkdc1

### Cones

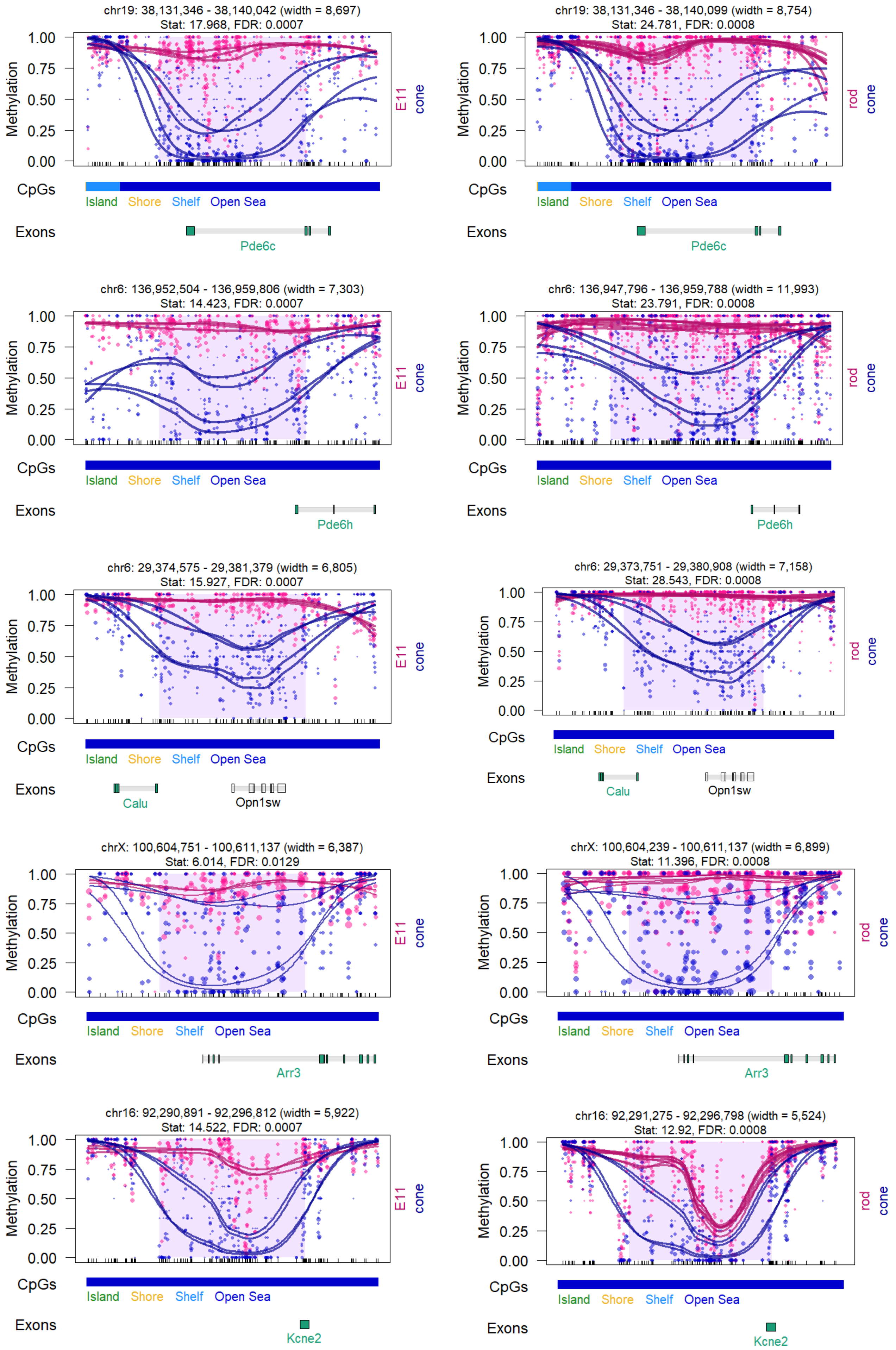
chr19: 38,131,346 - 38,140,042 (width = 8,697)
Stat: 17.968, FDR: 0.0007
chr19: 38,131,346 - 38,140,099 (width = 8,754)
Stat: 24.781, FDR: 0.0008
Methylation
E11
rod
cone
CpGs
Island Shore Shelf Open Sea
Exons
Pde6c
chr6: 136,952,504 - 136,959,806 (width = 7,303)
Stat: 14.423, FDR: 0.0007
chr6: 136,947,796 - 136,959,788 (width = 11,993)
Stat: 23.791, FDR: 0.0008
Pde6h
chr6: 29,374,575 - 29,381,379 (width = 6,805)
Stat: 15.927, FDR: 0.0007
chr6: 29,373,751 - 29,380,908 (width = 7,158)
Stat: 28.543, FDR: 0.0008
Calu
Opn1sw
chrX: 100,604,751 - 100,611,137 (width = 6,387)
Stat: 6.014, FDR: 0.0129
chrX: 100,604,239 - 100,611,137 (width = 6,899)
Stat: 11.396, FDR: 0.0008
Arr3
chr16: 92,290,891 - 92,296,812 (width = 5,922)
Stat: 14.522, FDR: 0.0007
chr16: 92,291,275 - 92,296,798 (width = 5,524)
Stat: 12.92, FDR: 0.0008
Kcne2

### Both, Rods and Cones

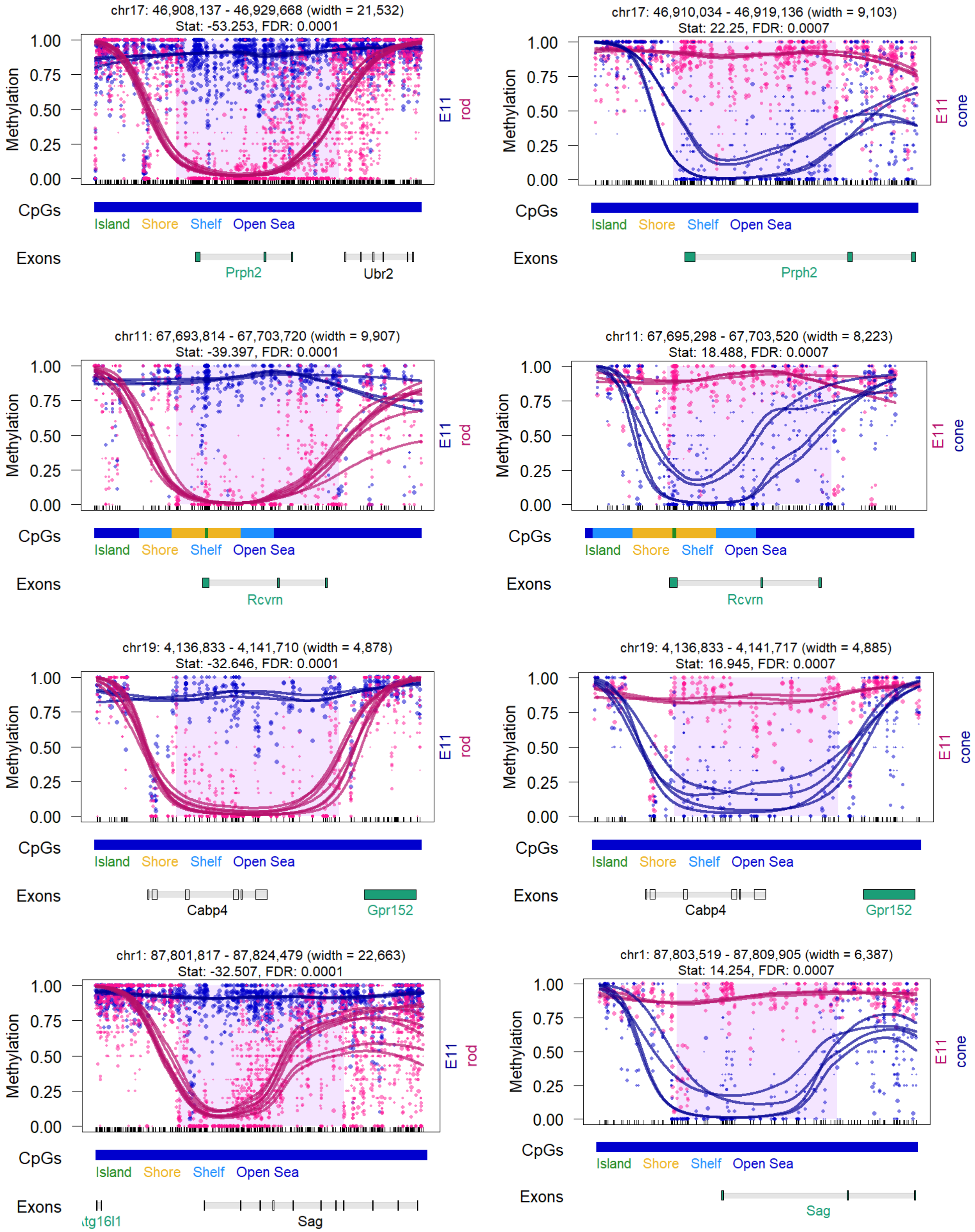

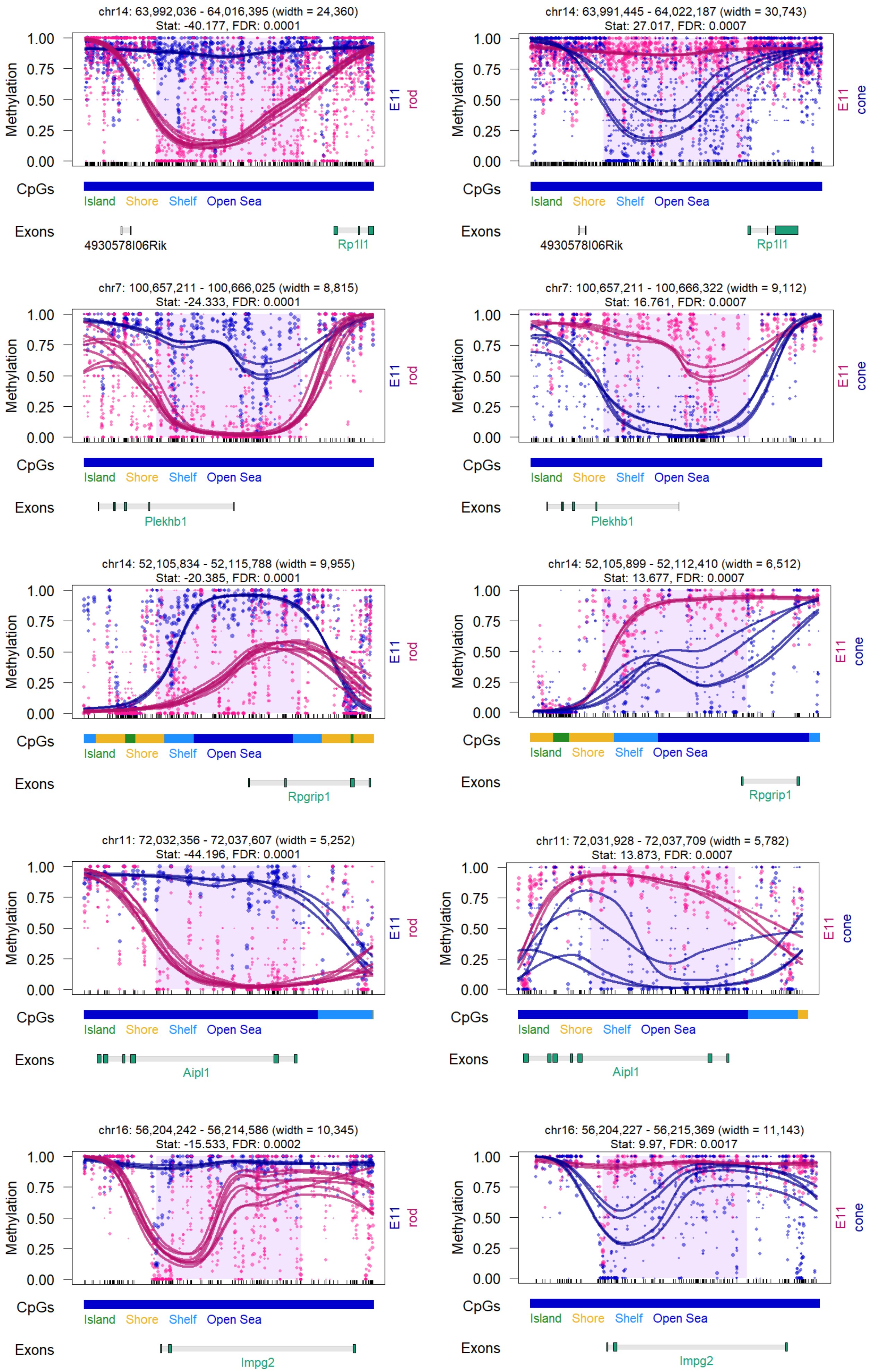
chr14: 63,992,036 - 64,016,395 (width = 24,360)
Stat: -40.177, FDR: 0.0001
Methylation
E11 rod
CpGs
Island Shore Shelf Open Sea
Exons
4930578I06Rik
Rp1l1
chr14: 63,991,445 - 64,022,187 (width = 30,743)
Stat: 27.017, FDR: 0.0007
Methylation
E11 cone
CpGs
Island Shore Shelf Open Sea
Exons
4930578I06Rik
Rp1l1
chr7: 100,657,211 - 100,666,025 (width = 8,815)
Stat: -24.333, FDR: 0.0001
Methylation
E11 rod
CpGs
Island Shore Shelf Open Sea
Exons
Plekhb1
chr7: 100,657,211 - 100,666,322 (width = 9,112)
Stat: 16.761, FDR: 0.0007
Methylation
E11 cone
CpGs
Island Shore Shelf Open Sea
Exons
Plekhb1
chr14: 52,105,834 - 52,115,788 (width = 9,955)
Stat: -20.385, FDR: 0.0001
Methylation
E11 rod
CpGs
Island Shore Shelf Open Sea
Exons
Rpgrip1
chr14: 52,105,899 - 52,112,410 (width = 6,512)
Stat: 13.677, FDR: 0.0007
Methylation
E11 cone
CpGs
Island Shore Shelf Open Sea
Exons
Rpgrip1
chr11: 72,032,356 - 72,037,607 (width = 5,252)
Stat: -44.196, FDR: 0.0001
Methylation
E11 rod
CpGs
Island Shore Shelf Open Sea
Exons
Aipl1
chr11: 72,031,928 - 72,037,709 (width = 5,782)
Stat: 13.873, FDR: 0.0007
Methylation
E11 cone
CpGs
Island Shore Shelf Open Sea
Exons
Aipl1
chr16: 56,204,242 - 56,214,586 (width = 10,345)
Stat: -15.533, FDR: 0.0002
Methylation
E11 rod
CpGs
Island Shore Shelf Open Sea
Exons
Impg2
chr16: 56,204,227 - 56,215,369 (width = 11,143)
Stat: 9.97, FDR: 0.0017
Methylation
E11 cone
CpGs
Island Shore Shelf Open Sea
Exons
Impg2

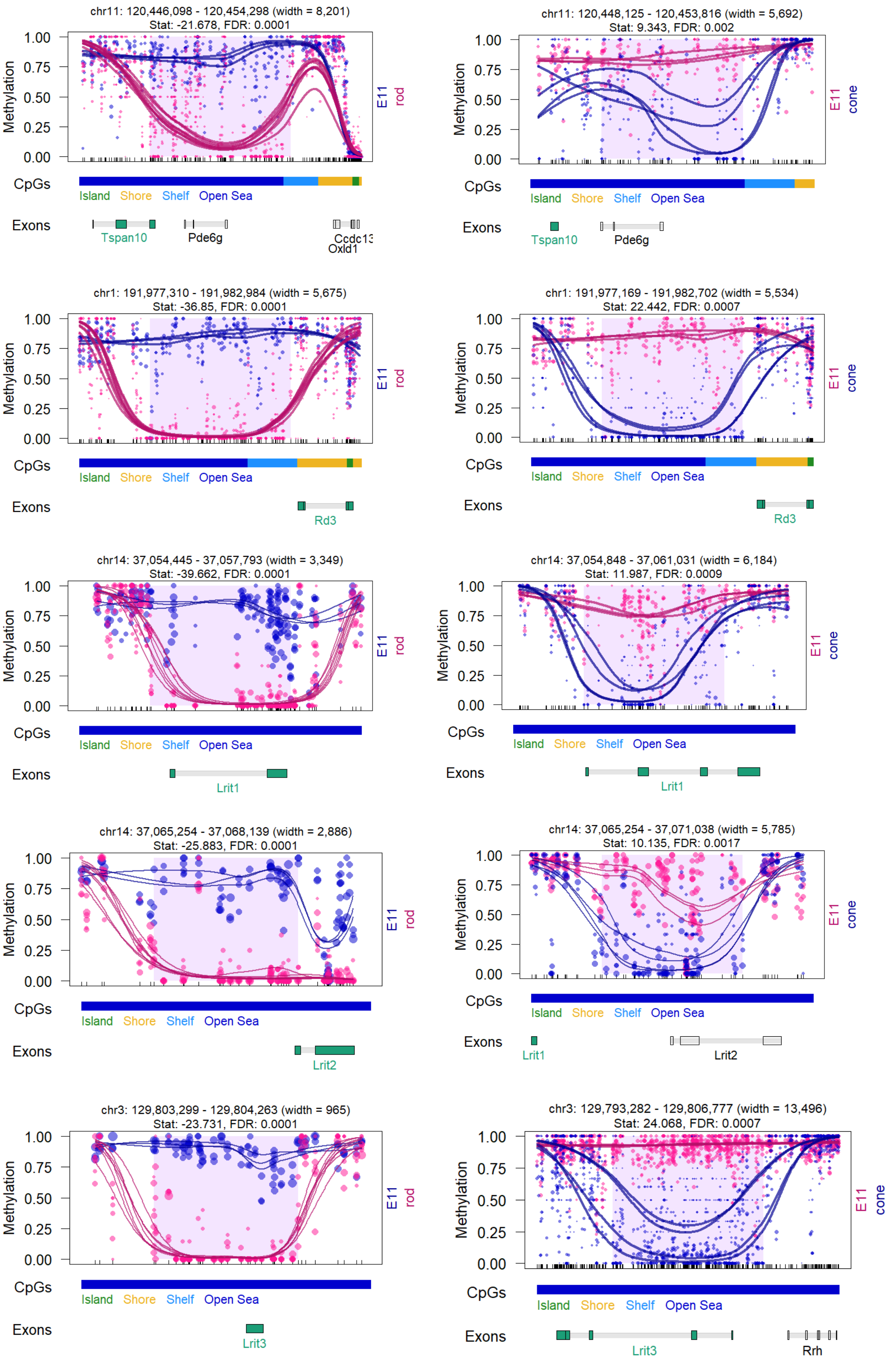
chr11: 120,446,098 - 120,454,298 (width = 8,201)
Stat: -21.678, FDR: 0.0001
Methylation
E11 rod
CpGs
Island Shore Shelf Open Sea
Exons
Tspan10
Pde6g
Ccdc13
Oxld1
chr11: 120,448,125 - 120,453,816 (width = 5,692)
Stat: 9.343, FDR: 0.002
E11 cone
Tspan10
Pde6g
chr1: 191,977,310 - 191,982,984 (width = 5,675)
Stat: -36.85, FDR: 0.0001
Rd3
chr1: 191,977,169 - 191,982,702 (width = 5,534)
Stat: 22.442, FDR: 0.0007
Rd3
chr14: 37,054,445 - 37,057,793 (width = 3,349)
Stat: -39.662, FDR: 0.0001
Lrit1
chr14: 37,054,848 - 37,061,031 (width = 6,184)
Stat: 11.987, FDR: 0.0009
Lrit1
chr14: 37,065,254 - 37,068,139 (width = 2,886)
Stat: -25.883, FDR: 0.0001
Lrit2
chr14: 37,065,254 - 37,071,038 (width = 5,785)
Stat: 10.135, FDR: 0.0017
Lrit1
Lrit2
chr3: 129,803,299 - 129,804,263 (width = 965)
Stat: -23.731, FDR: 0.0001
Lrit3
chr3: 129,793,282 - 129,806,777 (width = 13,496)
Stat: 24.068, FDR: 0.0007
Lrit3
Rrh

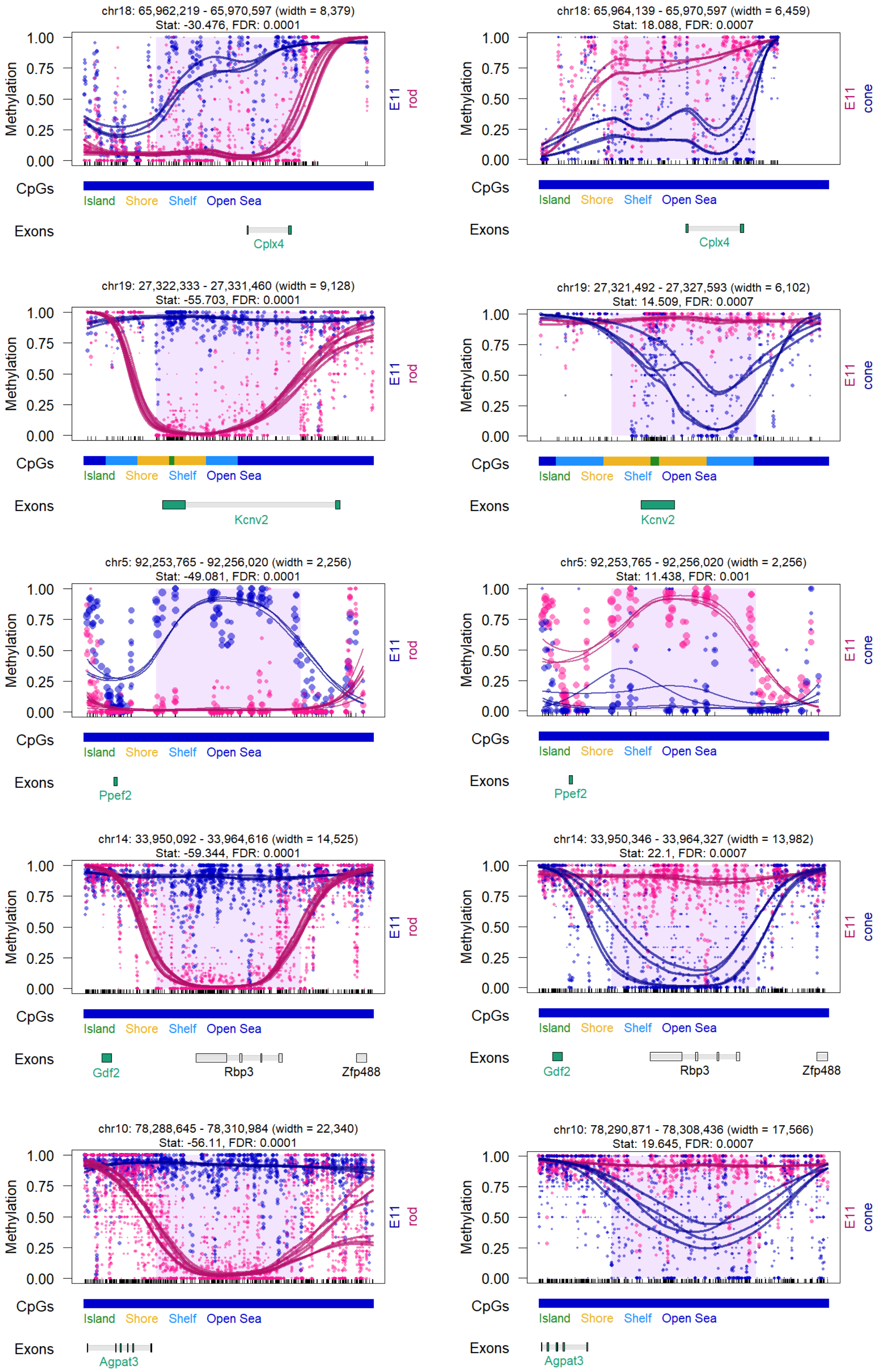
chr18: 65,962,219 - 65,970,597 (width = 8,379)
Stat: -30.476, FDR: 0.0001
Methylation
1.00
0.75
0.50
0.25
0.00
E11
rod
CpGs
Island
Shore
Shelf
Open Sea
Exons
Cplx4
chr18: 65,964,139 - 65,970,597 (width = 6,459)
Stat: 18.088, FDR: 0.0007
E11
cone
Cplx4
chr19: 27,322,333 - 27,331,460 (width = 9,128)
Stat: -55.703, FDR: 0.0001
Kcnv2
chr19: 27,321,492 - 27,327,593 (width = 6,102)
Stat: 14.509, FDR: 0.0007
Kcnv2
chr5: 92,253,765 - 92,256,020 (width = 2,256)
Stat: -49.081, FDR: 0.0001
Ppef2
chr5: 92,253,765 - 92,256,020 (width = 2,256)
Stat: 11.438, FDR: 0.001
Ppef2
chr14: 33,950,092 - 33,964,616 (width = 14,525)
Stat: -59.344, FDR: 0.0001
Gdf2
Rbp3
Zfp488
chr14: 33,950,346 - 33,964,327 (width = 13,982)
Stat: 22.1, FDR: 0.0007
Gdf2
Rbp3
Zfp488
chr10: 78,288,645 - 78,310,984 (width = 22,340)
Stat: -56.11, FDR: 0.0001
Agpat3
chr10: 78,290,871 - 78,308,436 (width = 17,566)
Stat: 19.645, FDR: 0.0007
Agpat3

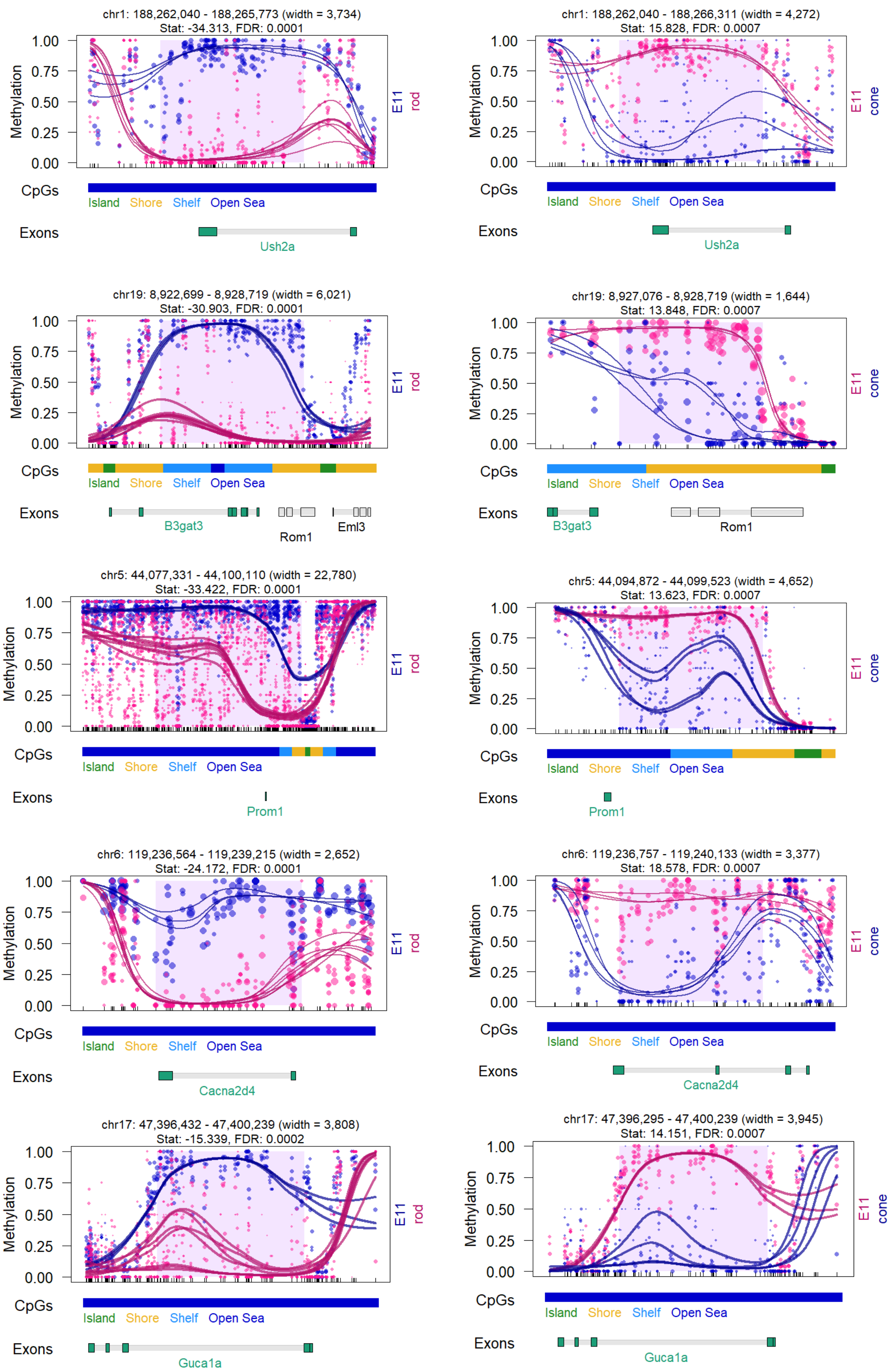

chr1: 188,262,040 - 188,265,773 (width = 3,734)
Stat: -34.313, FDR: 0.0001
Methylation
E11
rod
CpGs
Island
Shore
Shelf
Open Sea
Exons
Ush2a
chr1: 188,262,040 - 188,266,311 (width = 4,272)
Stat: 15.828, FDR: 0.0007
Methylation
E11
cone
CpGs
Island
Shore
Shelf
Open Sea
Exons
Ush2a
chr19: 8,922,699 - 8,928,719 (width = 6,021)
Stat: -30.903, FDR: 0.0001
Methylation
E11
rod
CpGs
Island
Shore
Shelf
Open Sea
Exons
B3gat3
Rom1
Eml3
chr19: 8,927,076 - 8,928,719 (width = 1,644)
Stat: 13.848, FDR: 0.0007
Methylation
E11
cone
CpGs
Island
Shore
Shelf
Open Sea
Exons
B3gat3
Rom1
chr5: 44,077,331 - 44,100,110 (width = 22,780)
Stat: -33.422, FDR: 0.0001
Methylation
E11
rod
CpGs
Island
Shore
Shelf
Open Sea
Exons
Prom1
chr5: 44,094,872 - 44,099,523 (width = 4,652)
Stat: 13.623, FDR: 0.0007
Methylation
E11
cone
CpGs
Island
Shore
Shelf
Open Sea
Exons
Prom1
chr6: 119,236,564 - 119,239,215 (width = 2,652)
Stat: -24.172, FDR: 0.0001
Methylation
E11
rod
CpGs
Island
Shore
Shelf
Open Sea
Exons
Cacna2d4
chr6: 119,236,757 - 119,240,133 (width = 3,377)
Stat: 18.578, FDR: 0.0007
Methylation
E11
cone
CpGs
Island
Shore
Shelf
Open Sea
Exons
Cacna2d4
chr17: 47,396,432 - 47,400,239 (width = 3,808)
Stat: -15.339, FDR: 0.0002
Methylation
E11
rod
CpGs
Island
Shore
Shelf
Open Sea
Exons
Guca1a
chr17: 47,396,295 - 47,400,239 (width = 3,945)
Stat: 14.151, FDR: 0.0007
Methylation
E11
cone
CpGs
Island
Shore
Shelf
Open Sea
Exons
Guca1a

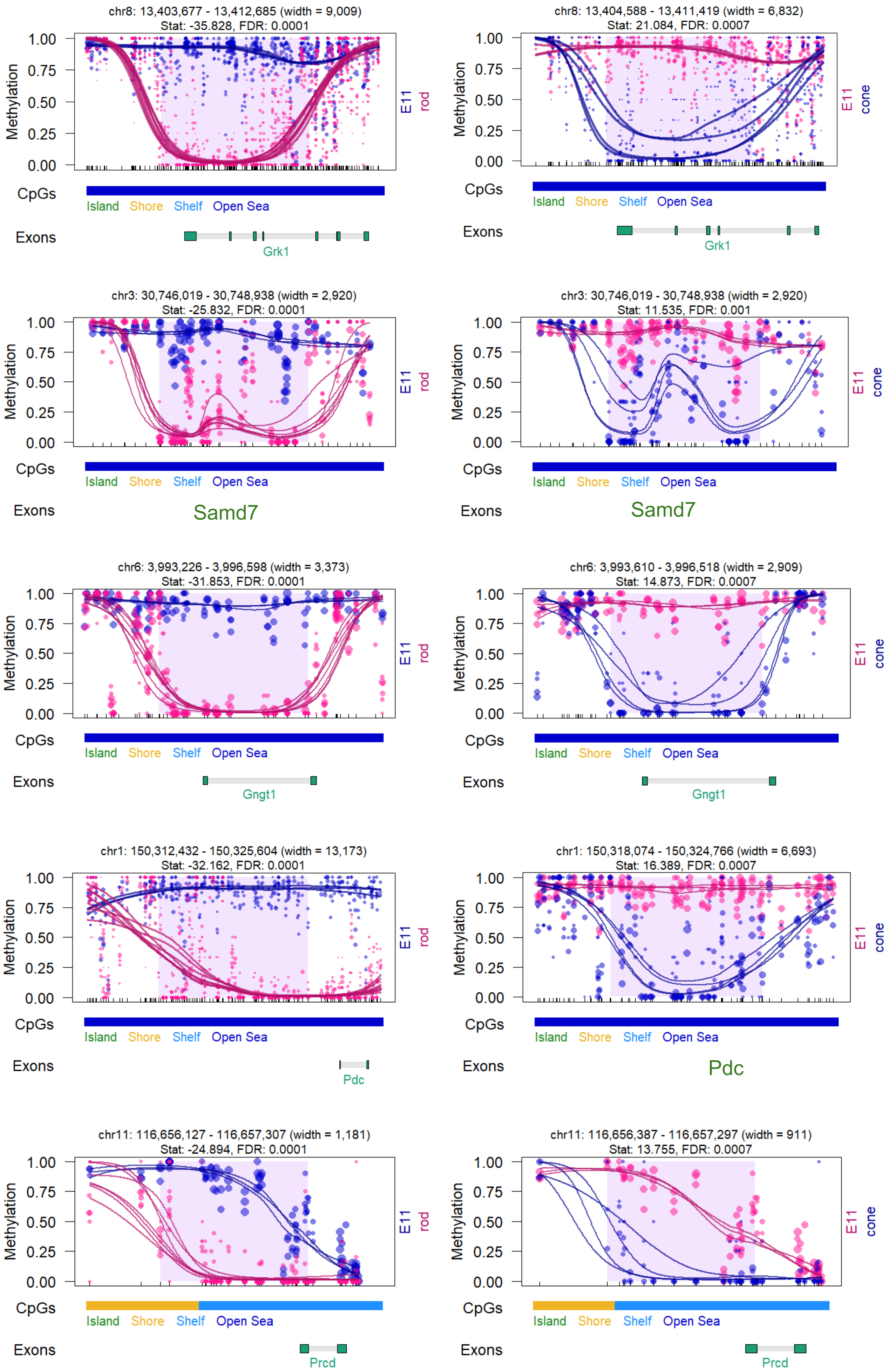

chr8: 13,403,677 - 13,412,685 (width = 9,009)
Stat: -35.828, FDR: 0.0001
Methylation
1.00
0.75
0.50
0.25
0.00
E11
rod
CpGs
Island Shore Shelf Open Sea
Exons
Grk1
chr8: 13,404,588 - 13,411,419 (width = 6,832)
Stat: 21.084, FDR: 0.0007
E11
cone
Grk1
chr3: 30,746,019 - 30,748,938 (width = 2,920)
Stat: -25.832, FDR: 0.0001
Samd7
chr3: 30,746,019 - 30,748,938 (width = 2,920)
Stat: 11.535, FDR: 0.001
Samd7
chr6: 3,993,226 - 3,996,598 (width = 3,373)
Stat: -31.853, FDR: 0.0001
Gngt1
chr6: 3,993,610 - 3,996,518 (width = 2,909)
Stat: 14.873, FDR: 0.0007
Gngt1
chr1: 150,312,432 - 150,325,604 (width = 13,173)
Stat: -32.162, FDR: 0.0001
Pdc
chr1: 150,318,074 - 150,324,766 (width = 6,693)
Stat: 16.389, FDR: 0.0007
Pdc
chr11: 116,656,127 - 116,657,307 (width = 1,181)
Stat: -24.894, FDR: 0.0001
Prcd
chr11: 116,656,387 - 116,657,297 (width = 911)
Stat: 13.755, FDR: 0.0007
Prcd

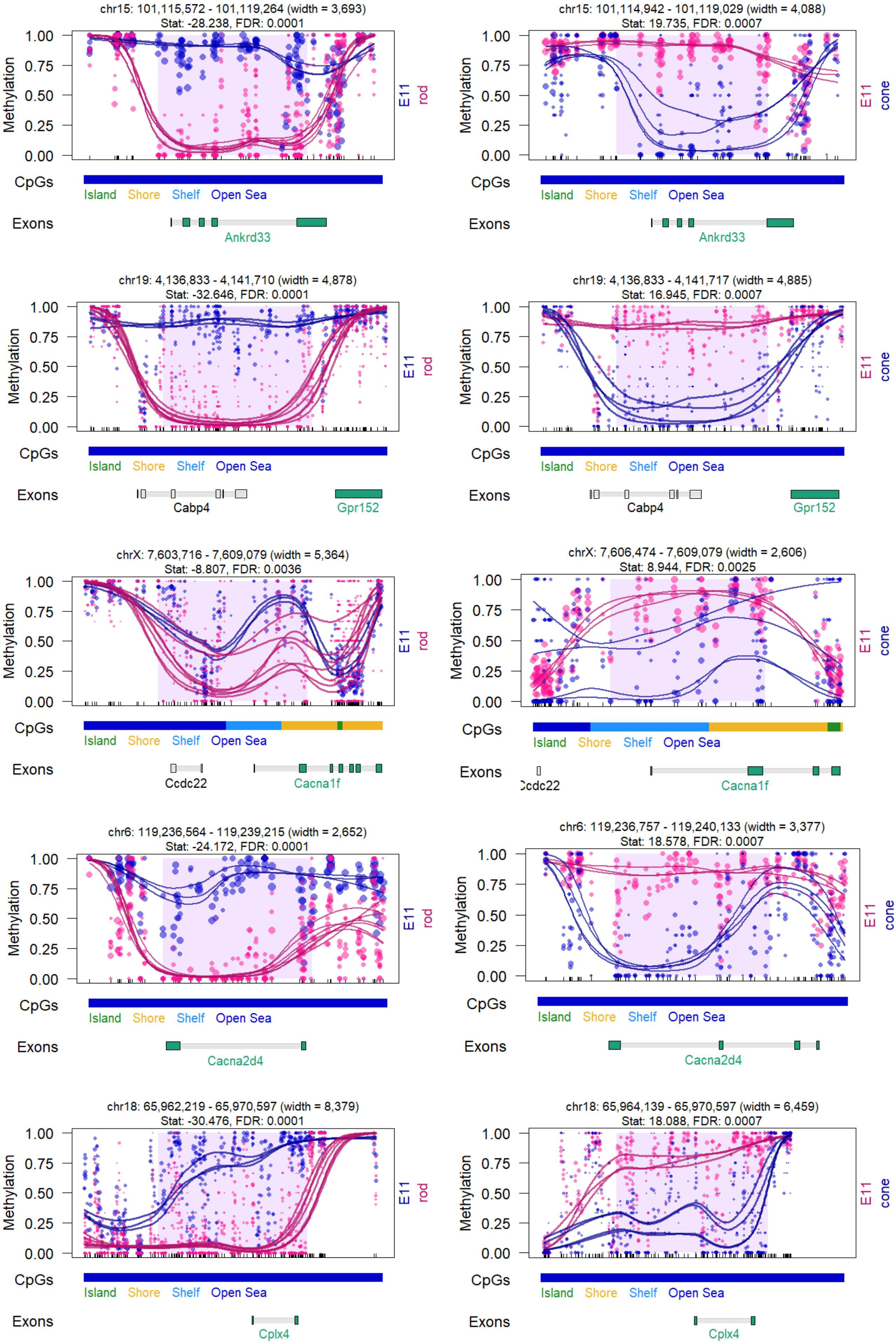
chr15: 101,115,572 - 101,119,264 (width = 3,693)
Stat: -28.238, FDR: 0.0001
Methylation
1.00
0.75
0.50
0.25
0.00
E11
rod
CpGs
Island
Shore
Shelf
Open Sea
Exons
Ankrd33
chr15: 101,114,942 - 101,119,029 (width = 4,088)
Stat: 19.735, FDR: 0.0007
E11
cone
Ankrd33
chr19: 4,136,833 - 4,141,710 (width = 4,878)
Stat: -32.646, FDR: 0.0001
Cabp4
Gpr152
chr19: 4,136,833 - 4,141,717 (width = 4,885)
Stat: 16.945, FDR: 0.0007
Cabp4
Gpr152
chrX: 7,603,716 - 7,609,079 (width = 5,364)
Stat: -8.807, FDR: 0.0036
Ccdc22
Cacna1f
chrX: 7,606,474 - 7,609,079 (width = 2,606)
Stat: 8.944, FDR: 0.0025
Cacna1f
chr6: 119,236,564 - 119,239,215 (width = 2,652)
Stat: -24.172, FDR: 0.0001
Cacna2d4
chr6: 119,236,757 - 119,240,133 (width = 3,377)
Stat: 18.578, FDR: 0.0007
Cacna2d4
chr18: 65,962,219 - 65,970,597 (width = 8,379)
Stat: -30.476, FDR: 0.0001
Cplx4
chr18: 65,964,139 - 65,970,597 (width = 6,459)
Stat: 18.088, FDR: 0.0007
Cplx4

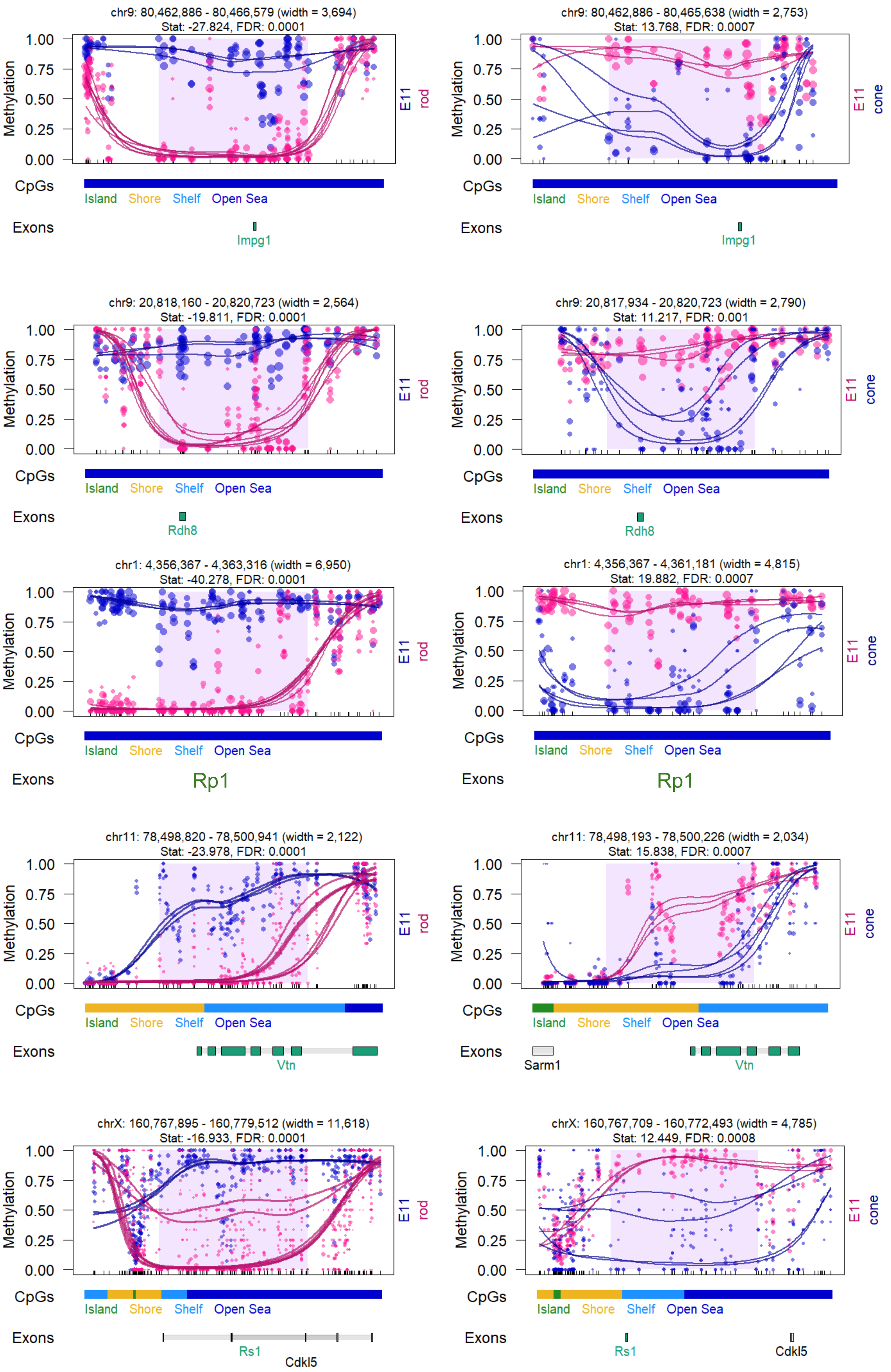
chr9: 80,462,886 - 80,466,579 (width = 3,694)
Stat: -27.824, FDR: 0.0001
Methylation
1.00
0.75
0.50
0.25
0.00
E11
rod
CpGs
Island Shore Shelf Open Sea
Exons
Impg1
chr9: 80,462,886 - 80,465,638 (width = 2,753)
Stat: 13.768, FDR: 0.0007
E11
cone
Impg1
chr9: 20,818,160 - 20,820,723 (width = 2,564)
Stat: -19.811, FDR: 0.0001
Rdh8
chr9: 20,817,934 - 20,820,723 (width = 2,790)
Stat: 11.217, FDR: 0.001
Rdh8
chr1: 4,356,367 - 4,363,316 (width = 6,950)
Stat: -40.278, FDR: 0.0001
Rp1
chr1: 4,356,367 - 4,361,181 (width = 4,815)
Stat: 19.882, FDR: 0.0007
Rp1
chr11: 78,498,820 - 78,500,941 (width = 2,122)
Stat: -23.978, FDR: 0.0001
Vtn
chr11: 78,498,193 - 78,500,226 (width = 2,034)
Stat: 15.838, FDR: 0.0007
Sarm1
Vtn
chrX: 160,767,895 - 160,779,512 (width = 11,618)
Stat: -16.933, FDR: 0.0001
Rs1
Cdkl5
chrX: 160,767,709 - 160,772,493 (width = 4,785)
Stat: 12.449, FDR: 0.0008
Rs1
Cdkl5
