## Supplementary Datafile S12 for "Retinal development driven by TET-dependent DNA demethylation"

**Supplementary Datafile S12/ R code**

A) Analysis of data obtained for control (TET) retinas

library(Seurat)

library(ggplot2)

library(gridExtra)

library(tidyr)

library(BPCells)

set.seed(123)

suppressPackageStartupMessages(library(scDblFinder))

library(SoupX)

nonExpressedGeneList = list(rods = c("Sag", "Pdc", "Rp1"), mg = c("Glul", "Rlbp1", "Slc1a3"))

obj_hdf5_D1 = load10X("/media/dmitry/d/scRNAseq_TET/D1/outs/")

head(obj_hdf5_D1$soupProfile[order(obj_hdf5_D1$soupProfile$est, decreasing = TRUE), ], n = 20)

useToEst_D1 = estimateNonExpressingCells(obj_hdf5_D1, nonExpressedGeneList = nonExpressedGeneList)

obj_hdf5_D1 = calculateContaminationFraction(obj_hdf5_D1, nonExpressedGeneList, useToEst = useToEst_D1)

head(obj_hdf5_D1$metaData)

sceD1 = adjustCounts(obj_hdf5_D1)

contr_TET1 <- CreateSeuratObject(counts = sceD1, min.cells = 3, min.features = 200, project = "TET1")

contr_TET1$mitoPercent <- PercentageFeatureSet(contr_TET1, pattern='^mt-')

contr_TET1$riboPercent <- PercentageFeatureSet(contr_TET1, pattern='^Rp[sl]')

contr_TET1_filtered <- subset(contr_TET1, subset = nCount_RNA > 500 & nFeature_RNA > 500 & mitoPercent < 5 & riboPercent < 5)

table(contr_TET1_filtered$orig.ident)

contr_TET1_filtered <- NormalizeData(object = contr_TET1_filtered)

contr_TET1_filtered <- FindVariableFeatures(object = contr_TET1_filtered)

contr_TET1_filtered <- ScaleData(object = contr_TET1_filtered)

contr_TET1_filtered <- RunPCA(object = contr_TET1_filtered)

contr_TET1_filtered <- FindNeighbors(object = contr_TET1_filtered, dims = 1:20, reduction = "pca")

contr_TET1_filtered <- FindClusters(object = contr_TET1_filtered, resolution = 0.4, cluster.name = "Cluster")

contr_TET1_filtered <- RunUMAP(object = contr_TET1_filtered, dims = 1:20, reduction = "pca", reduction.name = "UMAP")

DimPlot(contr_TET1_filtered, reduction = "UMAP", shuffle = TRUE)

sceT1 <- scDblFinder(GetAssayData(contr_TET1_filtered, slot = "counts"), clusters = Idents(contr_TET1_filtered))

contr_TET1_filtered$scDblFinder.class <- sceT1$scDblFinder.class

table(contr_TET1_filtered@meta.data$scDblFinder.class)

DimPlot(contr_TET1_filtered, reduction = "UMAP", shuffle = TRUE, group.by = "scDblFinder.class")

DimPlot(contr_TET1_filtered, reduction = "UMAP", shuffle = TRUE, split.by = "scDblFinder.class")

Idents(contr_TET1_filtered) <- contr_TET1_filtered@meta.data$scDblFinder.class

VlnPlot(contr_TET1_filtered, features = c("nFeature_RNA", "nCount_RNA"), ncol = 2)

contr_TET1_filtered <- subset(contr_TET1_filtered, subset = scDblFinder.class == "singlet")

DimPlot(contr_TET1_filtered, reduction = "UMAP", shuffle = TRUE, label = TRUE)

obj_hdf5_D2 = load10X("/media/dmitry/d/scRNAseq_TET/D2/outs/")

head(obj_hdf5_D2$soupProfile[order(obj_hdf5_D2$soupProfile$est, decreasing = TRUE), ], n = 20)

useToEst_D2 = estimateNonExpressingCells(obj_hdf5_D2, nonExpressedGeneList = nonExpressedGeneList)

obj_hdf5_D2 = calculateContaminationFraction(obj_hdf5_D2, nonExpressedGeneList, useToEst = useToEst_D2)

head(obj_hdf5_D2$metaData)

sceD2 = adjustCounts(obj_hdf5_D2)

contr_TET2 <- CreateSeuratObject(counts = sceD2, min.cells = 3, min.features = 200, project = "TET2")

contr_TET2$mitoPercent <- PercentageFeatureSet(contr_TET2, pattern='^mt-')

contr_TET2$riboPercent <- PercentageFeatureSet(contr_TET2, pattern='^Rp[sl]')

contr_TET2_filtered <- subset(contr_TET2, subset = nCount_RNA > 500 & nFeature_RNA > 500 & mitoPercent < 5 & riboPercent < 5)

table(contr_TET2_filtered$orig.ident)

contr_TET2_filtered <- NormalizeData(object = contr_TET2_filtered)

contr_TET2_filtered <- FindVariableFeatures(object = contr_TET2_filtered)

contr_TET2_filtered <- ScaleData(object = contr_TET2_filtered)

contr_TET2_filtered <- RunPCA(object = contr_TET2_filtered)

contr_TET2_filtered <- FindNeighbors(object = contr_TET2_filtered, dims = 1:20, reduction = "pca")

contr_TET2_filtered <- FindClusters(object = contr_TET2_filtered, resolution = 0.4, cluster.name = "Cluster")

contr_TET2_filtered <- RunUMAP(object = contr_TET2_filtered, dims = 1:20, reduction = "pca", reduction.name = "UMAP")

DimPlot(contr_TET2_filtered, reduction = "UMAP", shuffle = TRUE)

sceT2 <- scDblFinder(GetAssayData(contr_TET2_filtered, slot = "counts"), clusters = Idents(contr_TET2_filtered))

contr_TET2_filtered$scDblFinder.class <- sceT2$scDblFinder.class

table(contr_TET2_filtered@meta.data$scDblFinder.class)

DimPlot(contr_TET2_filtered, reduction = "UMAP", shuffle = TRUE, group.by = "scDblFinder.class")

DimPlot(contr_TET2_filtered, reduction = "UMAP", shuffle = TRUE, split.by = "scDblFinder.class")

Idents(contr_TET2_filtered) <- contr_TET2_filtered@meta.data$scDblFinder.class

VlnPlot(contr_TET2_filtered, features = c("nFeature_RNA", "nCount_RNA"), ncol = 2)

contr_TET2_filtered <- subset(contr_TET2_filtered, subset = scDblFinder.class == "singlet")

DimPlot(contr_TET2_filtered, reduction = "UMAP", shuffle = TRUE, label = TRUE)

merged_seurat <- merge(contr_TET1_filtered, y = contr_TET2_filtered, add.cell.ids = c("contr_TET1", "contr_TET2"), project = "integrated")

table(merged_seurat$orig.ident)

merged_seurat$sample <- rownames(merged_seurat@meta.data)

merged_seurat@meta.data <- separate(merged_seurat@meta.data, col = 'sample', into = c('Type', 'mouse', 'Barcode'), sep = '_')

merged_seurat <- NormalizeData(object = merged_seurat)

merged_seurat <- FindVariableFeatures(object = merged_seurat)

merged_seurat <- ScaleData(object = merged_seurat)

merged_seurat <- RunPCA(object = merged_seurat)

merged_seurat <- FindNeighbors(object = merged_seurat, dims = 1:20, reduction = "pca")

merged_seurat <- FindClusters(object = merged_seurat, resolution = 0.4, cluster.name = "unintegrCluster")

merged_seurat <- RunUMAP(object = merged_seurat, dims = 1:20, reduction = "pca", reduction.name = "unintegrUMAP")

CCA_integrated <- IntegrateLayers(object = merged_seurat, method = CCAIntegration, orig.reduction = "pca", new.reduction = "CCA.Integration", verbose =TRUE)

CCA_integrated[["RNA"]] <- JoinLayers(CCA_integrated[["RNA"]])

CCA_integrated <- FindNeighbors(object = CCA_integrated, dims = 1:20, reduction = "CCA.Integration")

CCA_integrated <- FindClusters(object = CCA_integrated, resolution = 0.4, cluster.name = "CCACluster")

CCA_integrated <- RunUMAP(object = CCA_integrated, dims = 1:20, reduction = "CCA.Integration", reduction.name = "CCAUMAP")

CCA_integrated <- RunTSNE(object = CCA_integrated, dims = 1:20, reduction = "CCA.Integration", reduction.name = "CCATSNE")

B) Analysis of data obtained for TET-deficient (Chx10TET) retinas

library(Seurat)

library(ggplot2)

library(gridExtra)

library(tidyr)

library(BPCells)

set.seed(123)

suppressPackageStartupMessages(library(scDblFinder))

library(SoupX)

nonExpressedGeneList = list(rods = c("Sag", "Pdc", "Rp1"), mg = c("Glul", "Rlbp1", "Slc1a3"))

obj_hdf5_C2 = load10X("/media/dmitry/d/scRNAseq_TET/C2/outs/")

head(obj_hdf5_C2$soupProfile[order(obj_hdf5_C2$soupProfile$est, decreasing = TRUE), ], n = 20)

useToEst_C2 = estimateNonExpressingCells(obj_hdf5_C2, nonExpressedGeneList = nonExpressedGeneList)

obj_hdf5_C2 = calculateContaminationFraction(obj_hdf5_C2, nonExpressedGeneList, useToEst = useToEst_C2)

head(obj_hdf5_C2$metaData)

sceC2 = adjustCounts(obj_hdf5_C2)

ko_Chx10TET1 <- CreateSeuratObject(counts = sceC2, min.cells = 3, min.features = 200, project = "Chx10TET1")

ko_Chx10TET1$mitoPercent <- PercentageFeatureSet(ko_Chx10TET1, pattern='^mt-')

ko_Chx10TET1$riboPercent <- PercentageFeatureSet(ko_Chx10TET1, pattern='^Rp[sl]')

ko_Chx10TET1_filtered <- subset(ko_Chx10TET1, subset = nCount_RNA > 500 & nFeature_RNA > 500 & mitoPercent < 5 & riboPercent < 5)

table(ko_Chx10TET1$orig.ident)

ko_Chx10TET1_filtered <- NormalizeData(object = ko_Chx10TET1_filtered)

ko_Chx10TET1_filtered <- FindVariableFeatures(object = ko_Chx10TET1_filtered)

ko_Chx10TET1_filtered <- ScaleData(object = ko_Chx10TET1_filtered)

ko_Chx10TET1_filtered <- RunPCA(object = ko_Chx10TET1_filtered)

ko_Chx10TET1_filtered <- FindNeighbors(object = ko_Chx10TET1_filtered, dims = 1:20, reduction = "pca")

ko_Chx10TET1_filtered <- FindClusters(object = ko_Chx10TET1_filtered, resolution = 0.4, cluster.name = "Cluster")

ko_Chx10TET1_filtered <- RunUMAP(object = ko_Chx10TET1_filtered, dims = 1:20, reduction = "pca", reduction.name = "UMAP")

DimPlot(ko_Chx10TET1_filtered, reduction = "UMAP", shuffle = TRUE)

sceCT1 <- scDblFinder(GetAssayData(ko_Chx10TET1_filtered, slot = "counts"), clusters = Idents(ko_Chx10TET1_filtered))

ko_Chx10TET1_filtered$scDblFinder.class <- sceCT1$scDblFinder.class

table(ko_Chx10TET1_filtered@meta.data$scDblFinder.class)

DimPlot(ko_Chx10TET1_filtered, reduction = "UMAP", shuffle = TRUE, group.by = "scDblFinder.class")

DimPlot(ko_Chx10TET1_filtered, reduction = "UMAP", shuffle = TRUE, split.by = "scDblFinder.class")

Idents(ko_Chx10TET1_filtered) <- ko_Chx10TET1_filtered@meta.data$scDblFinder.class

VlnPlot(ko_Chx10TET1_filtered, features = c("nFeature_RNA", "nCount_RNA"), ncol = 2)

ko_Chx10TET1_filtered <- subset(ko_Chx10TET1_filtered, subset = scDblFinder.class == "singlet")

DimPlot(ko_Chx10TET1_filtered, reduction = "UMAP", shuffle = TRUE, label = TRUE)

obj_hdf5_C3 = load10X("/media/dmitry/d/scRNAseq_TET/C3/outs/")

head(obj_hdf5_C3$soupProfile[order(obj_hdf5_C3$soupProfile$est, decreasing = TRUE), ], n = 20)

useToEst_C3 = estimateNonExpressingCells(obj_hdf5_C3, nonExpressedGeneList = nonExpressedGeneList)

obj_hdf5_C3 = calculateContaminationFraction(obj_hdf5_C3, nonExpressedGeneList, useToEst = useToEst_C3)

head(obj_hdf5_C3$metaData)

sceC3 = adjustCounts(obj_hdf5_C3)

ko_Chx10TET2 <- CreateSeuratObject(counts = sceC3, min.cells = 3, min.features = 200, project = "Chx10TET2")

ko_Chx10TET2$mitoPercent <- PercentageFeatureSet(ko_Chx10TET2, pattern='^mt-')

ko_Chx10TET2$riboPercent <- PercentageFeatureSet(ko_Chx10TET2, pattern='^Rp[sl]')

ko_Chx10TET2_filtered <- subset(ko_Chx10TET2, subset = nCount_RNA > 500 & nFeature_RNA > 500 & mitoPercent < 5 & riboPercent < 5)

table(ko_Chx10TET2$orig.ident)

ko_Chx10TET2_filtered <- NormalizeData(object = ko_Chx10TET2_filtered)

ko_Chx10TET2_filtered <- FindVariableFeatures(object = ko_Chx10TET2_filtered)

ko_Chx10TET2_filtered <- ScaleData(object = ko_Chx10TET2_filtered)

ko_Chx10TET2_filtered <- RunPCA(object = ko_Chx10TET2_filtered)

ko_Chx10TET2_filtered <- FindNeighbors(object = ko_Chx10TET2_filtered, dims = 1:20, reduction = "pca")

ko_Chx10TET2_filtered <- FindClusters(object = ko_Chx10TET2_filtered, resolution = 0.4, cluster.name = "Cluster")

ko_Chx10TET2_filtered <- RunUMAP(object = ko_Chx10TET2_filtered, dims = 1:20, reduction = "pca", reduction.name = "UMAP")

DimPlot(ko_Chx10TET2_filtered, reduction = "UMAP", shuffle = TRUE)

sceCT2 <- scDblFinder(GetAssayData(ko_Chx10TET2_filtered, slot = "counts"), clusters = Idents(ko_Chx10TET2_filtered))

ko_Chx10TET2_filtered$scDblFinder.class <- sceCT2$scDblFinder.class

table(ko_Chx10TET2_filtered@meta.data$scDblFinder.class)

DimPlot(ko_Chx10TET2_filtered, reduction = "UMAP", shuffle = TRUE, group.by = "scDblFinder.class")

DimPlot(ko_Chx10TET2_filtered, reduction = "UMAP", shuffle = TRUE, split.by = "scDblFinder.class")

Idents(ko_Chx10TET2_filtered) <- ko_Chx10TET2_filtered@meta.data$scDblFinder.class

VlnPlot(ko_Chx10TET2_filtered, features = c("nFeature_RNA", "nCount_RNA"), ncol = 2)

ko_Chx10TET2_filtered <- subset(ko_Chx10TET2_filtered, subset = scDblFinder.class == "singlet")

DimPlot(ko_Chx10TET2_filtered, reduction = "UMAP", shuffle = TRUE, label = TRUE)

merged_seurat <- merge(ko_Chx10TET1_filtered, y = c(ko_Chx10TET2_filtered), add.cell.ids = c("ko_Chx10TET1", "ko_Chx10TET2"), project = "integrated")

table(merged_seurat$orig.ident)

merged_seurat$sample <- rownames(merged_seurat@meta.data)

merged_seurat@meta.data <- separate(merged_seurat@meta.data, col = 'sample', into = c('Type', 'mouse', 'Barcode'), sep = '_')

merged_seurat <- NormalizeData(object = merged_seurat)

merged_seurat <- FindVariableFeatures(object = merged_seurat)

merged_seurat <- ScaleData(object = merged_seurat)

merged_seurat <- RunPCA(object = merged_seurat)

merged_seurat <- FindNeighbors(object = merged_seurat, dims = 1:20, reduction = "pca")

merged_seurat <- FindClusters(object = merged_seurat, resolution = 0.4, cluster.name = "unintegrCluster")

merged_seurat <- RunUMAP(object = merged_seurat, dims = 1:20, reduction = "pca", reduction.name = "unintegrUMAP")

CCA_integrated <- IntegrateLayers(object = merged_seurat, method = CCAIntegration, orig.reduction = "pca", new.reduction = "CCA.Integration", verbose =TRUE)

CCA_integrated[["RNA"]] <- JoinLayers(CCA_integrated[["RNA"]])

CCA_integrated <- FindNeighbors(object = CCA_integrated, dims = 1:20, reduction = "CCA.Integration")

CCA_integrated <- FindClusters(object = CCA_integrated, resolution = 0.4, cluster.name = "CCACluster")

CCA_integrated <- RunUMAP(object = CCA_integrated, dims = 1:20, reduction = "CCA.Integration", reduction.name = "CCAUMAP")

CCA_integrated <- RunTSNE(object = CCA_integrated, dims = 1:20, reduction = "CCA.Integration", reduction.name = "CCATSNE")

C) TET and Chx10TET data integration

library(Seurat)

library(ggplot2)

library(gridExtra)

library(tidyr)

library(BPCells)

set.seed(123)

suppressPackageStartupMessages(library(scDblFinder))

library(SoupX)

nonExpressedGeneList = list(rods = c("Sag", "Pdc", "Rp1"), mg = c("Glul", "Rlbp1", "Slc1a3"))

obj_hdf5_D1 = load10X("/media/dmitry/d/scRNAseq_TET/D1/outs/")

head(obj_hdf5_D1$soupProfile[order(obj_hdf5_D1$soupProfile$est, decreasing = TRUE), ], n = 20)

useToEst_D1 = estimateNonExpressingCells(obj_hdf5_D1, nonExpressedGeneList = nonExpressedGeneList)

obj_hdf5_D1 = calculateContaminationFraction(obj_hdf5_D1, nonExpressedGeneList, useToEst = useToEst_D1)

head(obj_hdf5_D1$metaData)

sceD1 = adjustCounts(obj_hdf5_D1)

contr_TET1 <- CreateSeuratObject(counts = sceD1, min.cells = 3, min.features = 200, project = "TET1")

contr_TET1$mitoPercent <- PercentageFeatureSet(contr_TET1, pattern='^mt-')

contr_TET1$riboPercent <- PercentageFeatureSet(contr_TET1, pattern='^Rp[sl]')

contr_TET1_filtered <- subset(contr_TET1, subset = nCount_RNA > 500 & nFeature_RNA > 500 & mitoPercent < 5 & riboPercent < 5)

table(contr_TET1_filtered$orig.ident)

contr_TET1_filtered <- NormalizeData(object = contr_TET1_filtered)

contr_TET1_filtered <- FindVariableFeatures(object = contr_TET1_filtered)

contr_TET1_filtered <- ScaleData(object = contr_TET1_filtered)

contr_TET1_filtered <- RunPCA(object = contr_TET1_filtered)

contr_TET1_filtered <- FindNeighbors(object = contr_TET1_filtered, dims = 1:20, reduction = "pca")

contr_TET1_filtered <- FindClusters(object = contr_TET1_filtered, resolution = 0.4, cluster.name = "Cluster")

contr_TET1_filtered <- RunUMAP(object = contr_TET1_filtered, dims = 1:20, reduction = "pca", reduction.name = "UMAP")

DimPlot(contr_TET1_filtered, reduction = "UMAP", shuffle = TRUE)

sceT1 <- scDblFinder(GetAssayData(contr_TET1_filtered, slot = "counts"), clusters = Idents(contr_TET1_filtered))

contr_TET1_filtered$scDblFinder.class <- sceT1$scDblFinder.class

table(contr_TET1_filtered@meta.data$scDblFinder.class)

DimPlot(contr_TET1_filtered, reduction = "UMAP", shuffle = TRUE, group.by = "scDblFinder.class")

DimPlot(contr_TET1_filtered, reduction = "UMAP", shuffle = TRUE, split.by = "scDblFinder.class")

Idents(contr_TET1_filtered) <- contr_TET1_filtered@meta.data$scDblFinder.class

VlnPlot(contr_TET1_filtered, features = c("nFeature_RNA", "nCount_RNA"), ncol = 2)

contr_TET1_filtered <- subset(contr_TET1_filtered, subset = scDblFinder.class == "singlet")

DimPlot(contr_TET1_filtered, reduction = "UMAP", shuffle = TRUE, label = TRUE)

obj_hdf5_D2 = load10X("/media/dmitry/d/scRNAseq_TET/D2/outs/")

head(obj_hdf5_D2$soupProfile[order(obj_hdf5_D2$soupProfile$est, decreasing = TRUE), ], n = 20)

useToEst_D2 = estimateNonExpressingCells(obj_hdf5_D2, nonExpressedGeneList = nonExpressedGeneList)

obj_hdf5_D2 = calculateContaminationFraction(obj_hdf5_D2, nonExpressedGeneList, useToEst = useToEst_D2)

head(obj_hdf5_D2$metaData)

sceD2 = adjustCounts(obj_hdf5_D2)

contr_TET2 <- CreateSeuratObject(counts = sceD2, min.cells = 3, min.features = 200, project = "TET2")

contr_TET2$mitoPercent <- PercentageFeatureSet(contr_TET2, pattern='^mt-')

contr_TET2$riboPercent <- PercentageFeatureSet(contr_TET2, pattern='^Rp[sl]')

contr_TET2_filtered <- subset(contr_TET2, subset = nCount_RNA > 500 & nFeature_RNA > 500 & mitoPercent < 5 & riboPercent < 5)

table(contr_TET2_filtered$orig.ident)

contr_TET2_filtered <- NormalizeData(object = contr_TET2_filtered)

contr_TET2_filtered <- FindVariableFeatures(object = contr_TET2_filtered)

contr_TET2_filtered <- ScaleData(object = contr_TET2_filtered)

contr_TET2_filtered <- RunPCA(object = contr_TET2_filtered)

contr_TET2_filtered <- FindNeighbors(object = contr_TET2_filtered, dims = 1:20, reduction = "pca")

contr_TET2_filtered <- FindClusters(object = contr_TET2_filtered, resolution = 0.4, cluster.name = "Cluster")

contr_TET2_filtered <- RunUMAP(object = contr_TET2_filtered, dims = 1:20, reduction = "pca", reduction.name = "UMAP")

DimPlot(contr_TET2_filtered, reduction = "UMAP", shuffle = TRUE)

sceT2 <- scDblFinder(GetAssayData(contr_TET2_filtered, slot = "counts"), clusters = Idents(contr_TET2_filtered))

contr_TET2_filtered$scDblFinder.class <- sceT2$scDblFinder.class

table(contr_TET2_filtered@meta.data$scDblFinder.class)

DimPlot(contr_TET2_filtered, reduction = "UMAP", shuffle = TRUE, group.by = "scDblFinder.class")

DimPlot(contr_TET2_filtered, reduction = "UMAP", shuffle = TRUE, split.by = "scDblFinder.class")

Idents(contr_TET2_filtered) <- contr_TET2_filtered@meta.data$scDblFinder.class

VlnPlot(contr_TET2_filtered, features = c("nFeature_RNA", "nCount_RNA"), ncol = 2)

contr_TET2_filtered <- subset(contr_TET2_filtered, subset = scDblFinder.class == "singlet")

DimPlot(contr_TET2_filtered, reduction = "UMAP", shuffle = TRUE, label = TRUE)

obj_hdf5_C2 = load10X("/media/dmitry/d/scRNAseq_TET/C2/outs/")

head(obj_hdf5_C2$soupProfile[order(obj_hdf5_C2$soupProfile$est, decreasing = TRUE), ], n = 20)

useToEst_C2 = estimateNonExpressingCells(obj_hdf5_C2, nonExpressedGeneList = nonExpressedGeneList)

obj_hdf5_C2 = calculateContaminationFraction(obj_hdf5_C2, nonExpressedGeneList, useToEst = useToEst_C2)

head(obj_hdf5_C2$metaData)

sceC2 = adjustCounts(obj_hdf5_C2)

ko_Chx10TET1 <- CreateSeuratObject(counts = sceC2, min.cells = 3, min.features = 200, project = "Chx10TET1")

ko_Chx10TET1$mitoPercent <- PercentageFeatureSet(ko_Chx10TET1, pattern='^mt-')

ko_Chx10TET1$riboPercent <- PercentageFeatureSet(ko_Chx10TET1, pattern='^Rp[sl]')

ko_Chx10TET1_filtered <- subset(ko_Chx10TET1, subset = nCount_RNA > 500 & nFeature_RNA > 500 & mitoPercent < 5 & riboPercent < 5)

table(ko_Chx10TET1$orig.ident)

ko_Chx10TET1_filtered <- NormalizeData(object = ko_Chx10TET1_filtered)

ko_Chx10TET1_filtered <- FindVariableFeatures(object = ko_Chx10TET1_filtered)

ko_Chx10TET1_filtered <- ScaleData(object = ko_Chx10TET1_filtered)

ko_Chx10TET1_filtered <- RunPCA(object = ko_Chx10TET1_filtered)

ko_Chx10TET1_filtered <- FindNeighbors(object = ko_Chx10TET1_filtered, dims = 1:20, reduction = "pca")

ko_Chx10TET1_filtered <- FindClusters(object = ko_Chx10TET1_filtered, resolution = 0.4, cluster.name = "Cluster")

ko_Chx10TET1_filtered <- RunUMAP(object = ko_Chx10TET1_filtered, dims = 1:20, reduction = "pca", reduction.name = "UMAP")

DimPlot(ko_Chx10TET1_filtered, reduction = "UMAP", shuffle = TRUE)

sceCT1 <- scDblFinder(GetAssayData(ko_Chx10TET1_filtered, slot = "counts"), clusters = Idents(ko_Chx10TET1_filtered))

ko_Chx10TET1_filtered$scDblFinder.class <- sceCT1$scDblFinder.class

table(ko_Chx10TET1_filtered@meta.data$scDblFinder.class)

DimPlot(ko_Chx10TET1_filtered, reduction = "UMAP", shuffle = TRUE, group.by = "scDblFinder.class")

DimPlot(ko_Chx10TET1_filtered, reduction = "UMAP", shuffle = TRUE, split.by = "scDblFinder.class")

Idents(ko_Chx10TET1_filtered) <- ko_Chx10TET1_filtered@meta.data$scDblFinder.class

VlnPlot(ko_Chx10TET1_filtered, features = c("nFeature_RNA", "nCount_RNA"), ncol = 2)

ko_Chx10TET1_filtered <- subset(ko_Chx10TET1_filtered, subset = scDblFinder.class == "singlet")

DimPlot(ko_Chx10TET1_filtered, reduction = "UMAP", shuffle = TRUE, label = TRUE)

obj_hdf5_C3 = load10X("/media/dmitry/d/scRNAseq_TET/C3/outs/")

head(obj_hdf5_C3$soupProfile[order(obj_hdf5_C3$soupProfile$est, decreasing = TRUE), ], n = 20)

useToEst_C3 = estimateNonExpressingCells(obj_hdf5_C3, nonExpressedGeneList = nonExpressedGeneList)

obj_hdf5_C3 = calculateContaminationFraction(obj_hdf5_C3, nonExpressedGeneList, useToEst = useToEst_C3)

head(obj_hdf5_C3$metaData)

sceC3 = adjustCounts(obj_hdf5_C3)

ko_Chx10TET2 <- CreateSeuratObject(counts = sceC3, min.cells = 3, min.features = 200, project = "Chx10TET2")

ko_Chx10TET2$mitoPercent <- PercentageFeatureSet(ko_Chx10TET2, pattern='^mt-')

ko_Chx10TET2$riboPercent <- PercentageFeatureSet(ko_Chx10TET2, pattern='^Rp[sl]')

ko_Chx10TET2_filtered <- subset(ko_Chx10TET2, subset = nCount_RNA > 500 & nFeature_RNA > 500 & mitoPercent < 5 & riboPercent < 5)

table(ko_Chx10TET2$orig.ident)

ko_Chx10TET2_filtered <- NormalizeData(object = ko_Chx10TET2_filtered)

ko_Chx10TET2_filtered <- FindVariableFeatures(object = ko_Chx10TET2_filtered)

ko_Chx10TET2_filtered <- ScaleData(object = ko_Chx10TET2_filtered)

ko_Chx10TET2_filtered <- RunPCA(object = ko_Chx10TET2_filtered)

ko_Chx10TET2_filtered <- FindNeighbors(object = ko_Chx10TET2_filtered, dims = 1:20, reduction = "pca")

ko_Chx10TET2_filtered <- FindClusters(object = ko_Chx10TET2_filtered, resolution = 0.4, cluster.name = "Cluster")

ko_Chx10TET2_filtered <- RunUMAP(object = ko_Chx10TET2_filtered, dims = 1:20, reduction = "pca", reduction.name = "UMAP")

DimPlot(ko_Chx10TET2_filtered, reduction = "UMAP", shuffle = TRUE)

sceCT2 <- scDblFinder(GetAssayData(ko_Chx10TET2_filtered, slot = "counts"), clusters = Idents(ko_Chx10TET2_filtered))

ko_Chx10TET2_filtered$scDblFinder.class <- sceCT2$scDblFinder.class

table(ko_Chx10TET2_filtered@meta.data$scDblFinder.class)

DimPlot(ko_Chx10TET2_filtered, reduction = "UMAP", shuffle = TRUE, group.by = "scDblFinder.class")

DimPlot(ko_Chx10TET2_filtered, reduction = "UMAP", shuffle = TRUE, split.by = "scDblFinder.class")

Idents(ko_Chx10TET2_filtered) <- ko_Chx10TET2_filtered@meta.data$scDblFinder.class

VlnPlot(ko_Chx10TET2_filtered, features = c("nFeature_RNA", "nCount_RNA"), ncol = 2)

ko_Chx10TET2_filtered <- subset(ko_Chx10TET2_filtered, subset = scDblFinder.class == "singlet")

DimPlot(ko_Chx10TET2_filtered, reduction = "UMAP", shuffle = TRUE, label = TRUE)

merged_seurat <- merge(contr_TET1_filtered, y = c(contr_TET2_filtered, ko_Chx10TET1_filtered, ko_Chx10TET2_filtered), add.cell.ids = c("contr_TET1", "contr_TET2", "ko_Chx10TET1", "ko_Chx10TET2"), project = "integrated")

table(merged_seurat$orig.ident)

merged_seurat$sample <- rownames(merged_seurat@meta.data)

merged_seurat@meta.data <- separate(merged_seurat@meta.data, col = 'sample', into = c('Type', 'mouse', 'Barcode'), sep = '_')

merged_seurat <- NormalizeData(object = merged_seurat)

merged_seurat <- FindVariableFeatures(object = merged_seurat)

merged_seurat <- ScaleData(object = merged_seurat)

merged_seurat <- RunPCA(object = merged_seurat)

merged_seurat <- FindNeighbors(object = merged_seurat, dims = 1:20, reduction = "pca")

merged_seurat <- FindClusters(object = merged_seurat, resolution = 0.4, cluster.name = "unintegrCluster")

merged_seurat <- RunUMAP(object = merged_seurat, dims = 1:20, reduction = "pca", reduction.name = "unintegrUMAP")

CCA_integrated <- IntegrateLayers(object = merged_seurat, method = CCAIntegration, orig.reduction = "pca", new.reduction = "CCA.Integration", verbose =TRUE)

CCA_integrated[["RNA"]] <- JoinLayers(CCA_integrated[["RNA"]])

CCA_integrated <- FindNeighbors(object = CCA_integrated, dims = 1:20, reduction = "CCA.Integration")

CCA_integrated <- FindClusters(object = CCA_integrated, resolution = 0.4, cluster.name = "CCACluster")

CCA_integrated <- RunUMAP(object = CCA_integrated, dims = 1:20, reduction = "CCA.Integration", reduction.name = "CCAUMAP")

CCA_integrated <- RunTSNE(object = CCA_integrated, dims = 1:20, reduction = "CCA.Integration", reduction.name = "CCATSNE")
